# Origin of chaperone dependence and assembly complexity in Rubisco’s biogenesis

**DOI:** 10.1101/2025.09.22.677769

**Authors:** Jediael Z. Y. Ng, Dennis Wiens, Luca Schulz, Alisa Hergenröder, Andreas M. Küffner, Christin Geil, Tobias J. Erb, Georg K. A. Hochberg

**Author notes:** Max Planck Foundation Research Group, Max Planck Institute for Multidisciplinary Sciences; Göttingen, 37077, Germany (Andreas M. Küffner).

## Abstract

Molecular chaperones assist with the folding and assembly of protein clients. Consequently, they are essential to diverse cellular functions. In most aerobic photosynthetic organisms such as β-cyanobacteria and all plants, the key CO_2_-fixing enzyme Rubisco strictly relies on dedicated chaperones for its assembly. Why these chaperones were recruited is unclear, as they are absent and not required in other prokaryotic autotrophs. By retracing the evolution of Rubisco-chaperone interactions, we find that Rubisco’s dependence on chaperones evolved neutrally and not by adaptive optimization, and that this may be reversed without compromising catalytic function. Our work shows how chaperones are broad modifiers of their clients’ sequence spaces that can become totally essential through non-adaptive processes.

## Introduction

Proteins must fold and assemble correctly in a crowded, chaotic cellular environment. To accomplish this, many proteins depend entirely on molecular chaperones, which provide transient assistance to their clients during folding or assembly (*1*). How proteins come to rely on chaperones is not well understood. Chaperone requirements can vary within protein families, suggesting that the need for chaperone assistance during folding or assembly is not always an unavoidable feature of a protein’s fold, but can be acquired or lost in the course of history. Evolutionary theory suggests that the acquisition of chaperone dependence (i.e. the inability to fold or assemble without chaperones) may be indirectly beneficial to proteins, through so-called adaptive buffering: Chaperones may allow the fixation of mutations in their clients that are functionally beneficial, but are also destabilizing and intolerable without a chaperone (*2– 6*). In this theory, improved function of the client comes at the cost of total dependence on a chaperone. An alternative theory is that proteins come to depend on chaperones through the action of a neutral evolutionary ratchet (*7–9*): this can happen through mutations that are very detrimental to function on their own, which become neutral in the presence of chaperones (*10*). If such mutations fix through the action of neutral genetic drift, clients can become completely dependent on chaperones without gaining any improvement in their function in return (*11*).

One example of a protein family known to have become dependent on several new chaperones in its history is ribulose-1,5-bisphosphate carboxylase/oxygenase (Rubisco) (Fig. 1A) (*12–14*), the enzyme responsible for CO_2_ fixation in oxygenic photosynthesis and probably the most abundant protein on Earth (*15–17*). The most ancient versions of Form I Rubisco are first folded in the chaperonin cage of the folding chaperone GroEL/ES and then self-assemble without further assistance into either homo-oligomers consisting of eight catalytic large (RbcL), or hetero-oligomers incorporating eight non-catalytic small subunits (RbcS) (*18–21*). The latter then became dependent on additional chaperones that help Rubisco’s large subunit assemble into complexes after it emerges from the chaperonin cage (Fig. 1A). Some β-cyanobacterial Rubiscos (1B clade) strictly require the action of either of two dedicated chaperones - Raf1 or RbcX - for the assembly of functional Rubisco complexes (*22–25*). Another increase in assembly complexity then followed with the addition of assembly chaperones Raf2 and BSD2 on top of Raf1 and RbcX in plants (Fig. 1A) (*26–29*).

**Figure 1.**
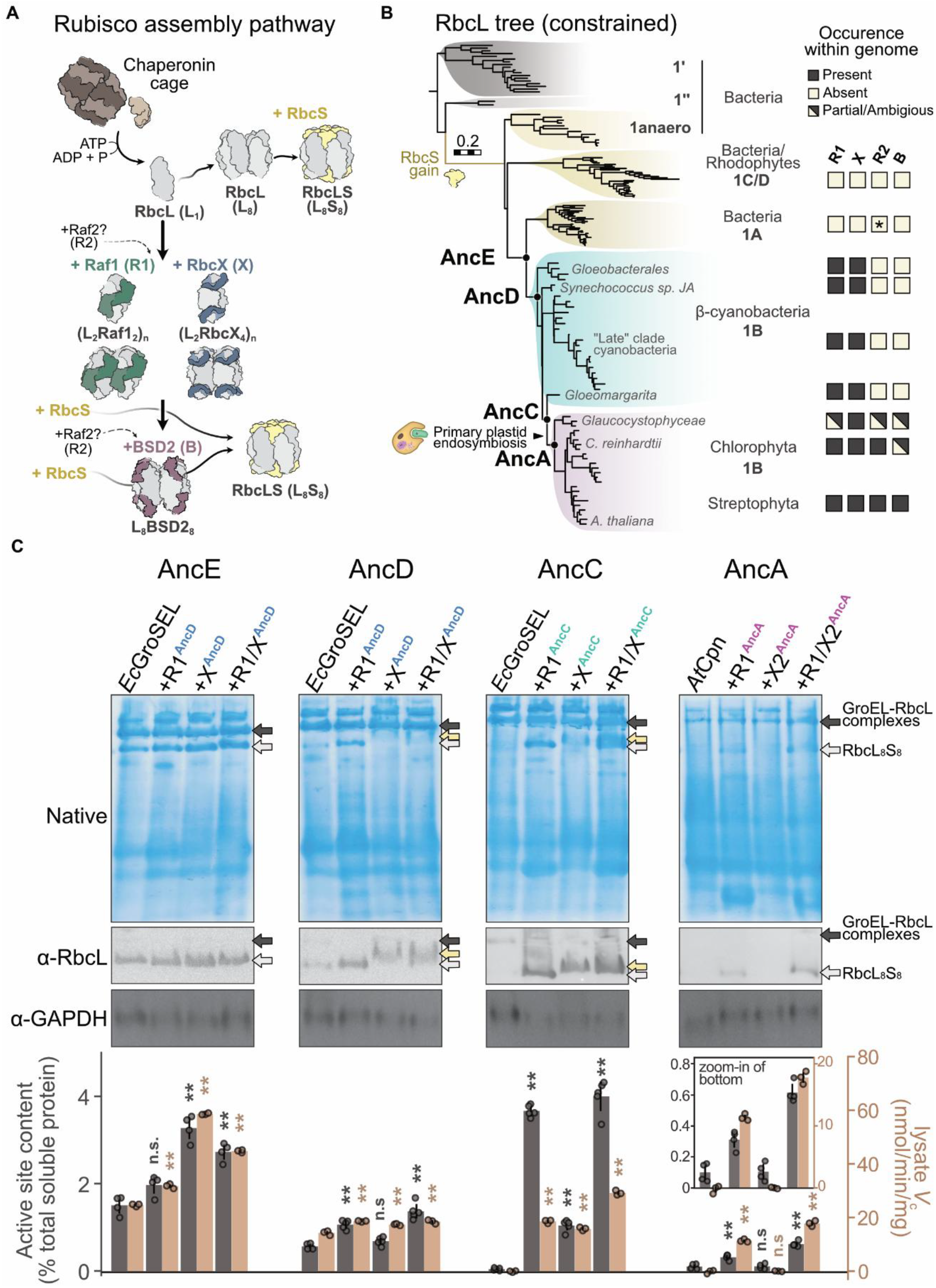
Evolution of Rubisco and assembly chaperones. **(A)** Biogenesis pathway in Form I Rubiscos. Early branching Form I Rubiscos may co-assemble directly with the small subunit (RbcS). Reliance on assembly chaperones Raf1 (R1) and RbcX (X) before RbcS addition first arose in β-cyanobacteria. Requirement of additional chaperones factors Raf2 (R2) and BSD2 (B) emerged later within green plastids. **(B)** Constrained Maximum Likelihood phylogeny of RbcL based on known species relationships. Ancestral Rubiscos at indicated nodes were resurrected and characterized. Clades according to their biogenesis requirements and final assemblies are coloured as per (A), with their designated Form 1 type and host organisms indicated on the right. Right column panel: The presence or absence of chaperones within the genomes of extant organisms within corresponding clades shown to the left in the tree. A putative Raf2-like homolog of a similar fold is found in α-Cyanobacteria as indicated by the asterisk. Partial or ambiguous classifications were assigned due to poor genome coverage or hits with low sequence identities. **(C)** Native PAGE and western blot analysis of ancestral Rubisco-chaperone co-expression combinations. White arrows indicate L_8_S_8_ assembly bands, black arrows indicate chaperonin-bound LSU complexes, and yellow arrows for plausible chaperone-bound LSU complexes. Bottom row left axis/grey bars: The amount of Rubisco active sites shown as percentages relative to total soluble protein obtained from lysate extracts of co-expressed combinations in the columns above. Error bars represent SDs. Significance as tested by one-way ANOVA (ANalysis Of VAriance) with post-hoc Tukey HSD (Honestly Significant Difference) test. *n*.*s*., nonsignificant; \*\**P* < 0.01. Data are from at least three technical replicates as indicated by coloured circles. Bottom row right axis: Rubisco activities measured in soluble lysate extracts from expression conditions indicated above at fixed CO_2_ (~200 µM) concentrations.

How and why these chaperones became essential is unclear, but adaptive buffering is a plausible mechanism: Rubisco’s history is plagued by a problematic relationship with oxygen, an undesired side substrate that results in the production of a toxic product. Over the course of its history, Rubisco has adapted to rising atmospheric oxygen concentrations by becoming more specific for CO_2_ over O_2_ (*30, 31*). Rubisco’s evolution is notoriously slow and potentially constrained by trade-offs between its catalytic speed and specificity for CO_2_ (*32–36*). It thus seems plausible that Rubisco may have become dependent on additional chaperones because they unlocked catalytically beneficial substitutions that would otherwise destabilize the enzyme. This theory, however, has remained untested, because both adaptive buffering and neutral entrenchment would result in the same indistinguishable phenotype when chaperones are removed in organisms: a complete loss of functional Rubisco assembly.

Here we recapitulate the evolution of Rubisco’s dependence on assembly chaperones Raf1 and RbcX using ancestral sequence reconstruction (*9, 37*). Through the reconstruction of a billion-year-old assembly pathway, we reveal that increased complexity in Rubisco’s biogenesis became essential via a neutral evolutionary ratchet. These findings also illuminate the historical trajectory transversed by the biogeochemically important Form I Rubiscos: Our study suggests that modern Rubiscos could be relieved from constraints imposed on them by chaperones, which may ultimately open up new strategies for engineering improved CO_2_ fixation in plants.

### Resurrection of Rubisco’s chaperone network

To investigate how Rubisco-chaperone interactions evolved, we began by inferring the phylogeny of Rubisco’s large subunit (LSU) (Fig. 1B, S2A, see methods). We then mapped onto it the occurrence of assembly chaperones within the genomes of extant species. Chaperones Raf1 and RbcX first appeared within β-cyanobacteria. To retrace when assembly chaperones first became essential, we inferred sequences of ancestral Rubiscos that existed before and after the occurrence of chaperones Raf1 and RbcX (Fig. 1B). The phylogeny of each chaperone was also inferred with matching topological constraints as our RbcL tree to obtain synchronized ancestral nodes (Fig. S1). This enabled us to reconstruct the ancient chaperone network that was likely to have co-existed with our Rubisco ancestors within the same ancestral organism as closely as possible (see Methods). This minimizes the potential for artefacts and incompatibility in Rubisco-chaperone pairings, which is a known obstacle to productive assembly (*38– 41*).

To determine when Rubisco started to rely on chaperone interactions for assembly, we resurrected 4 ancestors (Fig. 1B; AncE, D, C, A; AncB was insoluble, see Supplementary text). Going from the most ancient to the most recent reconstructions: AncE is the last common ancestor (LCA) of Form IA and Form IB Rubisco and would have existed before chaperones appeared in the genomes of β-cyanobacteria. AncD is the LCA of all Form IB Rubiscos, which would have existed in an ancestral β-cyanobacterium that already contained Raf1 and RbcX. AncC corresponds to the last node that unites cyanobacterial Rubiscos and their plastidal descendants, and therefore existed in an ancestral cyanobacterium immediately prior to the endosymbiosis event that birthed chloroplasts. Lastly, AncA corresponds to the LCA of all green algal and plant Rubiscos.

We next characterized the assembly characteristics of our ancestors by co-expressing them with or without their corresponding assembly chaperones within Escherichia coli (E. coli) and examining for assembled Rubisco complexes with native PAGE and immunoblot analysis (Fig. 1C). We also quantified the amount of functional active sites using the radiolabeled transition state analogue 2-^14^C-carboxy-D-arabinitol-1,5-bisphosphate (^14^C-CABP), and by assaying for Rubisco activity directly in lysates (Fig. 1C, bottom row). Our oldest ancestor AncE could assemble into functional complex without assembly chaperones (Fig. 1B), consistent with the fact that it predates the emergence of Raf1 and RbcX. Additionally, co-production of assembly chaperones Raf1 or RbcX boosted AncE’s assembly yields. Thus, the initial recruitment of these chaperones was likely beneficial because they probably would have increased the fidelity of Rubisco’s assembly pathway. It is important to note that this initially beneficial effect of adding chaperones on its own does not explain why Rubisco later becomes completely unable to assemble without them.

Our next ancestor, AncD, behaved similarly to its predecessor and could still assemble in absence of chaperones (Fig. 1C). Adding Raf1 also bolstered assembly yields, though RbcX no longer did so. Likewise, we saw the same capacity for self-assembly in extant Form I Rubiscos from Synechococcus sp. JA (SynJA) and Gloeobacter violaceus (*Gv*) that are close descendants of AncD (Fig. S2B) (*42*). In contrast, AncC could no longer assemble on its own and strictly required assistance from Raf1 for its assembly (Fig. 1C). This is also consistent with Rubisco from the cyanobacterium Gloeomargarita lithophora (*Gl*) (Fig. S2B), which descends from AncC and represents the closest cyanobacterial relative of plastidal Rubiscos (*43*). Our latest ancestor AncA remained totally dependent on assembly chaperones. Overall, our data show that assembly chaperones were not essential for assembly immediately after they first appeared, and that Rubisco became dependent on assembly chaperones within cyanobacteria sometime between AncD and AncC. This dependence, and a dominant role for Raf1, was then inherited by plastidal Rubiscos.

We next asked if the evolution of chaperone dependence coincided with improved enzyme function. To do so, we purified our ancestors and measured carboxylation kinetics with ribulose-1,5-bisphophate (RuBP) and specificity for CO_2_ over O_2_ (S_C/O_) (Table S1). From prokaryotic Form 1A/B (AncE) to the plastidal type Form 1B Rubisco (AncA), the Michaelis constant for CO_2_ (K_m_^*CO2*^) decreased 5.7-fold (Fig. 2B). This is coupled with somewhat fluctuating maximal carboxylation rates (*k*_cat_^c^) between 1.3 s^−1^ to 2.5 s^−1^ (Fig. 2A), resulting in a ~7.5-fold increase in overall carboxylation efficiencies over this interval (Fig. 2C). Likewise, a 1.6-fold improvement in specificity occurs between AncE to AncA (Fig. 2D). Intriguingly, the most significant improvements occurred between AncD and AncC, coinciding with the same interval over which chaperone dependence evolved. The temporal overlap between improved enzyme function and dependence thus hints that adaptive buffering by chaperones could have occurred.

**Figure 2.**
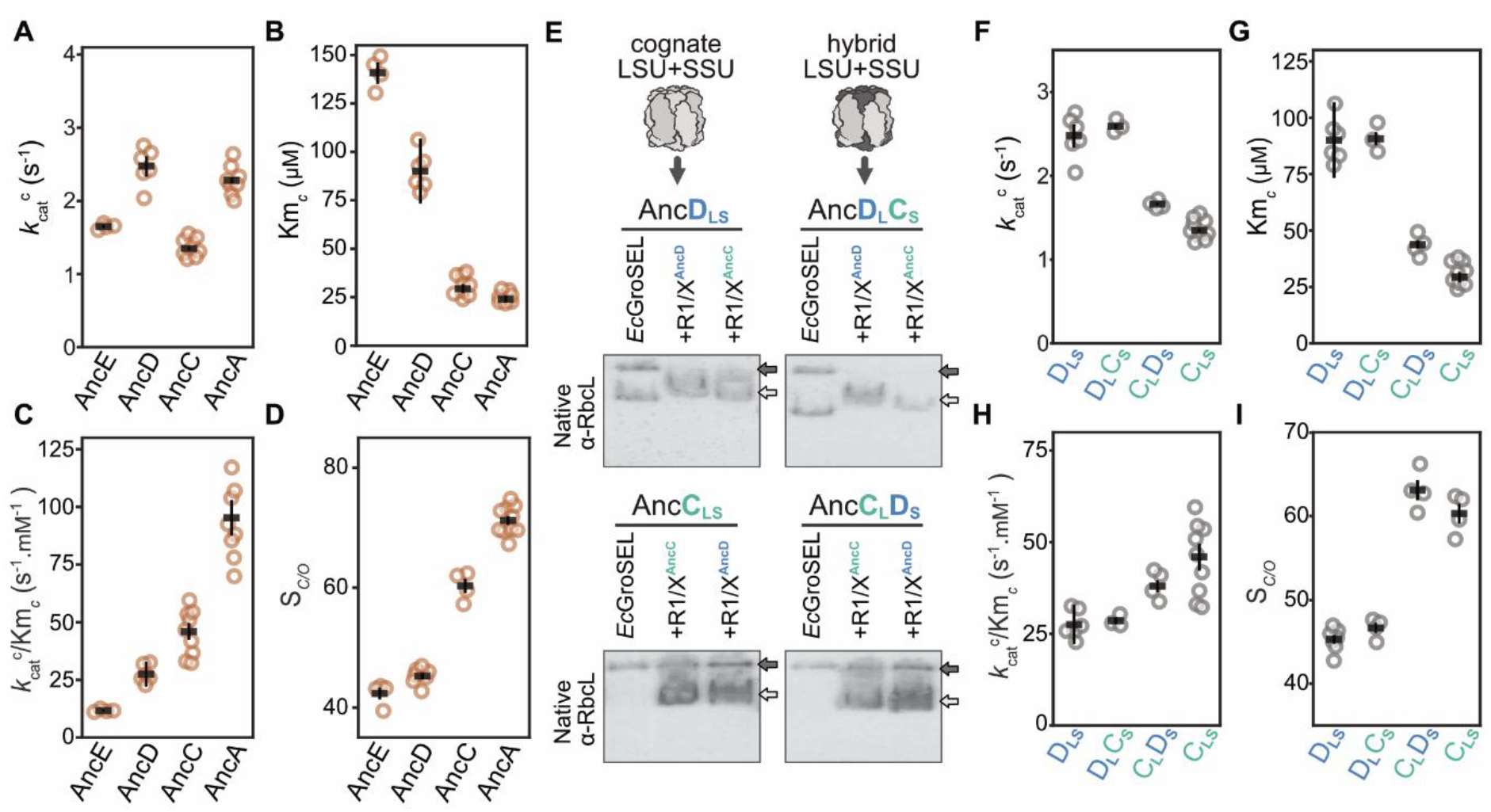
Changes in the LSU determine catalytic properties and assembly requirements. **(A-D)** Kinetic properties of ancestral Rubiscos. **(A)** Carboxylation rates, **(B)** Michaelis constant for CO_2_, **(C)** carboxylation efficiency, and **(D)** substrate specificity. **(E)** Native blot analysis of wild type and hybrid ancestral Rubisco assembly with matching or non-matching chaperone pairs. Neither the SSU nor chaperone identities influence the requirement for assembly chaperones. **(F-I)** Comparisons of kinetic parameters exhibited by hybrid AncD/C Rubiscos against wild-type enzymes. LSU identities are observed to dictate kinetic characteristics of hybrid enzymes. For all kinetic data, catalytic parameters extracted from individual Michaelis-Menten fits of technical replicates are shown as hollow circles, with globally fitted parameters denoted as a horizontal bar. Error bars represent SEs of the global fit. Values of global fits and replicate numbers are further indicated in table. S1 and kinetic curve fits shown in fig. S17.

These results were verified against two kinds of uncertainty. First, we resurrected sequences of less likely ancestral Rubiscos from our constrained phylogeny (Fig. S2C), in which the maximum a posteriori amino acid was replaced with the next most plausible amino acid whenever the second state had a posterior probability of at least 0.2. In the second, we tested ancestors inferred using an unconstrained ML RbcL phylogeny (Fig. S2D-E), which would require many horizontal gene transfer (HGT) events to explain the branching order of RbcL within cyanobacteria (Fig. S2A). Our inferences remained robust in both cases: Rubisco is already dependent on chaperones at the node that unites plastid and cyanobacterial Rubiscos. Therefore, we proceeded with ancestors inferred from the constrained phylogeny for the rest of this work.

### Genetic basis of dependence on chaperones Raf1/RbcX

Next, we sought to assess if the evolution of chaperone dependence is causally linked to improved Rubisco function. To do this, we first needed to determine which changes in which proteins were responsible for the dependence on chaperones and improved catalysis. We therefore constructed hybrid ancestors containing swapped large and small subunits of AncD and AncC (AncD_L_C_S_, AncC_L_D_S_). Both hybrid ancestors assembled functionally when chaperones were provided (Fig. 2E, S3E-F). However, only the hybrid ancestor comprised of AncD’s LSU and AncC’s SSU (AncD_L_C_S_) retained the ability to assemble without chaperones, suggesting that historical substitutions in the LSU were responsible for chaperone dependence. We also conducted a similar swap of Rubisco-chaperone-ancestor pairs to exclude changes in Raf1 and RbcX as the cause of dependence. Our observations remained the same even when mis-matched chaperone ancestors were utilized, proving that only changes in Rubisco, and not assembly chaperones, were responsible. In addition, improvements to carboxylation kinetics and S_C/O_ were encoded by substitutions in the LSU (Fig. 2F-I). These results demonstrate that the genetic determinants for both improved function and chaperone dependence lie within the LSU.

To pinpoint causal changes in the LSU, we scrutinized all 20 historical substitutions that occurred between AncD to AncC. These were grouped into three spatial clusters of highly conserved substitutions (cluster 1, 2, 3, comprising 3, 3, and 5 substitutions respectively), as well as a fourth cluster (cluster 0 with 9 substitutions) containing all remaining substitutions (Fig. 3A, S4A). We reverted the amino acids of these clusters in AncC back to their ancestral states in AncD (Fig. 3B). Clusters 0 (rv0) or 1 (rv1) variants remained incapable of assembly without chaperones. Reversion of cluster 3 (rv3) weakly restored assembly in the absence of chaperones, requiring combinations with other clusters (rv1^+^3, rv1^+^2^+^3) to amplify this effect (Fig. 3C). Remarkably, reverting cluster 2 (rv2) substitutions alone was sufficient to achieve robust rescue of assembly without chaperones (Fig. 3B-C). We further tested whether reversion of the same three substitutions in cluster 2 would work in an extant enzyme. Rubisco from *G. lithophora* (*GL*), which branches after AncC and is sister to all plastid Rubiscos on our tree (*43*), contains all derived cluster 2 substitutions and is dependent on assembly chaperones (Fig. 3E). Reverting only cluster 2 substitutions in *Gl* Rubisco was likewise able to abolish its dependence on chaperones Raf1 and RbcX. Therefore, cluster 2 substitutions, which lie within the central solvent pore of Rubisco (Fig. 3D), remain as the key determinant for chaperone dependence even in extant descendants.

**Figure 3.**
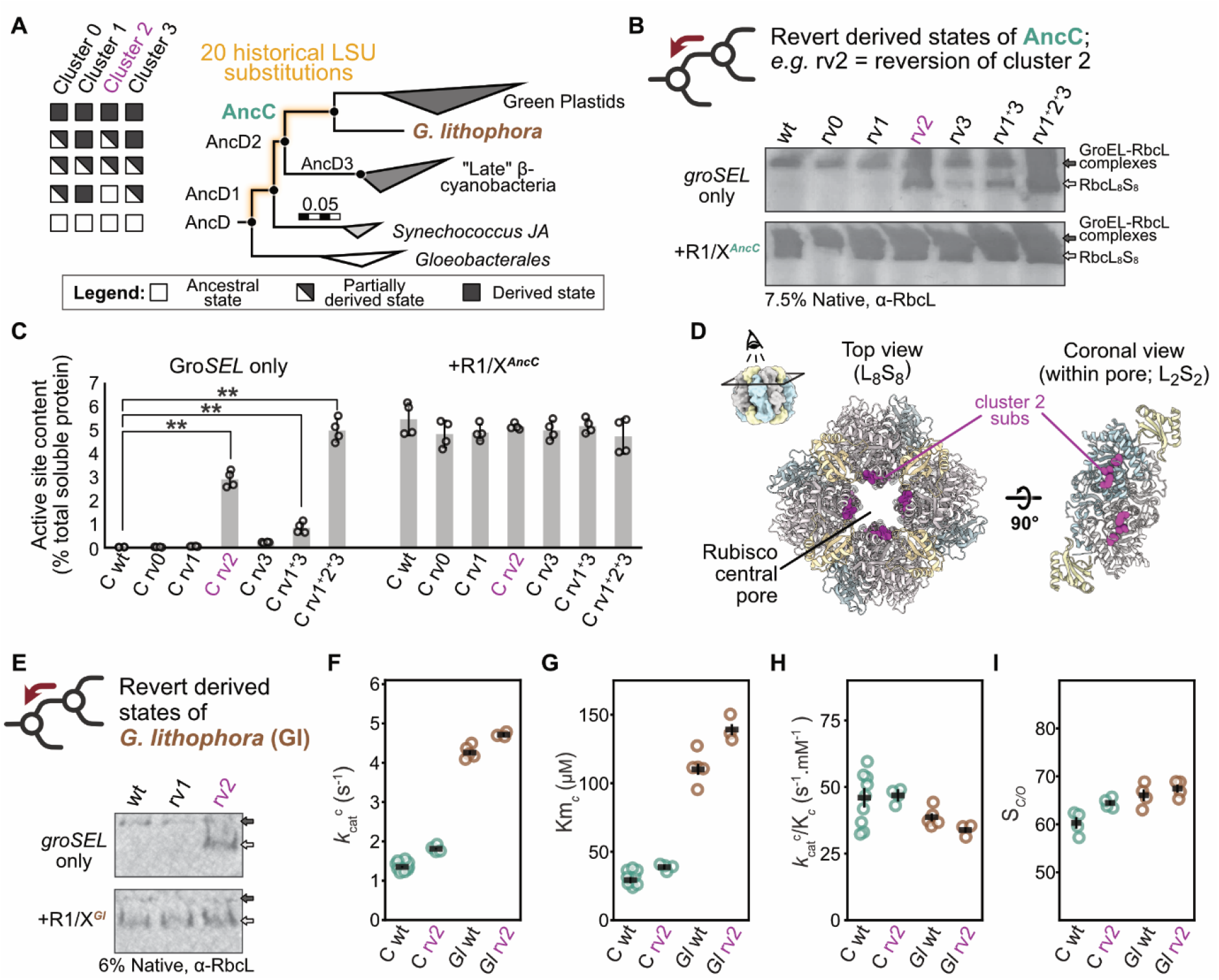
Dependence on chaperones is decoupled from improved catalysis. **(A)** Reduced phylogeny of RbcL between AncD and AncC. The presence or absence of derived substitutions in each cluster is shown on the left, with each row corresponding to the identity of ancestral nodes to the right. **(B)** Native blot analysis of AncC reversion variants carrying ancestral amino acid states at indicated clusters, co-expressed with (bottom row) and without (top row) chaperones. rv: reversion mutant. Black arrows indicate probable chaperonin bound RbcL complexes, White arrows indicate assembled Rubiscos. **(C)** The effect of cluster reversions on assembly yields as in (B), quantified by active site content measurements. Only cluster 2 reversion variant rescues chaperone independent assembly. Significance as tested by one-way ANOVA with post-hoc Tukey HSD test. \*\**P* < 0.01. **(D)** Location of cluster 2 (α4-β5) substitutions in the central solvent channel of Rubisco. On the right, a single L_2_S_2_ dimer is visualized from the coronal plane within the pore. **(E)** Native blot analysis of wild type and reverted mutant Rubisco from the extant cyanobacterium *G. lithophora*. Reversion of cluster 2 but not cluster 1 states rescue self-assembly. **(F-I)** Reversion of cluster 2 substitutions in AncC (wt and rv2 in green) and *G. lithophora* (wt and rv2 in brown) Rubisco do not perturb catalysis. Kinetic parameters extracted from individual fits of technical replicates are shown as hollow circles, with global parameters denoted by a horizontal bar. Error bars represent SEs of the global fit. Exact values of global fits and replicate numbers are indicated in table S1 and kinetic fits in fig. S17.

### Entrenchment of Raf1/RbcX interaction was catalytically neutral

We now had the means to examine if Rubisco’s reliance on chaperones came hand-in-hand with improved catalytic function, as predicted by adaptive buffering. To do this, we compared the enzymatic parameters of AncC (the better, chaperone-dependent enzyme) and AncD (the inferior, but chaperone-independent ancestor) to AncC rv2. Remarkably, the chaperone-independent reversion was as good a catalyst as AncC: we observed no significant changes in S_C/O_ and carboxylation kinetics of AncC rv2 compared to AncC (Fig. 3F-I). To verify that this is not an artefact of our ancestral sequences, we performed the same experiment on the chaperone-independent rv2 variant of the extant *G. lithophora* enzyme (*Gl* rv2). Again, S_C/O_ was unaffected in the rv2 variant of the extant *G. lithophora* enzyme (*Gl* rv2), though *k*_cat_^c^ and K_m_^*CO2*^ increased slightly (Fig. 3F-I). We also did not observe any significant differences in the Michaelis constant for RuBP (K_m_^*RuBP*^) or apparent rate of inhibitor release (*k*_obs_) of reverted variants (Fig. S4F-G). We further tested whether our reversions made the enzyme less stable. To do this, we exposed our variant to elevated temperatures and measured its remaining activity. We found no difference between the reverted variant and its wild-type counterpart (Fig. S4D-E). Furthermore, we also did not find any significant differences in Rubisco assembly yields of the rv2 variant if chaperones were co-produced, indicating that chaperone-dependent variants do not gain added benefits to assembly fidelity from the presence of assembly chaperones (Fig. 3C).

These data imply that entrenchment and catalytic improvements are genetically separable from each other. We next wondered whether we could determine which occurred first: catalytic improvement or chaperone dependence. On our constrained topology there are two nodes between AncD and AncC, which we term AncD1 and AncD2. These proteins already contain cluster 1 and 3 substitutions, but not cluster 2 substitutions, and are not yet dependent on chaperones (Fig. 3A, S3G, S4B-C). If clusters 1 and 3 were responsible for improved catalysis, entrenchment would have come after these improvements. We verified this by experimentally resurrecting AncD2, as well as AncD with clusters 1 and 3 substituted to their states in AncC. Both proteins exhibited elevated S_C/O_ values relative to AncD (Table S1), implying that catalytic improvement at least partially preceded the entrenchment of chaperones. Together, these data show that entrenchment of the assembly chaperones Raf1 and RbcX was neutral on the level of the catalytic parameters of Rubisco. These inferences are robust to the topological uncertainty in our tree: causal substitutions and their impact on catalysis remain unchanged even when examined on the unconstrained maximum likelihood topology (Fig. S2D,S3A-D).

### An epistatic rachet keeps assembly chaperones essential

Within green algae and plants, Raf1 remains essential for Rubisco’s assembly and enhances photosynthetic growth when overexpressed (*29, 44, 45*). However, if only three catalytically neutral reversions are enough to cure Rubisco of this dependence, why are chaperones still necessary in the plastid lineage to this day (*23, 27, 29, 38*)? To answer this, we turned to the youngest of our Rubisco ancestors, AncA. This enzyme represents the LCA of all green algal and plant Rubiscos and is dependent on Raf1 (Fig. 1C). AncA, as well as most other plastid Rubiscos, maintains conservation of the same derived rv2 states as AncC (Fig. S5A). We therefore asked if reverting these substitutions would restore AncA’s ability to assemble without Raf1. AncA rv2, however, remained incapable of assembly without chaperones (Fig. 4B, S5E-F). This implies further changes must have accumulated which deepened dependence on assembly chaperones.

**Figure 4.**
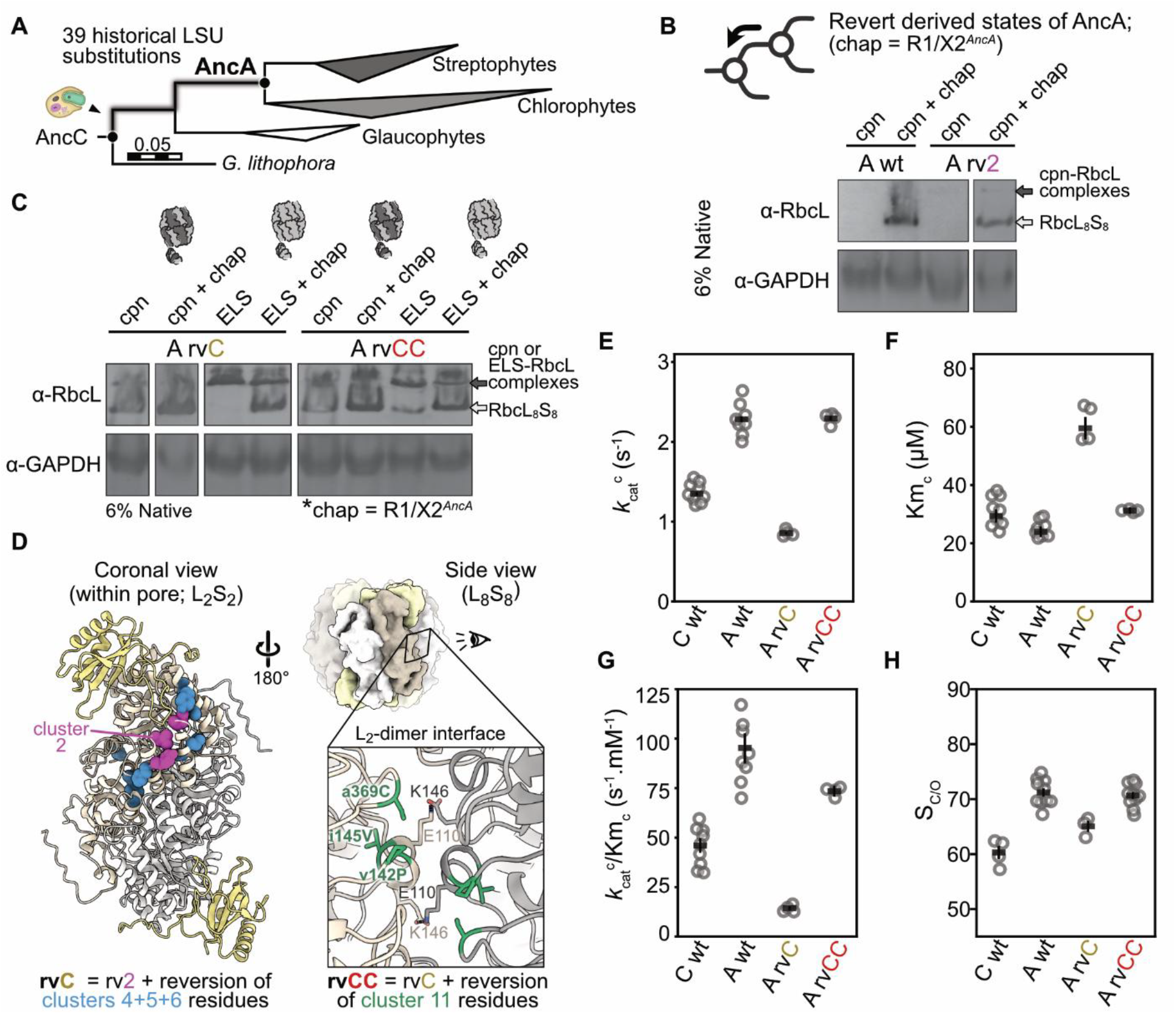
Persistence of chaperone interactions in the green plastid lineage. **(A)** Reduced phylogeny of RbcL between AncC and AncA **(B)** Native blot analysis of AncA wild type and reverted variants as indicated either in the absence or presence of assembly chaperones, **(C)** or with prokaryotic (ELS) or eukaryotic (cpn) chaperonin co-production. cpn: CPN60. chap: Raf1 and RbcX. **(D)** Schematic representation of the location of chaperone and complex plastid type chaperonin entrenching substitutions modelled on AncA’s Rubisco as predicted by Alphafold 3 (*50*). Clusters 4 and 6 surround initial cluster 2 entrenching substitutions, while cluster 5 substitutions occur in proximity to the active site of Rubisco, as additionally depicted in Fig. S6A. Right inset: Cluster 11 substitutions are found at the inter LSU-dimer interface supporting residues key to inter-LSU dimer interactions. **(F-I)** Kinetic parameters exhibited by assembly competent AncA Rubisco variants. Kinetic parameters extracted from individual fits are shown as hollow circles, with global parameters depicted as black horizontal bars. Errors bars denote SEs of the global fit. Replicate numbers are indicated in table. S1 and fitted kinetic curves in fig. S17.

To identify what triggered deeper dependence, we reasoned that the elongated βA-βB loop of plant SSUs, which extends into the central solvent channel of Rubisco, may coordinate with cluster 2 entrenching elements, which may influence the effect of reversions at these sites (Fig. S5B). However, hybrid Rubiscos containing AncC’s SSU, which lacks this elongated lool, remain insoluble without chaperones, excluding a role for the SSU in causing deeper dependence (Fig. S5C). Another plausible contributor could be the folding chaperone: Whereas the simpler prokaryotic GroEL chaperonin suffices for prokaryotic AncC (Fig. S5D; lanes 1 and 2), AncA requires the more structurally complex plastidal chaperonins CPN60α/β (Fig. S5D; lanes 3 and 4). Utilizing GroEL instead of CPN60 also failed to restore chaperone independent assembly of AncA rv2 (Fig. S5F-G, lanes 3). Therefore, we searched once more amongst the 39 historical substitutions that emerged between AncC and AncA, reverting subsets of these in an AncA rv2 background (Fig. S5E, S6A). Three clusters near cluster 2 residues (Fig. 4D) were found to confer weak rescue from chaperones as determined by assembly band intensities (Fig. S5E-F), but rescue was minimal and could be eliminated when co-expression was carried out at slightly elevated temperatures (30°C instead of 28°C) (Fig. S6B-C). In contrast, combining these subsets (AncA rvC; reversion of clusters 2^+^4^+^5^+^6, 11 total substitutions) resulted in robust self-assembly when eukaryotic folding chaperonins were utilized (Fig. S5F). The effect of these further reversions was contingent on also reverting the original cluster 2 sites, since AncA variants without rv2 reversions still required chaperones (Fig. S5F, lanes 6 and 9). These data show that further substitutions entrenched the need for assembly chaperones, and that this was reliant on the original dependence-causing substitutions (also see Supplementary text).

The assembly-competent AncA rvC exhibited compromised catalytic rates compared to AncA (Fig. 4E-H). We therefore identified 3 additional reversions in the inter-dimer interface of the LSU (cluster 11; Fig. 4D) that together with our prior reversions, resulted in a variant (AncA rvCC) with catalytic properties indistinguishable from AncA (Fig. 4E-H), but which could assemble independent of chaperones (Fig. 4C; lanes 5 and 6). Unlike wild-type AncA and our other reversion constructs of it (Fig. 4C lanes 3 and 4, Fig. S5G lanes 5 and 7), AncA rvCC could also fold using GroEL instead of the more complex CPN60 from plants (Fig. 4C; lanes 7 and 8). This demonstrates that it is possible to rid a plastidal Rubisco of its derived folding and assembly chaperone requirements. This prompted us to ask whether we could also rid Rubiscos from *Arabidopsis thaliana* of their need for these chaperones using the same reversions. This experiment, however, failed (Fig. S6D-E), implying that even more entrenching states have accumulated along the lineage to plants since AncA, potentially due to their additional assembly requirements like reliance on BSD2 (*27*).

Together, these results demonstrate that assembly chaperones remain essential because additional entrenching states accumulated along the lineage to plants, and because these states have been ratcheted in through other changes in the protein that makes their reversion functionally unfavourable. At the level of Rubisco’s catalytic function, these additional entrenching substitutions confer no functional advantage. We can at present only speculate about the structural mechanism of Rubisco’s dependence on assembly chaperones. Sites involved in causing the dependence on chaperones cluster at the surface facing the inside of the pore formed by Rubisco complexes, far from known chaperone binding-sites on Rubisco. A plausible mechanism would be that dependence-causing substitutions either make this surface aggregation-prone or unstable due to the loss of specific inter-residue contacts that stabilize the native fold (Fig. S4A, S7). This would pose a problem only during assembly, because this surface is buried and shielded inside Rubisco complexes once assembly is complete. Assembly chaperones, which facilitate the formation of RbcL octamers during assembly, may therefore allow Rubisco to avoid aggregation by minimizing the time it spends with this surface exposed to the cellular environment.

## Discussion

In this work we discover how Rubisco became dependent on its assembly chaperones Raf 1 and RbcX. Addition of these chaperones allowed Rubisco to accumulate substitutions that are neutral when chaperones are present, but highly deleterious when they are not. Though the initial gain of these chaperones was likely adaptive because it boosted assembly yields, the changes that made them essential later were likely non-adaptive, consistent with theoretical models of the evolution multi-layered biological systems (Fig. S8) (*46, 47*).

We note that while this entrenchment was neutral at the level of Rubisco’s catalytic function, a chaperone dependence could be useful, for example, as a means of regulating Rubisco’s abundance in vivo. Such adaptative use of a chaperone dependence itself could also evolve long after the initial dependence arose, as an evolutionary spandrel (*48*). But even if the dependence itself is somehow useful, this cannot explain why Rubisco accumulated substitutions later in in its history that deepened the genetic dependence (i.e. made it more difficult to revert) in an enzyme that could already not assemble at all without chaperones anymore. At present, our only explanation for this process is that the new chaperones unlocked a large and previously inaccessible part of Rubisco’s sequence space, into which it has since been drifting. This is reminiscent of the SSU’s role earlier in Rubisco’s history (*20, 49*). Unlike the SSU, however, Rubisco’s assembly chaperones appear not to have unlocked substitutions that improved Rubisco’s enzymatic function. This is clearly at odds with the theory that chaperones are generally adaptive buffering agents (*2, 5*) and raises the possibility that other protein-chaperone interactions are also the result of similar neutral ratchets.

Rather than adaptive evolutionary buffers, chaperones may be better understood as broad modifiers of their clients’ accessible sequence spaces, which sometimes serve as adaptive buffers and sometimes become essential neutrally. The potential for neutral entrenchment may add a selfish dimension to chaperone evolution: Chaperones can ensure their preservation by becoming essential for other proteins with important functions, even if this doesn’t improve their clients’ functions. The more promiscuous the chaperone, the more certain it is that at least one important protein forgets how to reach its functional conformation without the chaperone. Conversely the more important a potential chaperone client’s function is to an organism, the more effectively purifying selection will preserve any chaperone that becomes essential to that client.

As we show, we can still undo such purposeless increases in cellular complexity by reversing the genetic changes that made these increases essential in history. It may thus be possible to rid modern genomes of any useless cellular bureaucracy that evolution has allowed to accumulate.

## Acknowledgments

We thank Dr. Oliver Mueller-Cajar and members of the Hochberg and Erb labs for critical reading of the manuscript.

## Funding

This research was co-funded by the European Union (ERC, EVOCATION, 101040472 (GKAH, JZYN, DW). The views and opinions expressed are, however, those of the author(s) only and do not necessarily reflect those of the European Union or the European Research Council. Neither the European Union nor the granting authority can be held responsible for them. GKAH, TJE, JZYN, LS, AMK, DW, AH and CG were supported by the Max Planck Society. GKAH gratefully acknowledges funding from a LOEWE Excellence professorship. DW acknowledges funding from the International Max Planck Research School for Principles of Microbial Life: From molecules to cells, from cells to interactions, IMPRS-µLife. AMK was funded by a Long Term Fellowship by the European Molecular Biology Organization, (ALTF 684-2022) and a Marie Skłodowska-Curie fellowship (ECOfix, 101106795).

## Author contributions

Conceptualization: GKAH, TJE, JZYN, LS

Methodology: JZYN, LS, AMK

Investigation: JZYN, DW, LS, AH, CG

Visualization: JZYN

Funding acquisition: GKAH, THE

Project administration: GKAH

Supervision: GKAH

Writing – original draft: JZYN, GKAH

Writing – review & editing: all authors.

## Competing interests

The authors declare that they have no competing interests.

## Data and materials availability

All raw data for kinetic traces, phylogenetic trees, alignments, and ancestral sequences are deposited in the Edmund repository of the Max-Planck Institute and will be made available upon peer-review and publication.

## Notes

### Competing Interest Statement

The authors have declared no competing interest.

### Summary of Updates

1) Affiliations updated to correct error from "Department of Cell Biology, Philipps-Universitat Marburg; Marburg, 35043, Germany" to " Department of Biology, Philipps-Universitat Marburg; Marburg, 35043, Germany". 2) Slight additions and changes to discussion and introduction 3) Minor grammatical corrections to parts of the introduction 4) Update to Funding section in Acknowledgements

