## Supplementary material for "Origin of chaperone dependence and assembly complexity in Rubisco’s biogenesis": Methods and Supplemental Figures

**The PDF file includes:**

Materials and Methods

Supplementary Text

Figs. S1 to S25

Tables S1 to S4

**Other Supplementary Materials for this manuscript include the following:**

MDAR Reproducibility Checklist

#### Supplementary Materials

##### *Materials and Methods*

Phylogenetic analysis and ancestral sequence reconstruction

To retrace the evolutionary histories of Rubisco's large subunit (RbcL), small subunit (RbcS), and assembly chaperones Rubisco accumulation factor 1 (Raf1) and RbcX, amino acid sequences of each homolog were collected from UniPROT, the Joint Genome Institute's (JGI) integrated microbial genome and metagenomes database (IMG), NCBI RefSeq, or the 1000 Plant Genomes Project (OneKP) (*51*) for phylogenetic inference.

For the protein phylogeny of RbcL, 142 amino acid sequences were initially aligned with MUSCLE (v.3.8.31) (*52*) and screened in FastTree (*53*) to distinguish between specific Rubisco forms: Form 1', Form 1'', Form 1anaero (or Form 1F), Form 1C/D red clade Rubiscos, and Form 1A/B green clade associated Rubiscos. Sequences clustering within each clade were inspected to remove lineage-specific indels, before profile alignment with MUSCLE. From this alignment, the Maximum Likelihood (ML) RbcL phylogeny and branch lengths were inferred with RaxML (v8.2.10) (*54*) under the Le-Gascuel (LG) substitution matrix substitution model (*55*) as determined by automatic best-fit evolutionary model selection, with 4 gamma-distributed among-site rate variation categories and empirical base frequencies. As Form 1' and 1'' Rubiscos are known to be early branching forms that do not yet feature or co-assemble with RbcS, Form 1' clade RbcLs were used as the outgroup for rooting, as established by previously published trees (*20*). In the resulting RbcL phylogeny, the *Gloeobacter* clade was not the most basally diverging member of prokaryotic Form 1B β-cyanobacteria RbcLs, with several homologs in branching positions incongruent to the established species relationships for β-cyanobacteria (*42*). While plausible, this topology would require extensive horizontal gene transfer (HGT) events to explain its branching order; and is further inconsistent with the topologies exhibited by the ML trees of both RbcS and RbcX (see below). To allow reconstruction of Rubiscos and chaperones likely to have co-existed within the same ancestral organism, mild topological constraints reflecting the known species relationships of β-cyanobacteria were applied to the ML tree of RbcL. Branch lengths were re-optimized under these constraints in RAxML, yielding the final constrained RbcL phylogeny. The robustness of the ML or constrained tree topology was assessed by inferring 500 non-parametric bootstrap trees with RAxML, from which Felsenstein's and transfer bootstrap values (*56*) were derived and mapped.

For the protein phylogeny of RbcS, 138 amino acid sequences of homologs from species loosely corresponding to taxons utilized in the RbcL tree were collected and aligned in MUSCLE. Lineage specific indels and paralogs of similar sequences (within plastids) were removed and trimmed manually, further discarding likely chloroplast transit peptide (ctp) N-terminal regions based on predictions by targetP (*57*) and comparisons between cyanobacterial and plastid homologs in the multiple sequence alignment (MSA). From this alignment, the ML phylogeny and branch lengths were inferred with RaxML under the LG model as determined by automatic best-fit evolutionary model selection, with 4 gamma-distributed among-site rate variation categories and fixed base frequencies. The tree was rooted with the Form 1anaero clade based on previous reports of their ancestral nature (*20*) and consistency to our

RbcL trees. Although the monophyly of major RbcS clades (1C/D/A/B monophyly) and their relative branching orders were recovered following published Rubisco phylogenies, relationships within Form 1B RbcS gene were highly inconsistent with known species relationships of extant taxa. In particular, glaucophytes, which current phylogenies place as the most basal branching plastid bearing eukaryotes (*58*, *59*), are observed as the earliest Form 1B RbcS nesting within other RbcS of prokaryotic origins from β-cyanobacteria. Topological constraints were therefore applied to impose a branching order consistent with known species relationships, with an additional benefit of matching ancestral RbcS genes with corresponding RbcL ancestors likely to have co-occurred in the same extinct organism. Branch lengths were re-optimized under these constraints in RAxML to yield the final constrained tree. The robustness of the ML or the constrained tree was likewise assessed by inferring 100 non-parametric bootstrap trees with RAxML, from which Felsenstein's and transfer bootstrap values were derived and mapped.

For both Raf1 and RbcX protein phylogenies, a similar approach was utilized with slight modifications. Briefly, Raf1 and RbcX sequences from *G. violaceus* (Raf1: BAC90100.1, RbcX: BAC90098.1) and *A. thaliana* (Raf1.1: Q9LKR8.1, RbcX2: Q8L9X2) were utilized as initial queries to interrogate genomes from representative taxa in each major clade, resulting in 142 and 178 amino acid sequences of Raf1 and RbcX homologs spanning β-cyanobacteria to land plants and green algae respectively. These sequences were first profile-aligned with MAFFT (v7.505) (*60*) before inference of the ML phylogeny using IQ-TREE (v2.2.0.3) (*61*). Model selection was carried by automatic best-fit selection by Modelfinder (*62*) as implemented in IQ-TREE. For Raf1, the ML tree was inferred under the general Q matrix (Q.pfam) model estimated from the Pfam version 31 database (*63*), with 6 among-site rate variation categories under the FreeRate model (*64*, *65*) and fixed base frequencies. For RbcX, the ML tree was inferred under the LG model with uniform among-site rate variation and fixed base frequencies. The basal branching *Gloeobacter* clade was used as an outgroup to root all homologs in both trees. For both trees, branching orders of homologs were observed to be largely consistent with known species relationships, recovering the early divergence of plastids from β-cyanobacteria and a sister relationship of streptophytes and chlorophytes. However, slight differences in the next earliest diverging species were found, though early branching β-cyanobacterial species (*Pseudanabaena* or *Synechococcus JA*) were always observed. To enable resurrection of time-matched ancestors likely to have co-existed in the past, these trees were similarly constrained to reflect the published species relationships of the β-cyanobacteria radiation (*42*) (following *Gloeobacter*, *Synechococcus JA*, followed by a spilt between *Gloeomargarita*/plastids and all other β-cyanobacteria). Branch lengths were then re-optimized under these constraints in IQ-TREE to yield constrained trees. The robustness of the ML or constrained tree was assessed by inferring 100 non-parametric bootstrap trees with RAxML, from which Felsenstein's and transfer bootstrap values were derived and mapped. The constraints applied to Raf1 and RbcX trees were additionally assessed by approximately unbiased (AU) tests (*66*) as implemented in IQ-TREE, which did not reject constrained topologies in both cases and indicates of statistical compatibility with the sequence data.

Based on their respective phylogenies and MSA, ancestral sequences and posterior probabilities (PP) of ancestral states were reconstructed at internal nodes as implemented in IQ-TREE, using the same evolutionary model as per tree inference. For RbcL and RbcS, the CodeML module of PAML (v4.9) (*67*) was utilized with 8 instead of 4 rate categories during calculations. To account for differences at the N or C termini, as well as indel, gap assignment of ancestral sequences was determined using parsimony inference in PAUP (4.0a) (*68*) based on a binary version of the MSA (1 = amino acid, 0 = gap or no residue). The state assignment for each node in the tree (amino acid or gap) was then applied to ancestral sequences to infer the presence or absence of gaps.

Amino acid states with the highest PP at all sites were selected. For Raf1 and RbcX, ancestral amino acid states were inferred with IQ-TREE instead, and gap assignment calculated by Fitch parsimony in PastML (v.1.9.34) (*69*).

#### Molecular biology

The plasmids p11a-*At*Cpn60α/β/Cpn20 and pRSF-*At*RbcLScHis were a gift from Dr. Manajit Hayer-Hartl (*27*) and Dr. Oliver Mueller-Cajar respectively (*70*). All other plasmids were assembled using codon-optimized gene fragments ordered from Twist Bioscience or Genscript Biotech and introduced into domesticated destination vectors. Destination vectors were obtained by modifying pRSFDuet-1 (Novagen), pCDFDuet-1 (Novagen), or pet21(+) plasmid backbones with chosen type II restriction sites for insert cloning. Plasmid backbones were PCR-amplified with custom primers to incorporate desired restriction sites, treated with KLD enzymes (NEB) for ligation and circularization, then recovered and propagated by transformation into an *Escherichia coli* (*Ec*) host. Assemblies were verified by Sanger sequencing (Microsynth) and used for simultaneous digest and ligation of target inserts using the golden-gate cloning methodology with either *Bsa*I, BbsI, or PaqCI (NEB). For Rubisco plasmids (RbcLS-cHis), *rbcL* and *rbcS* gene fragments were synthesized containing upstream ribosome-binding sites without flanking promoter or terminator sequences, with the cassette under control of a single T7/lac promoter and T7 terminator native to the pRSF backbone. *rbcS* genes were designed with a C-terminal 6X-His moiety to enable efficient downstream purification. For assembly chaperones, each gene fragment was synthesized with a flanking T7/lac promoter, ribosome-binding site, T7 terminator, and inserted into the pCDF destination vector in a defined order (Raf1, Raf2, RbcX, BSD2). Short empty filler fragments were utilized when specific chaperones are excluded. For assembly of cyanobacterial chaperonins, *groS* and *groL* gene fragments were synthesized retaining their native operon structure and intergenic regions, but under the control of a single T7/lac promoter and T7 terminator. Chloroplast transit peptides of all genes were removed based of cleavage sites of *At* homologs, or prediction of transit peptide regions by multiple sequence alignment comparisons between plastidal and cyanobacterial homologs. For all cloning steps as required, Luria–Bertani broth (LB; tryptone 10 g/L, NaCl 10 g/L, yeast extract 5 g/L) was utilized to culture *E. coli* NEB 5-α cells in liquid suspensions or solid agar plates supplemented by plasmid-specific antibiotics where appropriate.

#### Chemicals and reagents

All chemicals were of the highest commercially available purity and obtained from Sigma-Aldrich and Carl Roth unless otherwise specified. $NaH^{14}CO_3$, $K^{14}CN$, and D-[2-$^3$H] were obtained from Hartmann Analytics (Germany). Biochemicals for cloning and protein production were obtained from New England Biolabs (NEB) or ThermoFisher Scientific. For determination of specificity constants, [1-$^3$H] RuBP was synthesized enzymatically from D-[2-$^3$H] glucose following published protocols (*49*, *71*). The [$^{14}$C]-labelled Rubisco inhibitor carboxyarabinitol 1,5-bisphosphosphate (CABP) was synthesized from $K^{14}CN$ using 1 mCi per synthesis reaction as previously described (*49*, *72*).

#### Protein purification

For heterologous overexpression, plasmids containing extant and ancestral Rubisco variants, chaperonins, and assembly chaperones were co-transformed into chemically competent *E. coli* BL21 (DE3) cells. With exception of AncE, AncD, AncD f1, AncD f3, $AncD_LC_S$, Gv, and AncA rvC, all other extant or ancestral Rubiscos were purified by co-production with their minimal set of required assembly chaperones as identified during assembly screens. For purification, starter cultures of 2 ml LB with appropriate antibiotics (100 µg/mL carbenicillin: pet11a/28b, 50 µg/mL kanamycin: pRSF, and 50 µg/mL streptomycin: pCDF) were grown overnight at 37 °C prior to inoculation of 800 mL LB cultures. Production cultures were grown for 3 hours at 37 °C in a shaking incubator to reach an $OD_{600}$ of 0.4-0.6 before temperatures were lowered to 23 °C and induced with 0.5 mM isopropyl β-d-thiogalactopyranoside (IPTG). The cells were harvested 18-22 hours after induction by centrifugation (4500 x *g*, 15 min, 4°C) and lysed using a Microfluidizer (Microfluidics) through 4 cycles at 15,000 psi in HisTrap buffer A (50 mM Tris-HCl, pH 8.0, 150 mM NaCl, 25 mM imidazole, 5% v/v glycerol). To obtain clarified lysates, total extracts were spun (22000 x *g*, 20 min, 4 °C) to remove cell debris and aggregates. Soluble fractions were then applied to HisTrap HP 5 mL columns (Sigma-Aldrich) equilibrated with HisTrap buffer A using a peristaltic pump (Hei-FLOW 06, Heidolph). Non-specific binders and contaminants were removed by column washes with 5 column volumes of HisTrap buffer B (50 mM Tris-HCl, pH 8.0, 150 mM NaCl, 50 mM imidazole, 5% v/v glycerol), followed by reverse flow elution with 50% HisTrap buffer C (50 mM Tris-HCl, pH 8.0, 150 mM NaCl, 400 mM imidazole, 5% v/v glycerol) in an NGC (Bio-Rad) system. Rubisco-containing fractions were immediately desalted and subjected to a Superdex 200 gel-filtration column (Cytiva) equilibrated with SEC buffer A (20 mM Tris-HCl, pH 8.0, 150 mM NaCl, 5% v/v glycerol). Rubisco fractions - as determined by SDS-PAGE analysis - were then pooled, concentrated, and flash-frozen for storage at −80 °C. Proteins were quantified by measuring their absorbance at 280 nm using extinction coefficients calculated using the ProtParam tool (*73*). For purification of extant cyanobacteria Rubiscos, all buffers contained 300 mM NaCl throughout and were supplemented with 5 mM 2-Mercaptoethanol (βME) during immobilized metal affinity chromatography (IMAC).

Protein biochemistry

For screening of Rubisco assembly characteristics, plasmids containing the relevant Rubisco, chaperonins, and assembly chaperones were co-transformed and heterologously expressed within *E. coli* BL21 (DE3) cells. Starter cultures of 2 mL LB with appropriate antibiotics were grown overnight at 37 °C prior to inoculation of 50 mL modified ZYM-5052 auto-induction medium cultures (AIM; 25 mM $Na_2HPO_4$, 25 mM $KH_2PO_4$, 50 mM $NH_4Cl$, 5 mM $Na_2SO_4$, 0.5% Glycerol, 0.05% Glucose, 0.2% Lactose, 2 mM $MgSO_4$, tryptone 10 g/L, NaCl 10 g/L, yeast extract 5 g/L) (*74*) containing appropriate antibiotics (if kanamycin, increased to 100 µg/mL). 50 mL cultures were grown for 3 hours at 37 °C in a shaking incubator (200 rpm) to an $OD_{600}$ of 0.4 before temperatures were lowered to 28 °C and incubated for another 22-24 hours. Cells were harvested by centrifugation (4200 x *g*, 15 min, 4°C) and stored dry at -20 °C or immediately lysed by resuspension in 2.5 mL SEC buffer A supplemented with cOmplete™ EDTA-free protease inhibitor cocktail (Roche). Cells were disrupted via five cycles (20 seconds on, 2 min off) of bead-beating homogenization (FastPrep-24™; MP-Bio) at 4 °C, then cleared by centrifugation (17000 x *g*, 20 min, 4 °C) for collection of soluble fractions for analysis.

To assess the presence of assembled Rubisco complexes, native PAGE and western blot analysis was conducted. Clarified lysate concentrations were first quantified by BCA assays (using Pierce™ BCA protein assay kit; Thermo-scientific) and normalized before addition of 5x native sample buffer (62.5 mM Tris-HCl pH 6.8, 0.1% w/v Bromophenol Blue, 40% v/v Glycerol). Sample were loaded on pre-cooled in-house casted 6% native gels or 7.5% Mini-PROTEAN TGX Precast Protein Gels (Bio-Rad) and separated in 1x Tris/Glycine Buffer (2.5 mM Tris-HCl, 19.2 mM glycine, pH 8.3) at 4 °C and 160 V for 100 mins prior to staining with InstantBlue Coomassie Stain. For western blots, separated proteins were transferred onto a PVDF membrane (Immuno-Blot®, Bio-Rad) via semi dry transfer in 1x western blot transfer buffer (25 mM Tris-HCl pH 8.0, 20% v/v methanol, 192 mM glycine) using the standard "SD" preset (30 min, 1.0 A, 25 V) in a Trans-Blot Turbo Transfer System (Bio-Rad). Blotted PVDF membranes were then incubated for 30 min at room temperature (RT) in blocking buffer (5% w/v nonfat, dry milk in 1x TBS (10 mM Tris-HCl, 150 mM NaCl, pH 7.5)) and subsequently washed 3x for 5 min in 1x TBS-T (TBS supplemented with 0.05% v/v Tween 20). Primary antibody hybridization was performed for 1 hour at RT or overnight at 4 °C with gentle shaking with appropriate antibodies (rabbit anti-RbcL, Agrisera AS03-037A, 1 mg/mL 1:8000 in TBS-T; mouse anti-GAPDH, Invitrogen MA5-15738, 1 mg/mL 1:7000 in TBS-T). Hybridized membranes were then washed 3x for 5 mins in 1x TBS-T before application of secondary HRP-conjugated antibodies (Goat anti-Rabbit, Agrisera AS09 602, 0.5 mg/mL 1:5000 in TBS-T; Goat anti-mouse, Sigma-Aldrich 12-349, 1 mg/mL 1:5000 in TBS-T). Following incubation of secondary antibodies for 1 hour at RT with gentle shaking, membranes were washed 3x for 5 mins with 1x TBS-T before detection. Visualization of HRP conjugates was conducted colorimetrically using 1-Step™ Ultra TMB Blotting solution (Thermo-Fisher). To detect GAPDH as a loading control, membranes probed with anti-RbcL were washed in blocking buffer for 50 min at RT and, without stripping, re-probed for GAPDH using the same method described above.

Enzymatic assays

To obtain the Michaelis-Menten constant value for RuBP ($K_M^{RuBP}$), the apparent rate of inhibitor release ($k_{obs}$) from inhibited complexes, and Rubisco activity after incubation at elevated temperatures, spectrophotometric assays were conducted using variants of the coupled-enzyme assay (*75*), with omission of Triose-P isomerase/glycerol-3-phosphate dehydrogenase in the overall enzyme reaction mix. This results in only two NADH oxidation events per RuBP carboxylated. All assays were performed in assay buffer (100 mm Tricine pH 8.0, 5 mM $MgCl_2$) with coupling enzymes (2.5 units/mL creatine phosphokinase (Roche; 10127566001), 2.5 units/mL glyceraldehyde-3-phosphate dehydrogenase (Sigma-Aldrich, G2267) and 2.5 units/mL 3-phosphoglycerate kinase (Sigma-Aldrich, P7634)), 20 mM $NaHCO_3$, 0.5 mM NADH, 1 mM ATP, 10 mM creatine phosphate, and 1 mM RuBP unless indicated otherwise. RuBP was synthesized enzymatically from ribose 5-phosphate (*76*) and purified by anion-exchange chromatography (*71*), as also described previously (*49*). All spectrophotometric assays were performed on different days with individual preparations of activated or inhibited Rubisco complexes to obtain a minimum of 2 technical replicates.

For determination of $K_M^{RuBP}$ values, activated Rubisco complexes (ECM) were formed by incubating 8-20 µM Rubisco active sites in SEC buffer A supplemented with 20 mM $NaHCO_3$ and 10 mM $MgCl_2$ for 1 hour at 25 °C. To initiate reactions, activated Rubisco complexes were added to the enzyme master mix containing varying RuBP concentrations (3.75 – 480 µM) to a final concentration of 0.2 µM active sites (defined as one LSU and one SSU). Turnover was assessed by depletion of NADH absorbance at 340 nm ($Abs_{340}$) at 25 °C using a Cary 60 spectrophotometer and PCB-1500 circulating water bath (Agilent), with reaction rates plotted against RuBP substrate concentrations to obtain Michaelis-Menten curve fits and $K_M$ values.

To assess thermostability and Rubisco activity at elevated temperatures, ECM complexes were first formed as described above. Activated complexes equal to a final concentration of 0.5 µM active sites were incubated at indicated temperatures for 10 mins prior to addition of the enzyme master mix for assay at 25 °C.

For determination of the apparent rate of inhibitor release ($k_{obs}$), inhibited Rubisco complexes (ER) were formed by incubating 8-31.25 µM Rubisco active sites in SEC buffer A containing 4 mM EDTA for an hour prior to further incubation with 1 mM RuBP at 25 °C for another hour. 0.2-0.75 µM ER active sites were then added to the enzyme master mix to initiate assays and track the recovery of Rubisco's function over time. Product accumulation as a function of time ($t$) was fitted using the non-linear function: ($[\text{product}] = v_f \cdot t + ((v_i - v_f)(1 - \exp(-k_{obs} \cdot t)/k_{obs})$ to estimate the initial ($v_i$) and final ($v_f$) steady-state rates, and the observed first-order rate constant ($k_{obs}$) and its associated errors (*77*, *78*).

Radiometric kinetic analysis and Specificity measurements

$^{14}CO_2$ fixation assays (0.5 mL reaction volumes) were conducted at 25 °C within 7.7-ml septum-capped glass scintillation vials in $^{14}C$ assay buffer (100 mm EPPS-NaOH pH 8.0, 20 mm $MgCl_2$, 1 mm EDTA), 10 µg/mL carbonic anhydrase (Sigma-Aldrich, C3934), and 1 mm RuBP. All assay components were equilibrated with $CO_2$-free $N_2$ gas prior to addition of 0.46 to 49.6 mM $NaH^{14}CO_3$ (Hartmann Analytics), corresponding to ~6 to 600 µM $^{14}CO_2$. Dissolved $CO_2$ concentrations were calculated using the Henderson-Hasselbalch equation with pK values for carbonic acid of pKa1 = 6.25 and pKa2 = 10.33, accounting for assay volume and headspace volume (*79*). Purified Rubisco (~10 µM active sites) were activated in assay buffer containing 20 mM $NaHCO_3$, with 20 µL of activation mixture used to initiate assays. Reactions were stopped after 2 min with quenching by 200 µL 50% (v/v) formic acid. The specific activity of the $NaH^{14}CO_3$ stock was measured by complete turnover of 5.4 and 10.8 nanomoles of RuBP using the highest employed $NaH^{14}CO_3$ concentration over 40 mins, with activities ranging from 200 to 600 cpm/nmol RuBP. All reactions were dried using a heat block set at 95 °C before resuspension in 500 µL water and 2.5 mL of ROTISZINT® HighCapacity liquid scintillation cocktail (Carl Roth) for quantification of product formed by scintillation counting with a Beckman LS 6000 scintillation counter.

Active site contents of purified Rubiscos were measured for each data set using [$^{14}$C]-2-carboxy-arabinitol 1,5 bisphosphate (CABP) binding assays (*75*). Assays were conducted with 0.1 nanomoles of activated Rubisco active sites, as estimated from absorbance at 280 nm ($Abs_{280}$). Following incubation of activation mixtures with 4-fold molar excess CABP for an hour, free ligand was separated from bound active sites by size exclusion chromatography either using a custom made SERAgel Pro 250 Column (ISERA), or a self-packed Tricorn™ 5/150 column containing Sephadex G-50 gel filtration medium (Cytiva). Chromatography was conducted on an Agilent 1200 HPLC system with CABP column buffer (25 mM Tricine-KOH pH 8.0, 20 mM $MgCl_2$, 50 mM NaCl), and active site quantification determined by scintillation count of bound [$^{14}$C]-CABP. For determination of Rubisco active site content in clarified *E. coli* lysates, the above was conducted, with modifications to the preparation of activation mixtures. In short, clarified lysate concentrations were first quantified using BCA assays before normalization to 3 or 4 mg/mL, then activated in CABP column buffer supplemented with 20 mM $NaHCO_3$ (1 hour, 25 °C), followed by incubation with CABP for another hour.

To determine Rubisco activity in clarified *E. coli* lysates, $^{14}CO_2$ incorporation assays were likewise adapted. Clarified lysate concentrations were quantified using BCA assays and normalized to 4 mg/mL for activation of Rubisco complexes in $^{14}$C assay buffer supplemented with 20 mM $NaHCO_3$ for an hour at 25 °C. Activation reactions were then diluted to 0.8 mg/mL total protein in 0.5 mL reaction mixtures containing $^{14}$C assay buffer, 1 mM RuBP, 0.02 mg/mL carbonic anhydrase, and 15.6 mM $NaH^{14}CO_3$ (corresponding to ~ 195.4 µM $^{14}CO_2$) to initiate the reaction. Reactions were then carried out as detailed previously, with Rubisco activities determined by the amount of product formation per minute per mg of total protein (nmol/min/mg).

$CO_2/O_2$ specificity ($S_{c/o}$) assays were carried out at 25 °C as previously described (*71*). In brief, purified Rubiscos were assayed in 20 mL of septum-capped glass scintillation vials containing 1 mL 30 mM triethanolamine-HCl pH 8.3, 10 mM $MgSO_4$, and 10 µg/mL carbonic anhydrase. Reactions were equilibrated in a defined gas mixture (999 291 M $O_2$; 709 M $CO_2$) prior to the addition of [1-$^{3}$H]-RuBP for assay initiation. After an hour, reaction products were dephosphorylated by alkaline phosphatase (10 U/reaction) and separated on an Aminex HPX-87H column (Bio-Rad). Eluted peaks corresponding to radiolabeled glycerate and glycolate were quantified by scintillation for calculation of $S_{c/o}$ as previously described (*71*).

***Supplementary Text***

On epistasis and contingency in Rubisco's evolution

Although reverting cluster 2 substitutions alone to their ancestral states successfully cures chaperone dependence in AncC, cluster 2 substitutions were unable to confer strict chaperone reliance when introduced into AncD (Fig. S4B-C; AncD f2). Likewise, introducing clusters 1 and 3 individually (AncD f1, f3) or combined (AncD f1$^{+}$2, f2$^{+}$3) was equally unsuccessful (Fig. S4B). These asymmetric outcomes suggest of context dependent, non-additive influences within Rubisco's mutational and folding energy landscape. In the latter set of combination variants (AncD f1$^{+}$2 or f2$^{+}$3), a mild reduction in assembly yields was observed (Fig. S4C), suggesting that the presence of derived cluster 2 states started becoming destabilizing to folding and assembly, albeit mildly. Since cluster 2 substitutions likely arose after both cluster 1 and 3 (Fig. 4A), this suggests that cluster 2 effects are contingent on prior changes established by clusters 1 and 3. Indeed, strict chaperone dependence only occurs when all three clusters are introduced (AncD f1$^{+}$2$^{+}$3), demonstrating that the phenotype requires coordinated changes across multiple sites (Fig. S4B-C).

We found that these effects are also not limited to specific changes or historical intervals. Within clusters, we observed hints of such influences. Mutational analysis of cluster 2 suggests that m289L and not s262T/M or w283Y as the likely cause of cluster 2 effects (Fig. S3H-I). A plausible structural mechanism for this stems from a reduction of side-chain network interactions in the derived state (Fig. S4A), which may be dependent on historical changes in other cluster 2 residues. Likewise, in our investigation of what led to a deeper chaperone dependence in AncA, we show that the effects of restrictive cluster 4, 5, and 6 substitutions require the presence of initial cluster 2 entrenching substitutions. This was not only in terms of chaperone dependence, but also for stability of the protein: Introducing clusters 4, 5, and 6 into an AncC background without derived cluster 2 states (AncC rv2) led to complete loss of solubility even if assembly chaperones were co-produced (Fig. S5H). This indicates that cluster 2 changes also permitted the presence of AncA restrictive substitutions to be tolerated.

Taken together, these observations illustrate the pervasive influence of epistasis throughout the evolution of Form 1 green lineage Rubiscos, with evolutionary optimizations having to navigate a delicate balance at minimum between assembly fidelity and enzyme function (*80*, *81*). It is clear, however, that multiple solutions may be found, as both chaperone-dependent (*Gl*) and independent (*Gv*, *SynJa*) β-cyanobacterial Rubiscos present comparable $S_{C/O}$ values to AncC (Table S1). Thus, these constraints have likely shaped the divergent kinetic properties and ancillary chaperone requirements exhibited by the distinct lineages of extant Rubiscos today.

The drift barrier hypothesis in Rubisco's evolution and its implications

The evolutionary trajectory of complex chaperone requirements in Rubisco's biogenesis is consistent with the drift barrier theory (Fig. S8) (*46*, *47*), which describes the minimum fitness effect that selection can act on, and is dependent on the effective population size. This barrier acts as a limit on the level of perfection any cellular parameter can attain. In the case of Rubisco, this could have been the assembly fidelity of its biogenesis pathway. Excursions past the drift barrier are possible only temporarily, after which the protein drifts back to the barrier. In Rubisco's history, the addition of the chaperones was likely to have boosted Rubisco's assembly efficiency above the drift barrier, after which Rubisco drifted back towards the barrier and in the process became completely reliant on these chaperones. A further implication of this model is that an increase in the height of the drift barrier through a severe population bottleneck could make it possible to lose the chaperones again, because the fitness cost of this loss would be less visible to selection. Given that simultaneous reversal of 14 (of rvCC) substitutions is unlikely, a plausible scenario involves the emergence of new substitution(s) that may passivate the deleterious effects of entrenching substitutions, or a combination of both. This, for example, may have occurred in green plastid lineages of Form I Rubiscos that experienced secondary or tertiary endosymbiosis events (*82*, *83*), which contain no detectable homologs of some of these chaperones in their current genomes, warranting further investigation. Alternatively, approaches using experimental evolution or targeted protein design may also provide a synthetic solution by leveraging useful substitutions that are already identified from previous studies (*84*–*86*).

Resurrecting Rubisco's chaperone network and constraints

The reconstruction of an ancestral chaperone-client network necessitated the resurrection of all corresponding subunits (RbcL, RbcS) and chaperone (Raf1, RbcX) ancestors. However, we observed that the histories of components differed slightly when inferred from their individual ML gene topologies. This hindered high-confidence pairings of ancestors that were likely to have existed in the same now extinct ancient organism. We therefore examined Rubisco's tree for the possibility of artefactual or systemic errors. For the RbcL tree, we noticed slight incongruences with the known species relationships of β-cyanobacteria (Fig. S10) (*42*, *43*), which would require many HGT events to explain its branching order. Alternatively, this discord could also result from insufficient phylogenetic signal due to the slow evolving nature of RbcL, as well as homoplasy at faster evolving sites. We therefore loosely constrained our RbcL tree to the known species relationships of β-cyanobacteria and plastids (Fig. S9) and explored both (topological) possibilities. We did the same for RbcS (Fig. S12), which suffered from greater violations to the species tree, such as the placement of the early branching plastid bearing glaucophytes at the base of β-cyanobacteria (Fig. S11). To obtain matched ancestors, we also replicated this approach for both chaperones (Fig. S13-S16), even though only minor rearrangements were only required for chaperone trees as they were mostly congruent with a deep placement of early-branching β-cyanobacteria species and plastids. Remarkably, despite constraints to RbcL and RbcS not being supported by tree topology tests, we found that both constrained and ML ancestors evolved similar dependence on chaperones at the same historical node (the node in β-cyanobacteria that unites all plastids; AncC and MLC) and could be abolished using the same reversions. These inferences were also further verified using extant Rubiscos. Thus, we utilized ancestors from constrained topologies for the study.

Several additional obstacles also prevented us from always resurrecting cognate ancestral pairings. As no outgroups for Raf1 and RbcX could be identified, the root of each tree corresponds to the node for AncD’s chaperones, or the LCA of β-cyanobacteria (Fig. S1, S13-S16). Thus, to avoid inferences at the root, we utilized the next immediate ancestor at the node where *Synechococcus sp. JA* (SynJA) diverges in the constrained phylogeny as surrogate to AncD’s chaperones. Likewise, because no ancestral node prior to AncD is accessible, we used AncD chaperones to complement AncE Rubiscos. Further, during our initial tests of ancestral Rubisco assembly, the LCA of all green plastid bearing lineages (containing glaucophytes, chlorophytes, and streptophytes), or AncB (Fig. S9), was barely soluble and readily aggregated in both soluble extracts and small-scale purifications tests despite several optimizations. We therefore omitted this ancestor from further investigations and chose to focus on AncA, which remains dependent on assembly chaperones and was therefore an ideal candidate to elucidate why chaperone interactions were still essential.

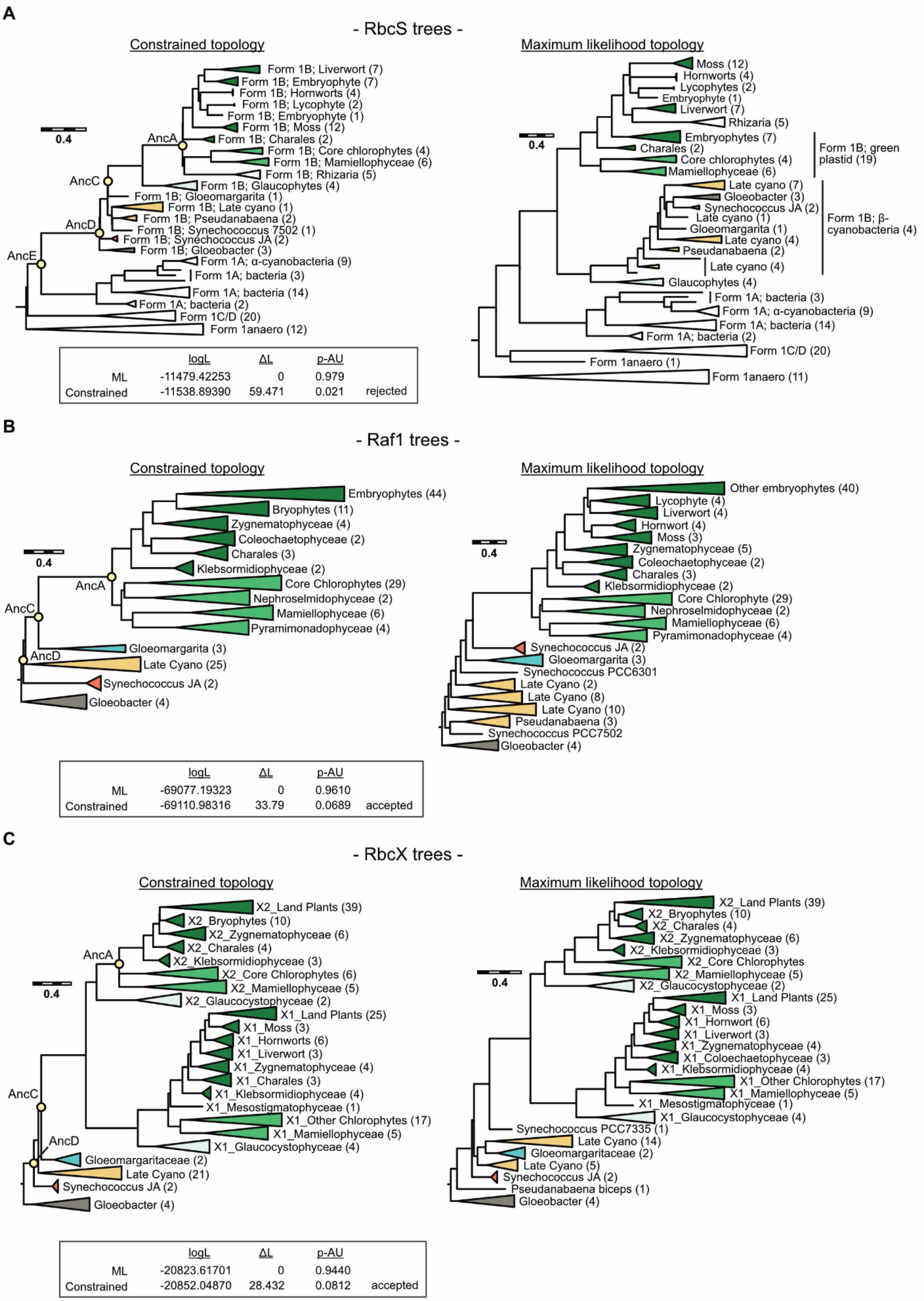
A
- RbcS trees -
Constrained topology
Maximum likelihood topology
AncA
AncC
AncD
AncE
Form 1B; Liverwort (7)
Form 1B; Embryophyte (7)
Form 1B; Hornworts (4)
Form 1B; Lycophyte (2)
Form 1B; Embryophyte (1)
Form 1B; Moss (12)
Form 1B; Charales (2)
Form 1B; Core chlorophytes (4)
Form 1B; Mamiellophyceae (6)
Form 1B; Rhizaria (5)
Form 1B; Glaucophytes (4)
Form 1B; Gloeomargarita (1)
Form 1B; Late cyano (1)
Form 1B; Pseudanabaena (2)
Form 1B; Synechococcus 7502 (1)
Form 1B; Synechococcus JA (2)
Form 1B; Gloeobacter (3)
Form 1A; α-cyanobacteria (9)
Form 1A; bacteria (3)
Form 1A; bacteria (14)
Form 1A; bacteria (2)
Form 1C/D (20)
Form 1anaero (12)
logL ΔL p-AU
ML -11479.42253 0 0.979
Constrained -11538.89390 59.471 0.021 rejected
Moss (12)
Hornworts (4)
Lycophytes (2)
Embryophyte (1)
Liverwort (7)
Rhizaria (5)
Embryophytes (7)
Charales (2)
Core chlorophytes (4)
Mamiellophyceae (6)
Form 1B; green plastid (19)
Late cyano (7)
Gloeobacter (3)
Synechococcus JA (2)
Late cyano (1)
Gloeomargarita (1)
Late cyano (4)
Pseudanabaena (2)
Late cyano (4)
Form 1B; β-cyanobacteria (4)
Glaucophytes (4)
Form 1A; bacteria (3)
Form 1A; α-cyanobacteria (9)
Form 1A; bacteria (14)
Form 1A; bacteria (2)
Form 1C/D (20)
Form 1anaero (1)
Form 1anaero (11)
0.4
B
- Raf1 trees -
Constrained topology
Maximum likelihood topology
Embryophytes (44)
Bryophytes (11)
Zygnematophyceae (4)
Coleochaetophyceae (2)
Charales (3)
Klebsormidiophyceae (2)
Core Chlorophytes (29)
Nephroselmidophyceae (2)
Mamiellophyceae (6)
Pyramimonadophyceae (4)
Gloeomargarita (3)
Late Cyano (25)
Synechococcus JA (2)
Gloeobacter (4)
logL ΔL p-AU
ML -69077.19323 0 0.9610
Constrained -69110.98316 33.79 0.0689 accepted
Other embryophytes (40)
Lycophyte (4)
Liverwort (4)
Hornwort (4)
Moss (3)
Zygnematophyceae (5)
Coleochaetophyceae (2)
Charales (3)
Klebsormidiophyceae (2)
Core Chlorophyte (29)
Nephroselmidophyceae (2)
Mamiellophyceae (6)
Pyramimonadophyceae (4)
Synechococcus JA (2)
Gloeomargarita (3)
Synechococcus PCC6301
Late Cyano (2)
Late Cyano (8)
Late Cyano (10)
Pseudanabaena (3)
Synechococcus PCC7502
Gloeobacter (4)
C
- RbcX trees -
Constrained topology
Maximum likelihood topology
X2_Land Plants (39)
X2_Bryophytes (10)
X2_Zygnematophyceae (6)
X2_Charales (4)
X2_Klebsormidiophyceae (3)
X2_Core Chlorophytes (6)
X2_Mamiellophyceae (5)
X2_Glaucocystophyceae (2)
X1_Land Plants (25)
X1_Moss (3)
X1_Hornworts (6)
X1_Liverwort (3)
X1_Zygnematophyceae (4)
X1_Charales (3)
X1_Klebsormidiophyceae (4)
X1_Mesostigmatophyceae (1)
X1_Other Chlorophytes (17)
X1_Mamiellophyceae (5)
X1_Glaucocystophyceae (4)
Gloeomargaritaceae (2)
Late Cyano (21)
Synechococcus JA (2)
Gloeobacter (4)
logL ΔL p-AU
ML -20823.61701 0 0.9440
Constrained -20852.04870 28.432 0.0812 accepted
X2_Land Plants (39)
X2_Bryophytes (10)
X2_Charales (4)
X2_Zygnematophyceae (6)
X2_Klebsormidiophyceae (3)
X2_Core Chlorophytes
X2_Mamiellophyceae (5)
X2_Glaucocystophyceae (2)
X1_Land Plants (25)
X1_Moss (3)
X1_Hornwort (6)
X1_Liverwort (3)
X1_Zygnematophyceae (4)
X1_Coleochaetophyceae (3)
X1_Klebsormidiophyceae (4)
X1_Other Chlorophytes (17)
X1_Mamiellophyceae (5)
X1_Mesostigmatophyceae (1)
X1_Glaucocystophyceae (4)
Synechococcus PCC7335 (1)
Late Cyano (14)
Gloeomargaritaceae (2)
Late Cyano (5)
Synechococcus JA (2)
Pseudanabaena biceps (1)
Gloeobacter (4)

**Fig. S1. Reconstructing synchronized ancestors of Rubisco and its assembly chaperone network.**

Comparison between constrained (left column) and Maximum Likelihood (right column) topologies of **(A)** RbcS, **(B)** Raf1, and **(C)** RbcX protein phylogenies. Minor topological constraints according to known species relationships were applied to obtain constrained topologies as described in the Methods section. Approximately unbiased (AU) tree topology test results are shown under each constrained tree. Full uncollapsed trees are shown in fig. S9-S16.

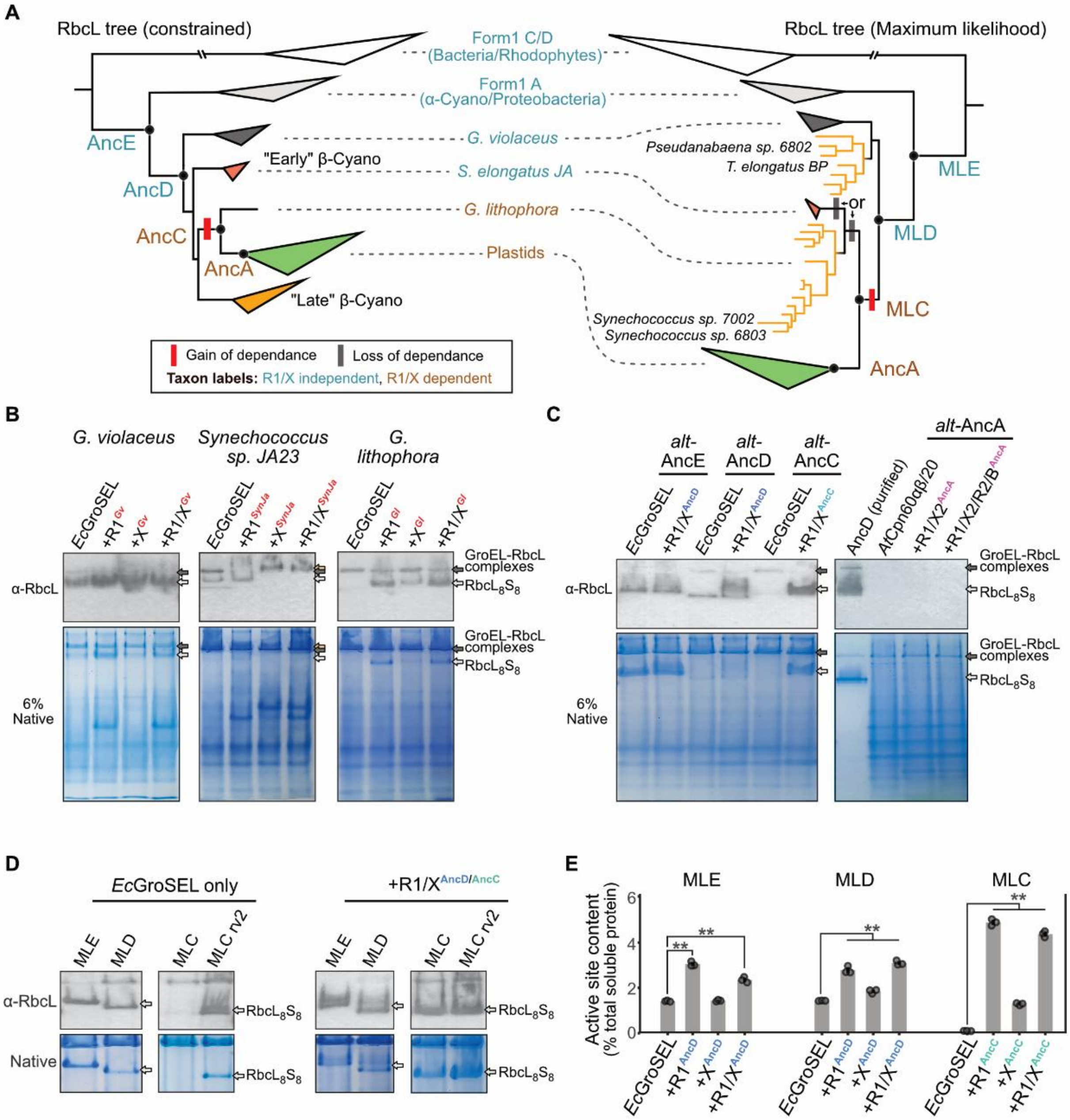

**Fig. S2. Robustness of chaperone dependence to topological and reconstruction uncertainty.**

**(A)** Differences between the constrained (left) and ML (right) RbcL trees are indicated. The basal branching *Gloeobacter* clade is in dark grey. Full uncollapsed constrained and ML trees are shown in fig. S9 and S10. **(B)** Examination of extant cyanobacteria Rubisco assembly characteristics from indicated species with cognate chaperones, **(C)** less-likely ancestors from the constrained topology, and **(D)** alternative ancestors from the ML topology. **(E)** Assembled Rubisco active site yields from ML ancestral Rubiscos from soluble lysate extracts quantified with binding of the radiolabelled inhibitor $^{14}$C-CABP. The boost to assembly yields afforded by Raf1 remains largely unchanged from ancestors obtained in the constrained topology. Significance as tested by one-way ANOVA with post-hoc Tukey test. Non-significant: *n.s.*; $^{**}P < 0.01$.

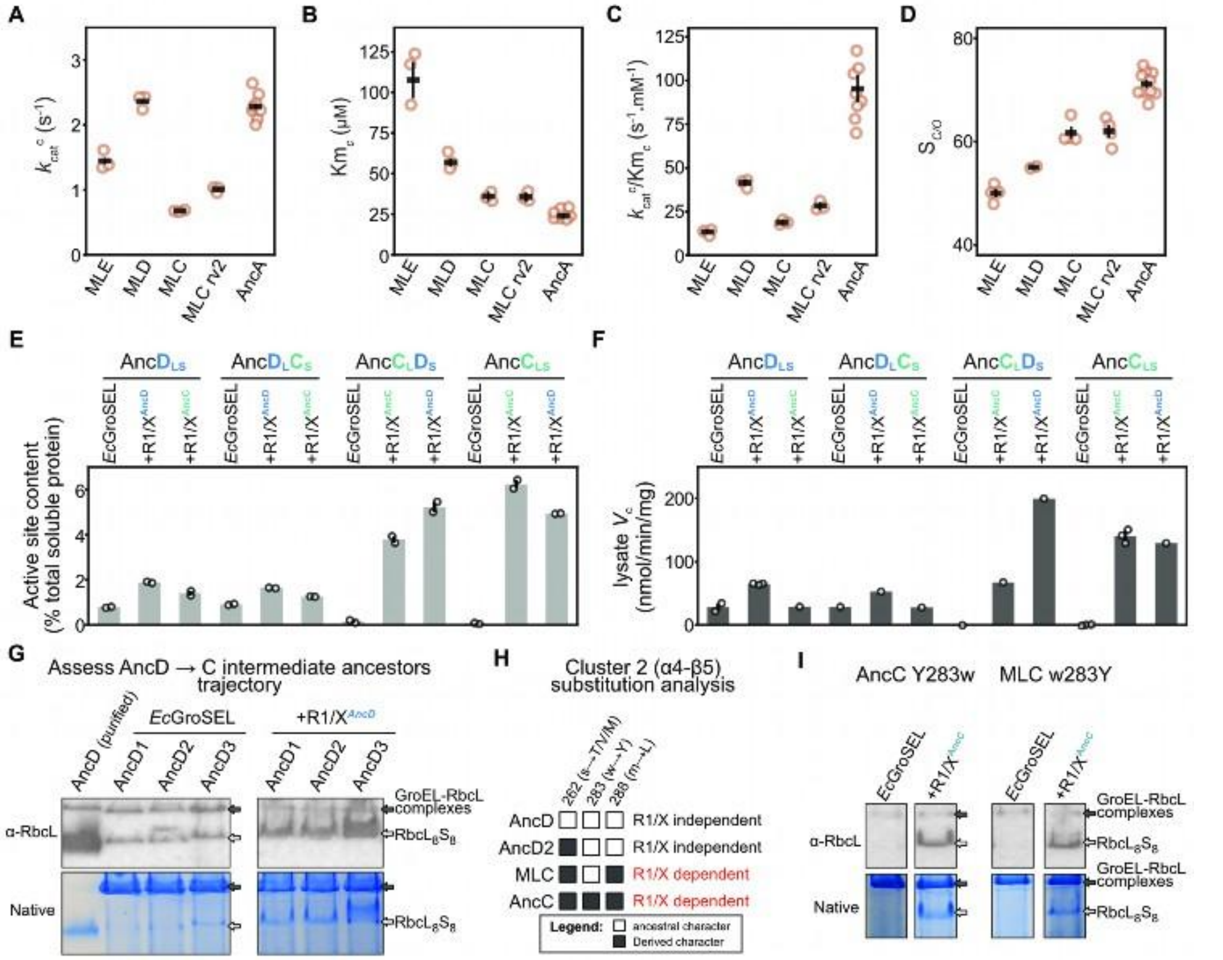

**Fig. S3. Genetic determinants of chaperone dependency and improved function**

**(A-C)** Carboxylation kinetics and **(D)** specificity of maximum likelihood Rubisco ancestors. Catalytic parameters of the chaperone-independent cluster 2 reversion variant (MLC rv2) remain similar to the chaperone dependent maximum-likelihood ancestor C (MLC). Kinetic parameters extracted from individual fits are depicted as hollow circles, with globally fitted values shown as horizontal black bars. Errors bars denote SEs of the global curve fit. Replicate numbers indicated in Table. S1. Fitted kinetic curves are shown in fig. S17. **(E)** Assembly characteristics of chimeric Rubiscos containing swapped subunits quantified by the amount of assembled Rubisco active sites and **(F)** activity in lysates. Changes in the LSU dictates chaperone requirement and is independent of chaperone identity. **(G)** Native PAGE and blot analysis of Rubisco ancestors inferred at intermediate ancestral nodes between AncD and AncC as depicted in Fig. 3A. All intermediate ancestors were co-produced with AncD's ancestral chaperonins when indicated. Black arrows indicate potential chaperonin-bound RbcL complexes while white arrows for assembled $L_8S_8$ complex. **(H)** Dissection of cluster 2 substitutions and their relative contributions to chaperone dependence as observed from wild type or mutant variants containing differing derived amino acid states. **(I)** Reversion of only the derived tyrosine 283 to its ancestral tryptophan in AncC verifies that MLC remains dependent on chaperones when analysed by Native blot and PAGE.

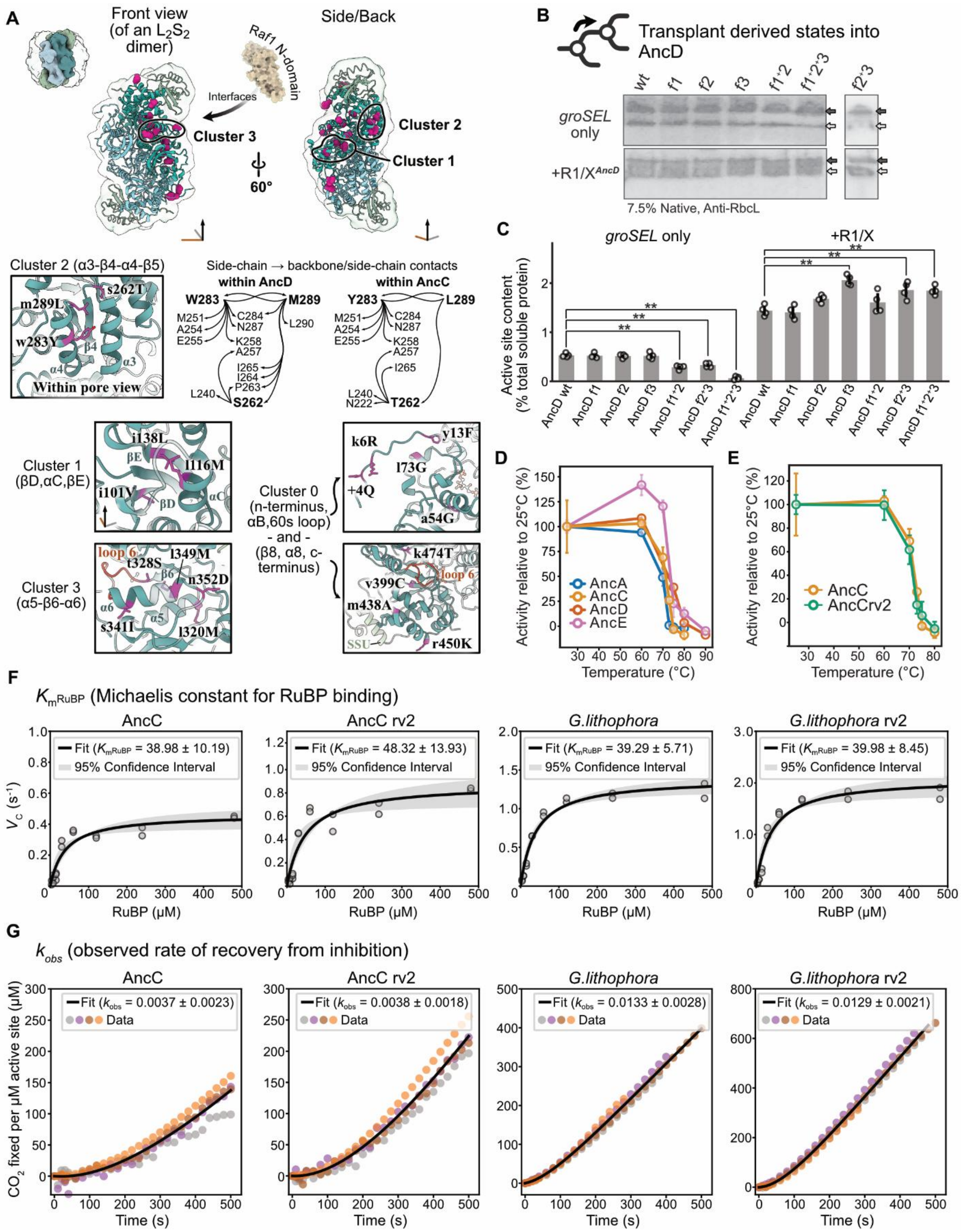

**Fig. S4. Reversion of chaperone entrenching substitutions do not significantly impact the activity of Rubisco.**

**(A)** Structural representation of historical substitutions (purple spheres) between AncD to AncC. An overview of the assembled $L_8S_8$ complex is shown in the top left, with LSU and SSU chains from a single $L_2S_2$ dimer colorized and expanded with a corresponding view from the front and the side. Only substitutions from one LSU in the dimer are depicted for clarity. Substitutions in spatial proximity are grouped into clusters as indicated. Cluster 3 substitutions bracket the active site closure element (loop 6) and lie at the interface with the assembly chaperone Raf1 (*87*). Raf1 N-terminal domain solid fill representation obtained from PDB: 6KKM. Bottom: A close-up of each cluster as visualized with an AncC Alphafold 3 (*50*) prediction. Small letters denote ancestral states while capitalized letters denote derived states. For cluster 2, side chain contacts made by historical substitutions are compared between AncD and AncC. A reduction in the co-interaction network is observed, with m289L eliminating multiple interactions between α3-β4 and α4 previously present in AncD. **(B)** Native blot analysis and **(C)** active site quantification of AncD wild-type or indicated variant Rubiscos co-produced with or without chaperones. Values are expressed as a percentage of total soluble protein content in lysates as determined by BCA measurements. Error bars represent SDs. A significant difference from the wild-type value is indicated by a double asterisk (One-way ANOVA, posthoc Tukey test, $p < 0.01$). **(D)** Thermal profiles of ancestral Rubisco activities after incubation at indicated temperatures for 10 min prior to measurements at fixed substrate concentrations. Activated Rubisco preparations were pre-formed before incubation. **(E)** No changes were observed when wild type and reverted variants of AncC were tested. Similarly, **(F)** the Michaelis constant for RuBP ($K_M^{RuBP}$) and **(G)** apparent rate of inhibitor release ($k_{obs}$) remained relatively unchanged between wild type and rv2 variants. For both (F) and (G), replicates are shown as circles with a global fit overlaid.

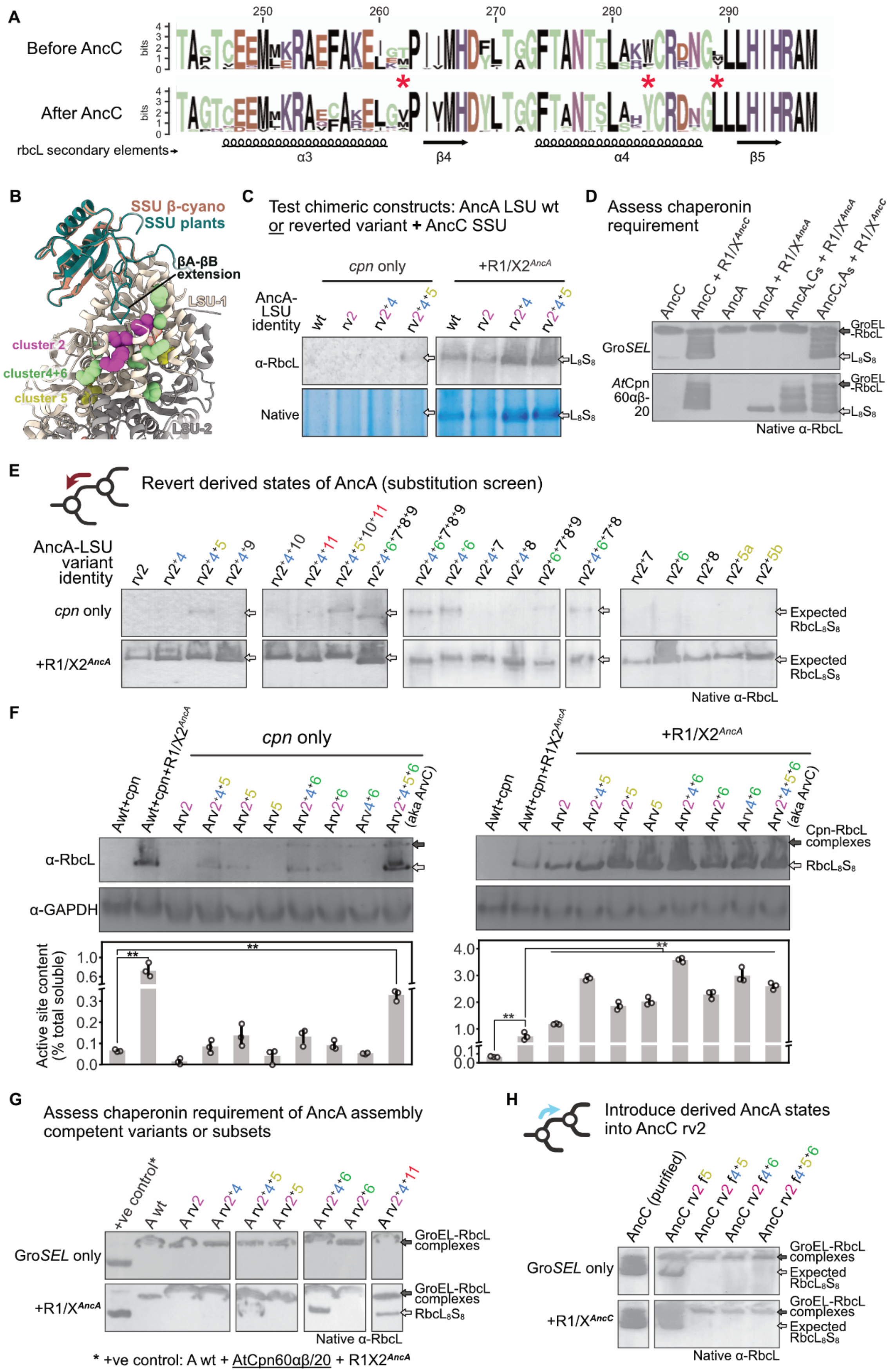
A
Before AncC
After AncC
rbcL secondary elements
α3
β4
α4
β5
B
SSU β-cyano
SSU plants
βA-βB extension
LSU-1
cluster 2
cluster4+6
cluster 5
LSU-2
C
Test chimeric constructs: AncA LSU wt or reverted variant + AncC SSU
cpn only
+R1/X2AncA
AncA-LSU identity
α-RbcL
Native
D
Assess chaperonin requirement
GroSEL
AtCpn 60αβ-20
GroEL-RbcL
Native α-RbcL
E
Revert derived states of AncA (substitution screen)
AncA-LSU variant identity
cpn only
+R1/X2AncA
Expected RbcL8S8
Native α-RbcL
F
cpn only
+R1/X2AncA
α-RbcL
α-GAPDH
Cpn-RbcL complexes
RbcL8S8
Active site content (% total soluble)
G
Assess chaperonin requirement of AncA assembly competent variants or subsets
GroSEL only
+R1/XAncA
GroEL-RbcL complexes
RbcL8S8
Native α-RbcL
* +ve control: A wt + AtCpn60αβ/20 + R1X2AncA
H
Introduce derived AncA states into AncC rv2
GroSEL only
+R1/XAncC
GroEL-RbcL complexes
Expected RbcL8S8
Native α-RbcL

**Fig. S5. Restrictive substitutions that further entrench chaperone interactions are encoded in the LSU.**

**(A)** Consensus identities of amino acid sequences in the α4-β5 region of the LSU before and after the ancestral node corresponding to AncC. Cluster 2 entrenching substitutions (denoted with a purple asterisk) remain highly conserved after AncC. **(B)** Differences in the βA-βB loop of SSUs from plants compared to β-cyanobacteria. The SSU from CABP-inhibited *Spinacia oleracea* Rubisco (PDB: 1RBO) was superimposed on the SSU from CABP-inhibited *Thermosynechoccus elongatus* Rubisco (PDB: 3ZXW). The elongated βA-βB loop extends into the central solvent channel of Rubisco and may coordinate with residues proximal to cluster 2 entrenching substitutions or subsequent restrictive substitutions. **(C)** Native blot and PAGE analysis of chimeric AncA wild type or indicated rv2 LSU variants complemented with AncC's SSU, or the **(D)** co-production of wild type AncA or AncC and hybrid variants AncC$_L$A$_S$ or AncA$_L$C$_S$ with indicated chaperones and chaperonins. A role for both changes in triggering a deeper reliance on chaperones is ruled out since assembly competence remains dependent on the identity of the LSU. **(E)** Native blot analysis of AncA rv2 reverted variants identifies substitutions in clusters 4, 5, and 6 as responsible for further entrenchment. The division of AncA rv2 B substitutions into subsets (Bb and Ba) failed to recapitulate the rescue phenotype. **(F)** Native blot analysis and active site content quantification of AncA wild type and reverted variants as indicated when chaperones were co-produced. All reverted variants are observed to exhibit significantly enhanced assembly yields in *E. coli* in the presence of chaperones as compared to wild-type AncA chaperone dependent yields. Significance as determined by one-way ANOVA with post-hoc Tukey HSD test. $^{**}P < 0.01$. **(H)** AncC mutants containing transplants of derived amino acid states from restrictive substitution clusters are insoluble even when chaperones are co-produced, with exception of cluster B that remains assembly competent. **(G)** Native blot analysis of AncA wild type or indicated reverted variants co-produced with *Ec*GroSEL or *At*Cpn60/α/β/20. Restrictive substitutions that kept chaperones essential are also involved in the requirement for complex plastidal chaperonin action, although chaperones remain necessary when the simpler prokaryotic chaperonin is utilized.

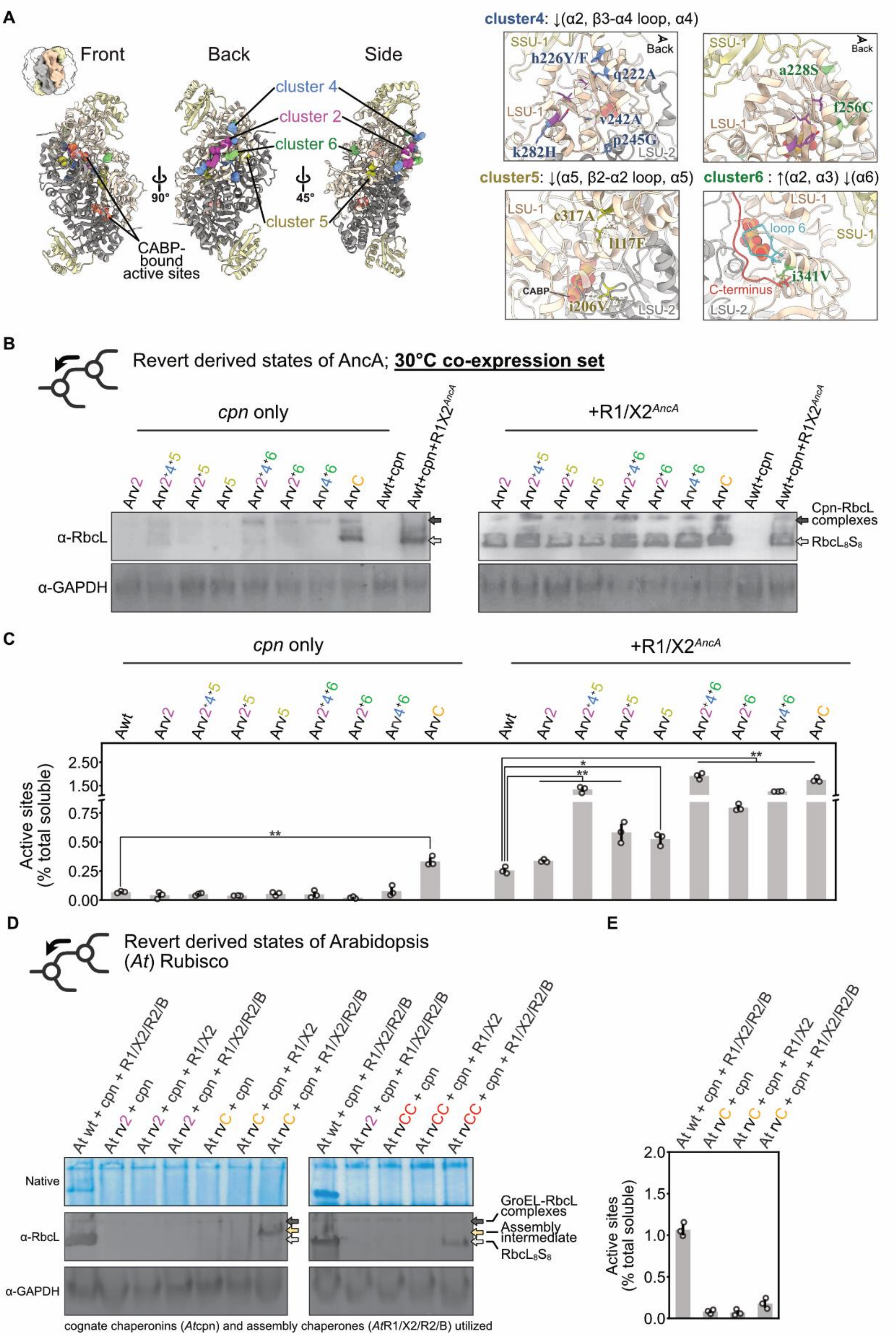

A
Front
Back
Side
cluster 4
cluster 2
cluster 6
cluster 5
CABP-bound active sites
cluster4: ↓(α2, β3-α4 loop, α4)
cluster5: ↓(α5, β2-α2 loop, α5)
cluster6 : ↑(α2, α3) ↓(α6)
B
Revert derived states of AncA; 30°C co-expression set
cpn only
+R1/X2AncA
α-RbcL
α-GAPDH
Cpn-RbcL complexes
RbcL8S8
C
Active sites (% total soluble)
D
Revert derived states of Arabidopsis (At) Rubisco
Native
GroEL-RbcL complexes
Assembly intermediate
cognate chaperonins (Atcpn) and assembly chaperones (AtR1/X2/R2/B) utilized
E

551

**Fig. S6. Analysis of restrictive LSU substitutions within Rubiscos of the green plastid lineage.**

**(A)** Structural depiction of entrenching substitutions occurring between AncC and AncA. A single $L_2S_2$ dimer colorized corresponding to the overall assembled Rubisco complex in the top left is shown from different angles. Only substitutions from one LSU in the dimer are represented for clarity. Close-ups of each cluster and their corresponding structural elements are shown on the right, based on Alphafold predictions of AncA's structure. The position of CABP within the active sites of AncA is modelled by a structural overlay of CABP-inhibited spinach Rubisco on AncA. Small letters denote ancestral states, capitalized letters denote derived states. **(B)** Native blot analysis and **(C)** active site quantification of extracted lysates of AncA assembly competent variants utilized in Fig. S5F, but with growth and co-expression at 30°C instead of 28°C. Weak rescue from assembly chaperones for subsets of AncA rvC in Fig. S5F is abolished, with assembly yields of AncA rvC diminished. **(D)** Examination of assembly characteristics of *A. thaliana* wild type, rv2, rvC, and rvCC reversion variants. Indicated chaperone combinations were co-produced. Both rvC and rvCC no longer restore chaperone independent assembly, mirroring the outcome observed with rv2 only reversions in AncA and indicates of additional restrictive substitutions having emerged. A higher molecular weight band above assembled $L_8S_8$ complexes was observed when *At* rvC is co-produced with all assembly chaperones, but not when only Raf1 and RbcX is given. **(E)** Active site quantification of *At* rvC band reveals little or no functional Rubisco complexes, indicative instead of intermediate assemblies.

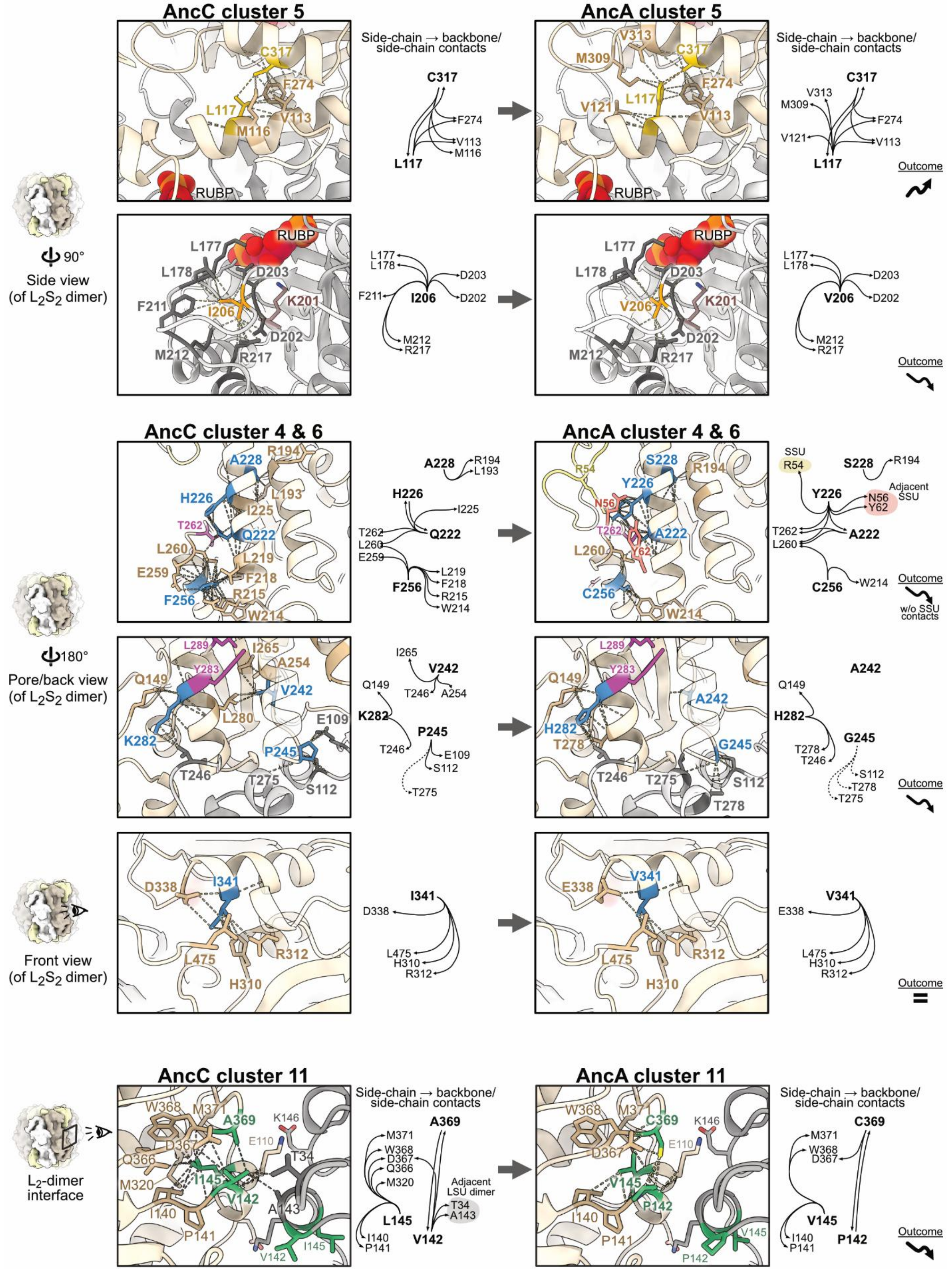
AncC cluster 5
AncA cluster 5
Side-chain → backbone/ side-chain contacts
C317
F274
L117
M116
V113
RUBP
M309
V313
V121
Outcome
Side view (of L2S2 dimer)
90°
L177
L178
D203
F211
I206
K201
D202
M212
R217
V206
AncC cluster 4 & 6
AncA cluster 4 & 6
A228
R194
L193
H226
I225
T262
Q222
L260
L219
E259
F218
F256
R215
W214
S228
SSU
R54
Y226
Adjacent SSU
N56
Y62
A222
C256
w/o SSU contacts
180°
Pore/back view (of L2S2 dimer)
L289
Y283
I265
A254
Q149
V242
L280
K282
E109
P245
T246
T275
S112
A242
H282
T278
G245
Front view (of L2S2 dimer)
D338
I341
L475
H310
R312
E338
V341
=
AncC cluster 11
AncA cluster 11
L2-dimer interface
W368
M371
A369
K146
E110
D367
Q366
M320
I145
T34
V142
A143
I140
P141
L145
Adjacent LSU dimer
C369
V145
P142

**Fig. S7. Structural analysis of AncA restrictive substitutions**

Structural representations of the effects of historical substitutions occurring between AncC to AncA. Intra-residue contacts are observed to be mostly reduced within RbcL, as denoted by the downward trending arrow, upward trending arrow, or equal sign in the bottom right of each panel row. All interpretations were modelled in the context of assembled Rubisco complexes ($L_8S_8$) using Alphafold 3 (*50*) with ancestral Rubisco sequences.

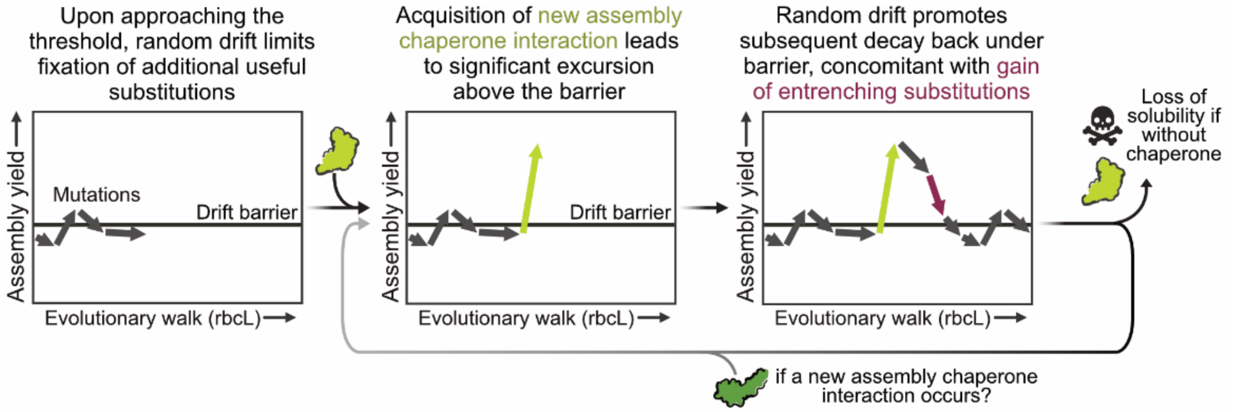

**Fig. S8. Neutral emergence of chaperone dependence in Rubisco's biogenesis**

Proposed mechanism for the evolution of increased complexity in the assembly of green lineage Rubiscos explained through the drift barrier hypothesis (*47*). Left panel: Once a certain level of optimization is achieved (in Rubisco's assembly pathway), the benefits of further useful substitutions become too small to overcome the power of random genetic drift, resulting in an equilibrium at the drift barrier. Middle panel: The initial introduction of assembly chaperones considerably bolstered assembled yields and fidelity of the pathway, resulting in a significant excursion above the drift barrier. Right panel: This advantage is gradually diminished and falls back towards the barrier as random drift dominates. If, during this decay, a mutation that renders chaperones essential gets fixed by drift, the interaction becomes entrenched without further benefit. Should new chaperone interactions be subsequently introduced, such as the additional chaperones that plants also require, this process may repeat, promoting the stepwise evolution of a complex, multilayered assembly pathway (*7*).

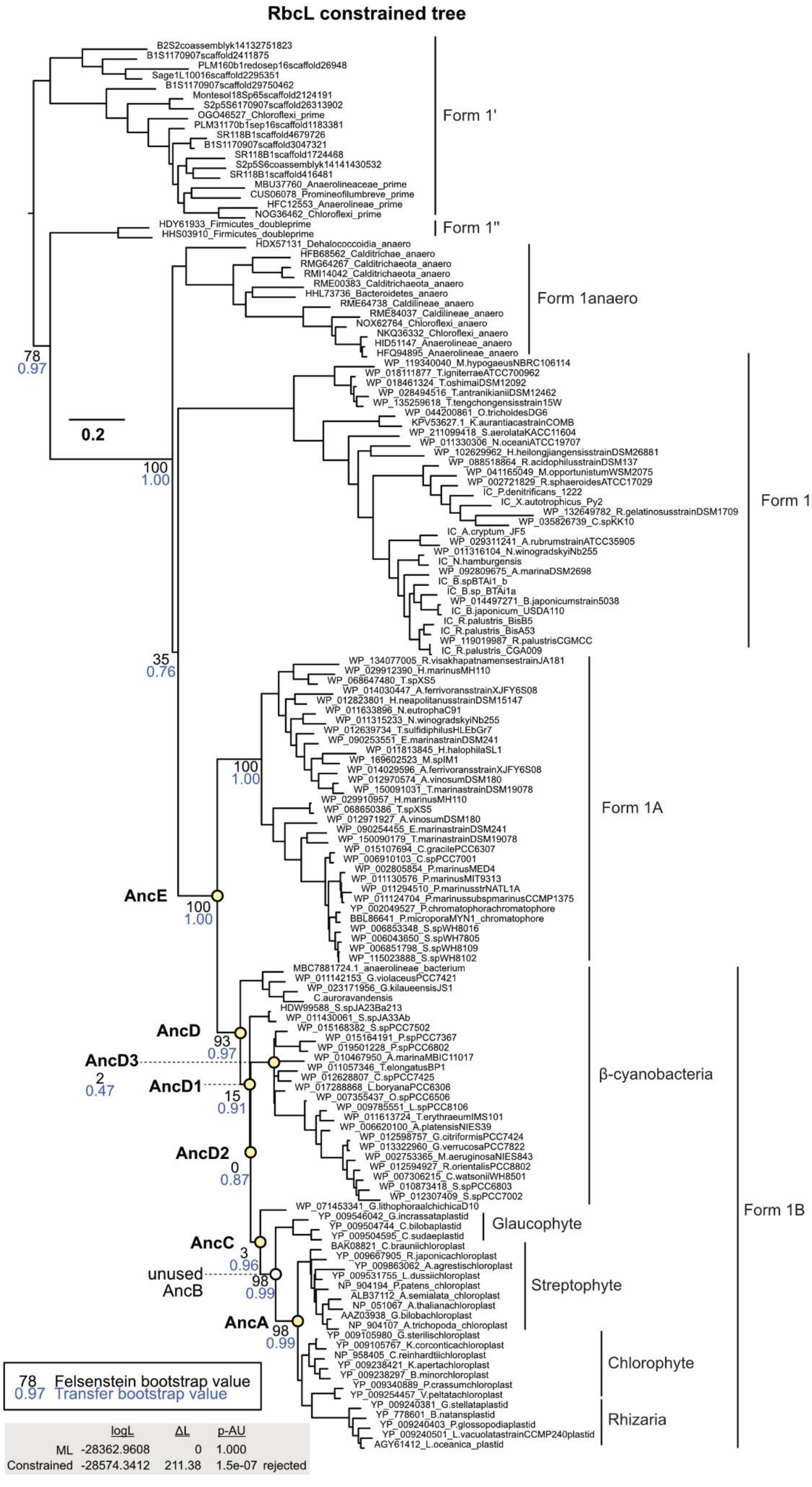

RbcL constrained tree
B2S2coassemblyk14132751823
B1S1170907scaffold2411875
PLM160b1redosep16scaffold26948
Sage1L10016scaffold2295351
B1S1170907scaffold29750462
Montesol18Sp65scaffold2124191
S2p5S6170907scaffold26313902
OGO46527_Chloroflexi_prime
PLM31170b1sep16scaffold1183381
SR118B1scaffold4679726
B1S1170907scaffold3047321
SR118B1scaffold1724468
S2p5S6coassemblyk14141430532
SR118B1scaffold416481
MBU37760_Anaerolineaceae_prime
CUS06078_Promineofilumbreve_prime
HFC12553_Anaerolineae_prime
NOG36462_Chloroflexi_prime
Form 1'
HDY61933_Firmicutes_doubleprime
HHS03910_Firmicutes_doubleprime
Form 1''
HDX57131_Dehalococcoidia_anaero
HFB68562_Calditrichae_anaero
RMG64267_Calditrichaeota_anaero
RMI14042_Calditrichaeota_anaero
RME00383_Calditrichaeota_anaero
HHL73736_Bacteroidetes_anaero
RME64738_Caldilineae_anaero
RME84037_Caldilineae_anaero
NOX62764_Chloroflexi_anaero
NKQ36332_Chloroflexi_anaero
HID51147_Anaerolineae_anaero
HFQ94895_Anaerolineae_anaero
Form 1anaero
78
0.97
0.2
100
1.00
WP_119340040_M.hypogaeusNBRC106114
WP_018111877_T.igniterraeATCC700962
WP_018461324_T.oshimaiDSM12092
WP_028494516_T.antranikianiiDSM12462
WP_135259618_T.tengchongensisstrain15W
WP_044200861_O.trichoidesDG6
KPV53627.1_K.aurantiacastrainCOMB
WP_211099418_S.aerolataKACC11604
WP_011330306_N.oceaniATCC19707
WP_102629962_H.heilongjiangensisstrainDSM26881
WP_088518864_R.acidophilusstrainDSM137
WP_041165049_M.opportunistumWSM2075
WP_002721829_R.sphaeroidesATCC17029
IC_P.denitrificans_1222
IC_X.autotrophicus_Py2
WP_132649782_R.gelatinosusstrainDSM1709
WP_035826739_C.spKK10
IC_A.cryptum_JF5
WP_029311241_A.rubrumstrainATCC35905
WP_011316104_N.winogradskyiNb255
IC_N.hamburgensis
WP_092809675_A.marinaDSM2698
IC_B.spBTAi1_b
IC_B.sp_BTAi1a
WP_014497271_B.japonicumstrain5038
IC_B.japonicum_USDA110
IC_R.palustris_BisB5
IC_R.palustris_BisA53
WP_119019987_R.palustrisCGMCC
IC_R.palustris_CGA009
Form 1C/D
35
0.76
WP_134077005_R.visakhapatnamensestrainJA181
WP_029912390_H.marinusMH110
WP_068647480_T.spXS5
WP_014030447_A.ferrivoransstrainXJFY6S08
WP_012823801_H.neapolitanusstrainDSM15147
WP_011633896_N.eutrophaC91
WP_011315233_N.winogradskyiNb255
WP_012639734_T.sulfidiphilusHLEbGr7
WP_090253551_E.marinastrainDSM241
WP_011813845_H.halophilaSL1
WP_169602523_M.spIM1
WP_014029596_A.ferrivoransstrainXJFY6S08
WP_012970574_A.vinosumDSM180
WP_150091031_T.marinastrainDSM19078
WP_029910957_H.marinusMH110
WP_068650386_T.spXS5
WP_012971927_A.vinosumDSM180
WP_090254455_E.marinastrainDSM241
WP_150090179_T.marinastrainDSM19078
WP_015107694_C.gracilePCC6307
WP_006910103_C.spPCC7001
WP_002805854_P.marinusMED4
WP_011130576_P.marinusMIT9313
WP_011294510_P.marinusstrNATL1A
WP_011124704_P.marinussubspmarinusCCMP1375
YP_002049527_P.chromatophorachromatophore
BBL86641_P.microporaMYN1_chromatophore
WP_006853348_S.spWH8016
WP_006043650_S.spWH7805
WP_006851798_S.spWH8109
WP_115023888_S.spWH8102
100
1.00
Form 1A
AncE
100
1.00
MBC7881724.1_anaerolineae_bacterium
WP_011142153_G.violaceusPCC7421
WP_023171956_G.kilaueensisJS1
C.auroravandensis
HDW99588_S.spJA23Ba213
WP_011430061_S.spJA33Ab
WP_015168382_S.spPCC7502
WP_015164191_P.spPCC7367
WP_019501228_P.spPCC6802
WP_010467950_A.marinaMBIC11017
WP_011057346_T.elongatusBP1
WP_012628807_C.spPCC7425
WP_017288868_L.boryanaPCC6306
WP_007355437_O.spPCC6506
WP_009785551_L.spPCC8106
WP_011613724_T.erythraeumIMS101
WP_006620100_A.platensisNIES39
WP_012598757_G.citriformisPCC7424
WP_013322960_G.verrucosaPCC7822
WP_002753365_M.aeruginosaNIES843
WP_012594927_R.orientalisPCC8802
WP_007306215_C.watsoniiWH8501
WP_010873418_S.spPCC6803
WP_012307409_S.spPCC7002
β-cyanobacteria
AncD
93
0.97
AncD3
2
0.47
AncD1
15
0.91
AncD2
0
0.87
WP_071453341_G.lithophoraalchichicaD10
YP_009546042_G.incrassataplastid
YP_009504744_C.bilobaplastid
YP_009504595_C.sudaeplastid
Glaucophyte
AncC
3
0.96
BAK08821_C.brauniichloroplast
YP_009667905_R.japonicachloroplast
YP_009863062_A.agrestischloroplast
YP_009531755_L.dussiichloroplast
NP_904194_P.patens_chloroplast
ALB37112_A.semialata_chloroplast
NP_051067_A.thalianachloroplast
AAZ03938_G.bilobachloroplast
NP_904107_A.trichopoda_chloroplast
Streptophyte
unused
AncB
98
0.99
AncA
98
0.99
YP_009105980_G.sterilischloroplast
YP_009105767_K.corconticachloroplast
NP_958405_C.reinhardtiichloroplast
YP_009238421_K.apertachloroplast
YP_009238297_B.minorchloroplast
YP_009340889_P.crassumchloroplast
YP_009254457_V.peltatachloroplast
Chlorophyte
YP_009240381_G.stellataplastid
YP_778601_B.natansplastid
YP_009240403_P.glossopodiaplastid
YP_009240501_L.vacuolatastrainCCMP240plastid
AGY61412_L.oceanica_plastid
Rhizaria
Form 1B
78 Felsenstein bootstrap value
0.97 Transfer bootstrap value
logL ΔL p-AU
ML -28362.9608 0 1.000
Constrained -28574.3412 211.38 1.5e-07 rejected

**Fig. S9. Constrained RbcL phylogeny**

Uncollapsed phylogeny of Rubisco's large subunit, based on the maximum-likelihood tree in fig. S10 but with minor topological constraints applied to reflect known species relationships between β-cyanobacteria (*42*) and green plastids (*88*, *89*). Rubisco forms are labelled, with host species indicated for Form 1B Rubiscos. Felsenstein bootstrap values (black) and transfer bootstrap values (blue) are indicated next to nodes of interest. Scale bar: average substitution per site. Ancestral nodes (yellow) of AncE, AncD, AncD1, AncD2, AncD3, AncC, AncB, and AncA are indicated. NCBI Reference Sequence, UniprotKB, or IMG gene identifiers are listed. Statistical analysis of the constrained topology as assessed by approximately unbiased (AU) test as implemented in IQ-TREE is shown at the bottom.

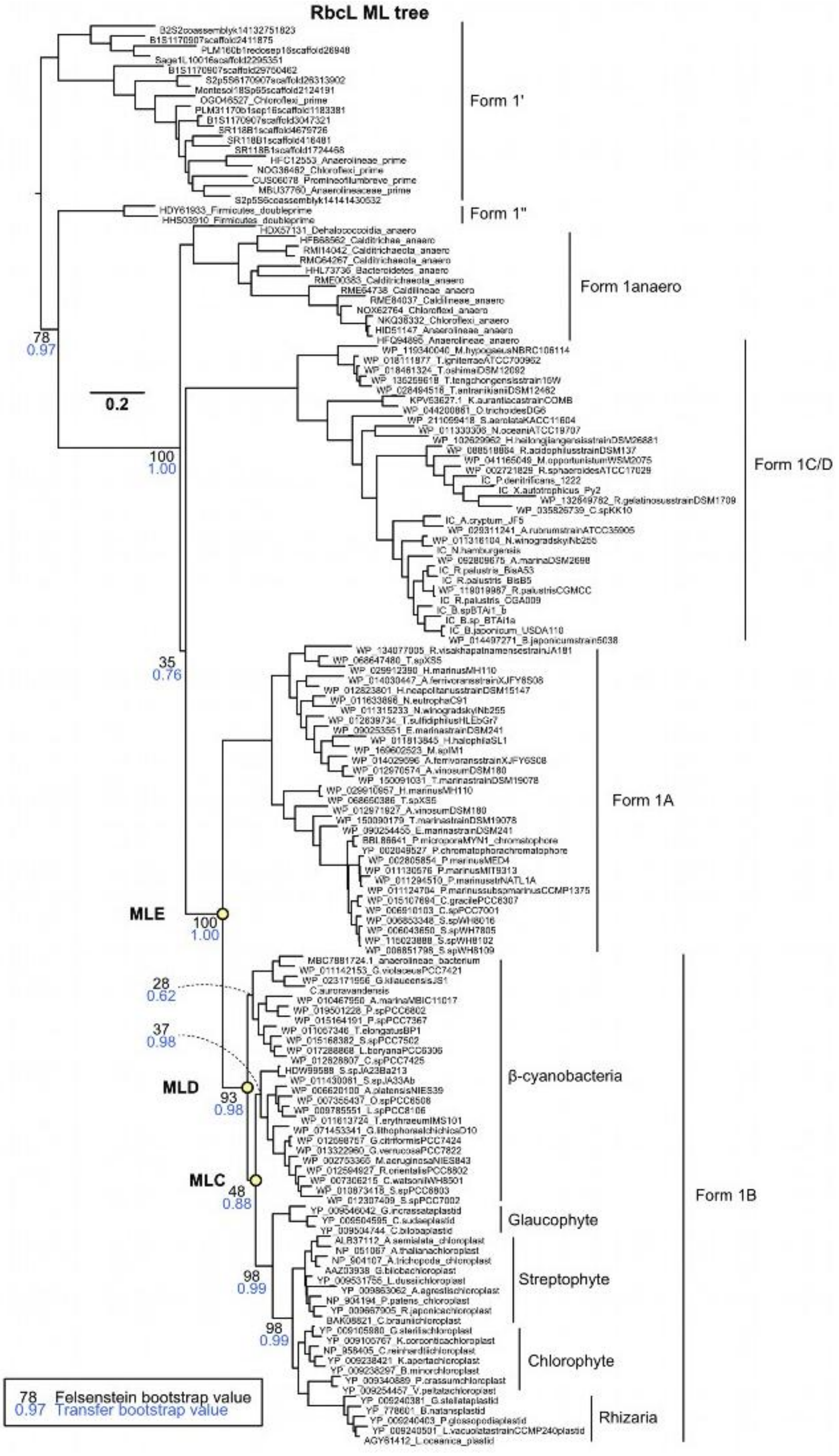
RbcL ML tree
Form 1'
Form 1''
Form 1anaero
Form 1C/D
Form 1A
β-cyanobacteria
Glaucophyte
Streptophyte
Chlorophyte
Rhizaria
Form 1B
MLE
MLD
MLC
78
0.97
0.2
100
1.00
35
0.76
100
1.00
28
0.62
37
0.98
93
0.98
48
0.88
98
0.99
98
0.99
78 Felsenstein bootstrap value
0.97 Transfer bootstrap value

**Fig. S10. Maximum-likelihood RbcL phylogeny**

Uncollapsed maximum-likelihood phylogeny of Rubisco's large subunit. Rubisco forms are labelled, with host species indicated for Form 1B Rubiscos. Felsenstein bootstrap values (black) and transfer bootstrap values (blue) are indicated at selected nodes of interest. Scale bar: average substitution per site. Ancestral nodes (yellow) that best correspond to resurrected Rubisco large subunits from the constrained topology are shown as MLE, MLD, or MLC. NCBI Reference Sequence, UniprotKB, or IMG gene identifiers are listed.

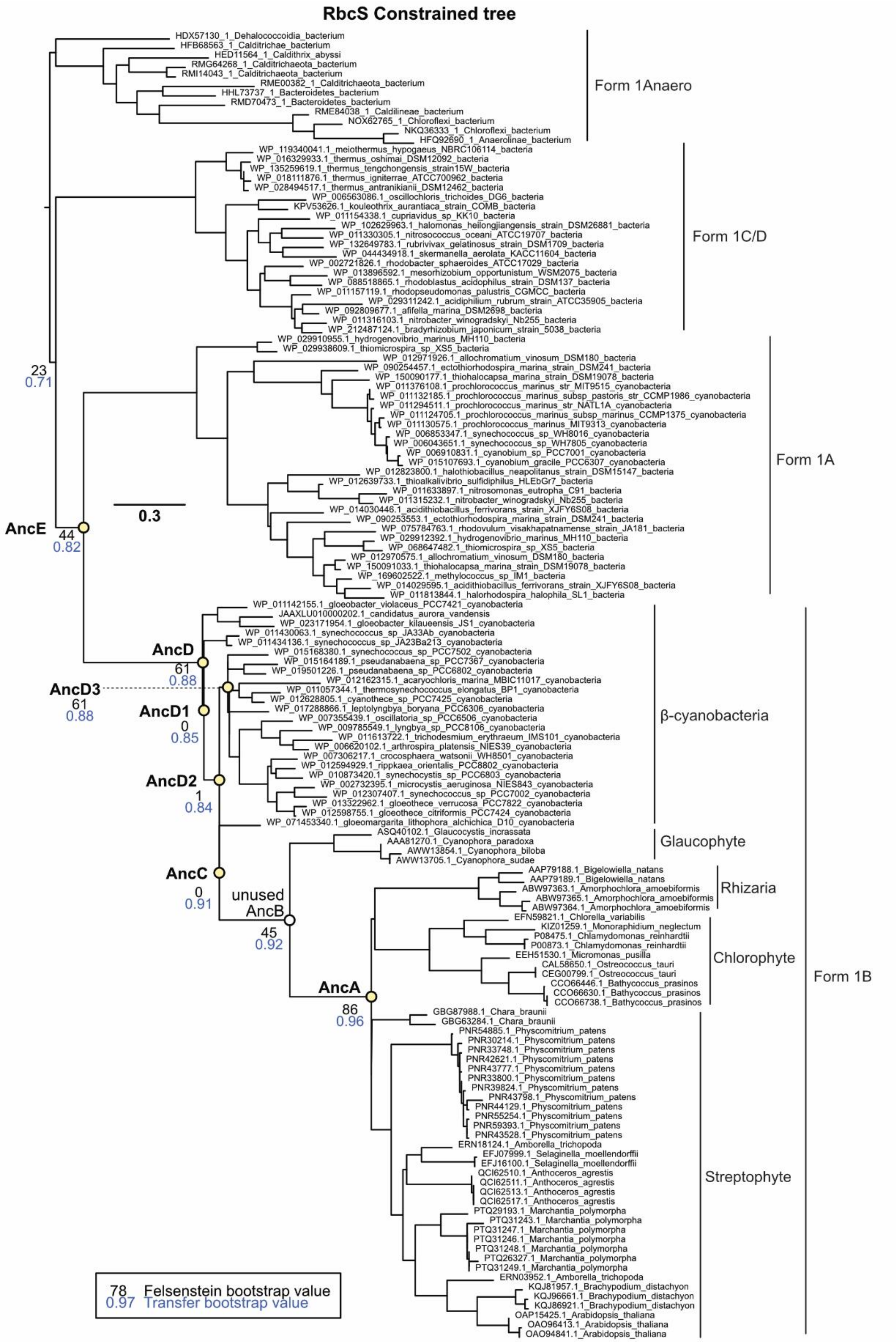

RbcS Constrained tree
HDX57130_1_Dehalococcoidia_bacterium
HFB68563_1_Calditrichae_bacterium
HED11564_1_Caldithrix_abyssi
RMG64268_1_Calditrichaeota_bacterium
RMI14043_1_Calditrichaeota_bacterium
RME00382_1_Calditrichaeota_bacterium
HHL73737_1_Bacteroidetes_bacterium
RMD70473_1_Bacteroidetes_bacterium
RME84038_1_Caldilineae_bacterium
NOX62765_1_Chloroflexi_bacterium
NKQ36333_1_Chloroflexi_bacterium
HFQ92690_1_Anaerolinae_bacterium
Form 1Anaero
WP_119340041.1_meiothermus_hypogaeus_NBRC106114_bacteria
WP_016329933.1_thermus_oshimai_DSM12092_bacteria
WP_135259619.1_thermus_tengchongensis_strain15W_bacteria
WP_018111876.1_thermus_igniterrae_ATCC700962_bacteria
WP_028494517.1_thermus_antranikianii_DSM12462_bacteria
WP_006563086.1_oscillochloris_trichoides_DG6_bacteria
KPV53626.1_kouleothrix_aurantiaca_strain_COMB_bacteria
WP_011154338.1_cupriavidus_sp_KK10_bacteria
WP_102629963.1_halomonas_heilongjiangensis_strain_DSM26881_bacteria
WP_011330305.1_nitrosococcus_oceani_ATCC19707_bacteria
WP_132649783.1_rubrivivax_gelatinosus_strain_DSM1709_bacteria
WP_044434918.1_skermanella_aerolata_KACC11604_bacteria
WP_002721826.1_rhodobacter_sphaeroides_ATCC17029_bacteria
WP_013896592.1_mesorhizobium_opportunistum_WSM2075_bacteria
WP_088518865.1_rhodoblastus_acidophilus_strain_DSM137_bacteria
WP_011157119.1_rhodopseudomonas_palustris_CGMCC_bacteria
WP_029311242.1_acidiphilium_rubrum_strain_ATCC35905_bacteria
WP_092809677.1_afifella_marina_DSM2698_bacteria
WP_011316103.1_nitrobacter_winogradskyi_Nb255_bacteria
WP_212487124.1_bradyrhizobium_japonicum_strain_5038_bacteria
Form 1C/D
WP_029910955.1_hydrogenovibrio_marinus_MH110_bacteria
WP_029938609.1_thiomicrospira_sp_XS5_bacteria
WP_012971926.1_allochromatium_vinosum_DSM180_bacteria
WP_090254457.1_ectothiorhodospira_marina_strain_DSM241_bacteria
WP_150090177.1_thiohalocapsa_marina_strain_DSM19078_bacteria
WP_011376108.1_prochlorococcus_marinus_str_MIT9515_cyanobacteria
WP_011132185.1_prochlorococcus_marinus_subsp_pastoris_str_CCMP1986_cyanobacteria
WP_011294511.1_prochlorococcus_marinus_str_NATL1A_cyanobacteria
WP_011124705.1_prochlorococcus_marinus_subsp_marinus_CCMP1375_cyanobacteria
WP_011130575.1_prochlorococcus_marinus_MIT9313_cyanobacteria
WP_006853347.1_synechococcus_sp_WH8016_cyanobacteria
WP_006043651.1_synechococcus_sp_WH7805_cyanobacteria
WP_006910831.1_cyanobium_sp_PCC7001_cyanobacteria
WP_015107693.1_cyanobium_gracile_PCC6307_cyanobacteria
WP_012823800.1_halothiobacillus_neapolitanus_strain_DSM15147_bacteria
WP_012639733.1_thioalkalivibrio_sulfidiphilus_HLEbGr7_bacteria
WP_011633897.1_nitrosomonas_eutropha_C91_bacteria
WP_011315232.1_nitrobacter_winogradskyi_Nb255_bacteria
WP_014030446.1_acidithiobacillus_ferrivorans_strain_XJFY6S08_bacteria
WP_090253553.1_ectothiorhodospira_marina_strain_DSM241_bacteria
WP_075784763.1_rhodovulum_visakhapatnamense_strain_JA181_bacteria
WP_029912392.1_hydrogenovibrio_marinus_MH110_bacteria
WP_068647482.1_thiomicrospira_sp_XS5_bacteria
WP_012970575.1_allochromatium_vinosum_DSM180_bacteria
WP_150091033.1_thiohalocapsa_marina_strain_DSM19078_bacteria
WP_169602522.1_methylococcus_sp_IM1_bacteria
WP_014029595.1_acidithiobacillus_ferrivorans_strain_XJFY6S08_bacteria
WP_011813844.1_halorhodospira_halophila_SL1_bacteria
Form 1A
0.3
23
0.71
AncE
44
0.82
WP_011142155.1_gloeobacter_violaceus_PCC7421_cyanobacteria
JAAXLU010000202.1_candidatus_aurora_vandensis
WP_023171954.1_gloeobacter_kilaueensis_JS1_cyanobacteria
WP_011430063.1_synechococcus_sp_JA33Ab_cyanobacteria
WP_011434136.1_synechococcus_sp_JA23Ba213_cyanobacteria
WP_015168380.1_synechococcus_sp_PCC7502_cyanobacteria
WP_015164189.1_pseudanabaena_sp_PCC7367_cyanobacteria
WP_019501226.1_pseudanabaena_sp_PCC6802_cyanobacteria
WP_012162315.1_acaryochloris_marina_MBIC11017_cyanobacteria
WP_011057344.1_thermosynechococcus_elongatus_BP1_cyanobacteria
WP_012628805.1_cyanothece_sp_PCC7425_cyanobacteria
WP_017288866.1_leptolyngbya_boryana_PCC6306_cyanobacteria
WP_007355439.1_oscillatoria_sp_PCC6506_cyanobacteria
WP_009785549.1_lyngbya_sp_PCC8106_cyanobacteria
WP_011613722.1_trichodesmium_erythraeum_IMS101_cyanobacteria
WP_006620102.1_arthrospira_platensis_NIES39_cyanobacteria
WP_007306217.1_crocosphaera_watsonii_WH8501_cyanobacteria
WP_012594929.1_rippkaea_orientalis_PCC8802_cyanobacteria
WP_010873420.1_synechocystis_sp_PCC6803_cyanobacteria
WP_002732395.1_microcystis_aeruginosa_NIES843_cyanobacteria
WP_012307407.1_synechococcus_sp_PCC7002_cyanobacteria
WP_013322962.1_gloeothece_verrucosa_PCC7822_cyanobacteria
WP_012598755.1_gloeothece_citriformis_PCC7424_cyanobacteria
WP_071453340.1_gloeomargarita_lithophora_alchichica_D10_cyanobacteria
β-cyanobacteria
AncD
61
0.88
AncD3
61
0.88
AncD1
0
0.85
AncD2
1
0.84
ASQ40102.1_Glaucocystis_incrassata
AAA81270.1_Cyanophora_paradoxa
AWW13854.1_Cyanophora_biloba
AWW13705.1_Cyanophora_sudae
Glaucophyte
AncC
0
0.91
unused
AncB
45
0.92
AAP79188.1_Bigelowiella_natans
AAP79189.1_Bigelowiella_natans
ABW97363.1_Amorphochlora_amoebiformis
ABW97365.1_Amorphochlora_amoebiformis
ABW97364.1_Amorphochlora_amoebiformis
Rhizaria
EFN59821.1_Chlorella_variabilis
KIZ01259.1_Monoraphidium_neglectum
P08475.1_Chlamydomonas_reinhardtii
P00873.1_Chlamydomonas_reinhardtii
EEH51530.1_Micromonas_pusilla
CAL58650.1_Ostreococcus_tauri
CEG00799.1_Ostreococcus_tauri
CCO66446.1_Bathycoccus_prasinos
CCO66630.1_Bathycoccus_prasinos
CCO66738.1_Bathycoccus_prasinos
Chlorophyte
AncA
86
0.96
GBG87988.1_Chara_braunii
GBG63284.1_Chara_braunii
PNR54885.1_Physcomitrium_patens
PNR30214.1_Physcomitrium_patens
PNR33748.1_Physcomitrium_patens
PNR42621.1_Physcomitrium_patens
PNR43777.1_Physcomitrium_patens
PNR33800.1_Physcomitrium_patens
PNR39824.1_Physcomitrium_patens
PNR43798.1_Physcomitrium_patens
PNR44129.1_Physcomitrium_patens
PNR55254.1_Physcomitrium_patens
PNR59393.1_Physcomitrium_patens
PNR43528.1_Physcomitrium_patens
ERN18124.1_Amborella_trichopoda
EFJ07999.1_Selaginella_moellendorffii
EFJ16100.1_Selaginella_moellendorffii
QCI62510.1_Anthoceros_agrestis
QCI62511.1_Anthoceros_agrestis
QCI62513.1_Anthoceros_agrestis
QCI62517.1_Anthoceros_agrestis
PTQ29193.1_Marchantia_polymorpha
PTQ31243.1_Marchantia_polymorpha
PTQ31247.1_Marchantia_polymorpha
PTQ31246.1_Marchantia_polymorpha
PTQ31248.1_Marchantia_polymorpha
PTQ26327.1_Marchantia_polymorpha
PTQ31249.1_Marchantia_polymorpha
ERN03952.1_Amborella_trichopoda
KQJ81957.1_Brachypodium_distachyon
KQJ96661.1_Brachypodium_distachyon
KQJ86921.1_Brachypodium_distachyon
OAP15425.1_Arabidopsis_thaliana
OAO96413.1_Arabidopsis_thaliana
OAO94841.1_Arabidopsis_thaliana
Streptophyte
Form 1B
78 Felsenstein bootstrap value
0.97 Transfer bootstrap value

**Fig. S11. Constrained RbcS phylogeny**

Uncollapsed phylogeny of Rubisco's small subunit, based on the maximum-likelihood tree in fig. S12, but with minor topological constraints applied to match the constrained topology of the Rubisco large subunit tree as shown in fig. S9. Rubisco forms are labelled, with host species indicated for Form 1B Rubiscos. Felsenstein bootstrap values (black) and transfer bootstrap values (blue) are indicated next to nodes of interest. Scale bar: average substitution per site. Ancestral nodes (yellow) of AncE, AncD, AncD1, AncD2, AncD3, AncC, AncB, and AncA are indicated. NCBI Reference Sequence, UniprotKB, or IMG gene identifiers are listed.

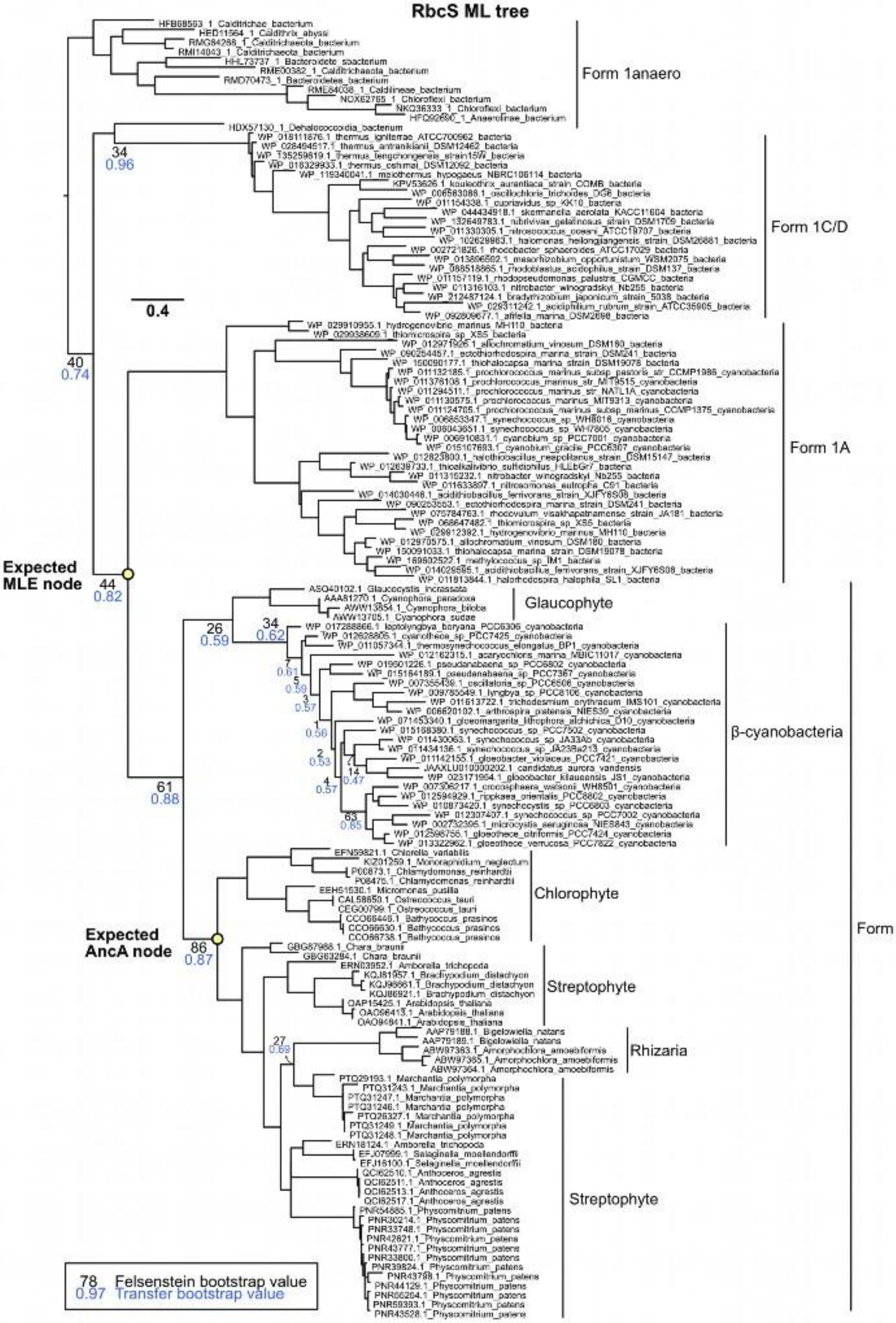
RbcS ML tree
Form 1anaero
Form 1C/D
0.4
Form 1A
Expected MLE node
Glaucophyte
β-cyanobacteria
Chlorophyte
Expected AncA node
Streptophyte
Rhizaria
Form 1B
Streptophyte
78 Felsenstein bootstrap value
0.97 Transfer bootstrap value

**Fig. S12. Maximum-likelihood RbcS phylogeny**

Uncollapsed maximum-likelihood phylogeny of Rubisco's small subunit. Rubisco forms are labelled, with host species indicated for Form 1B Rubiscos. Felsenstein bootstrap values (black) and transfer bootstrap values (blue) are indicated at selected nodes of interest. Scale bar: average substitution per site. Only the expected maximum-likelihood ancestral variants of AncE (MLE) and AncA is indicated, as incongruence between known species relationships (e.g. glaucophytes branching as the base of all β-cyanobacteria) and the maximum-likelihood RbcS topology obfuscates inference of ancestral nodes that best corresponds to either AncD/MLD or AncC/MLC. NCBI Reference Sequence, UniprotKB, or IMG gene identifiers are listed.

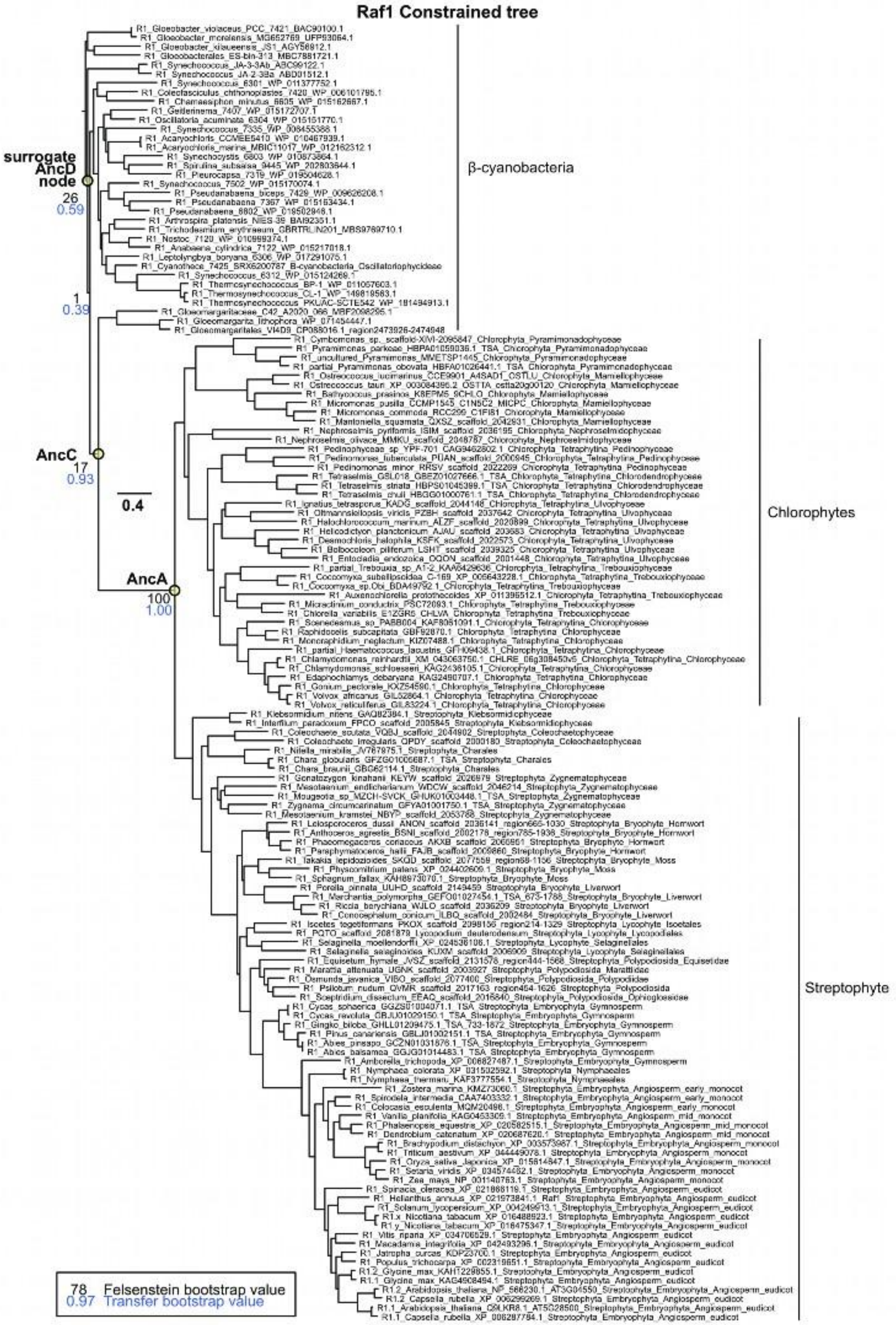

Raf1 Constrained tree
surrogate AncD node
26
0.59
1
0.39
AncC
17
0.93
0.4
AncA
100
1.00
β-cyanobacteria
Chlorophytes
Streptophyte
78 Felsenstein bootstrap value
0.97 Transfer bootstrap value

**Fig. S13. Constrained Raf1 phylogeny**

Uncollapsed phylogeny of Raf1 assembly chaperone, based on the maximum-likelihood tree in fig. S14, but with minor topological constraints within β-cyanobacteria applied to match the constrained topology of Rubisco's large subunit tree shown in fig. S9. Ancestral Raf1 nodes most likely to have co-existed within the same ancient organism as resurrected Rubisco ancestors are indicated with yellow circles. As AncD's Raf1 would correspond to the root of the tree, the succeeding ancestral node is utilized for AncD. Felsenstein bootstrap values (black) and transfer bootstrap values (blue) are indicated next to nodes of interest. Scale bar: average substitution per site. NCBI Reference Sequence, UniprotKB, or IMG gene identifiers are listed.

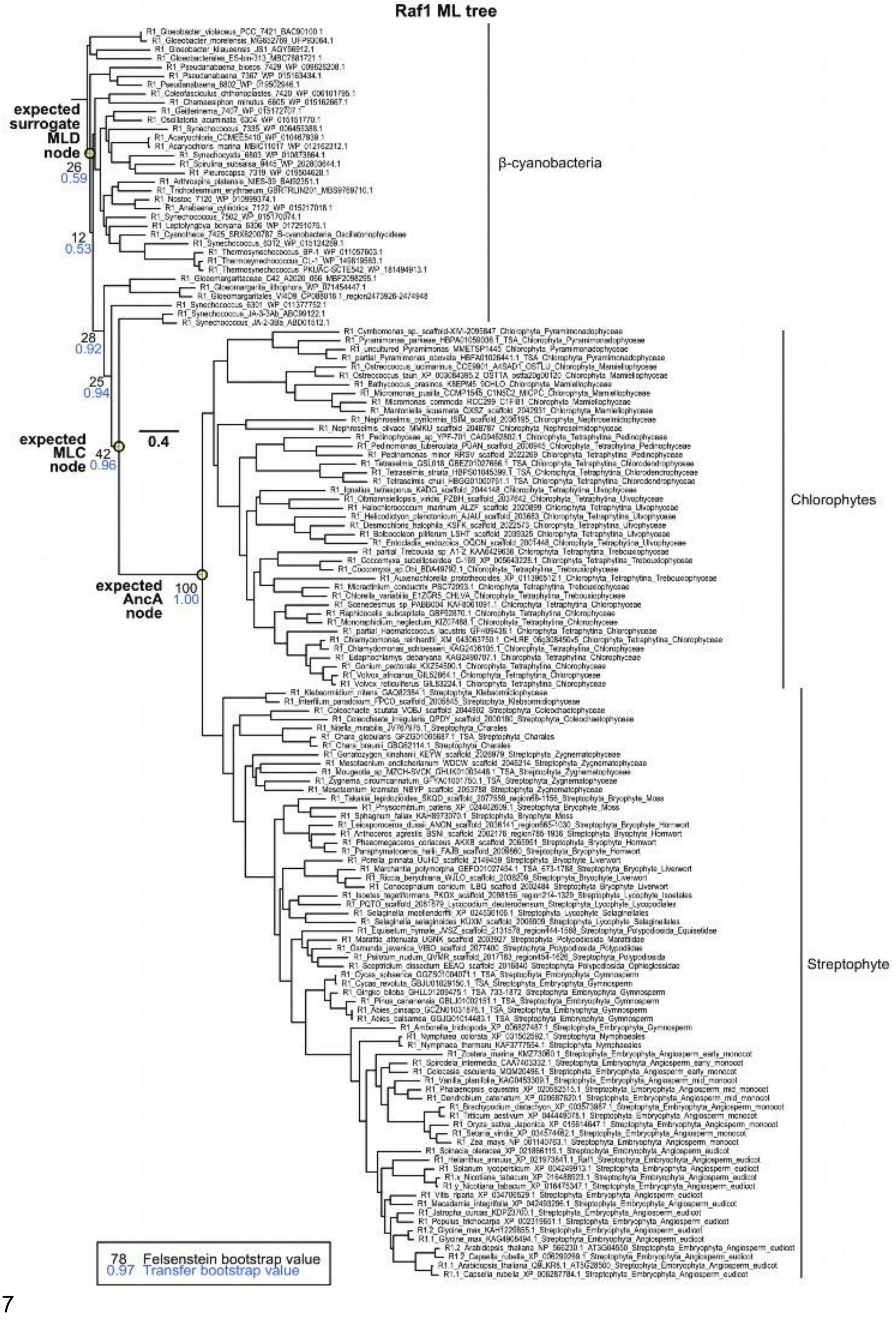
Raf1 ML tree
expected surrogate MLD node
26
0.59
12
0.53
28
0.92
25
0.94
expected MLC node
42
0.96
0.4
expected AncA node
100
1.00
β-cyanobacteria
Chlorophytes
Streptophyte
78 Felsenstein bootstrap value
0.97 Transfer bootstrap value

**Fig. S14. Maximum-likelihood Raf1 phylogeny**

Uncollapsed maximum-likelihood phylogeny of Raf1 rooted using the basal branching *Gloeobacter* clade as the outgroup. The ML topology is consistent with an early origin of plastids, with *Gloeomargaritales* and *Synechococcus sp. JA* being most closely related, though *Pseudanabaena* is observed to branch before this and not at the base of “late” diverging β-cyanobacterias. Expected ML ancestral nodes for Raf1 are indicated but are not examined, since chaperone dependence was due to changes in the LSU of Rubisco. Felsenstein bootstrap values (black) and transfer bootstrap values (blue) are indicated next to nodes of interest. Scale bar: average substitution per site. NCBI Reference Sequence, UniprotKB, or IMG gene identifiers are listed. Felsenstein bootstrap values (black) and transfer bootstrap values (blue) are indicated at selected nodes of interest. Scale bar: average substitution per site.

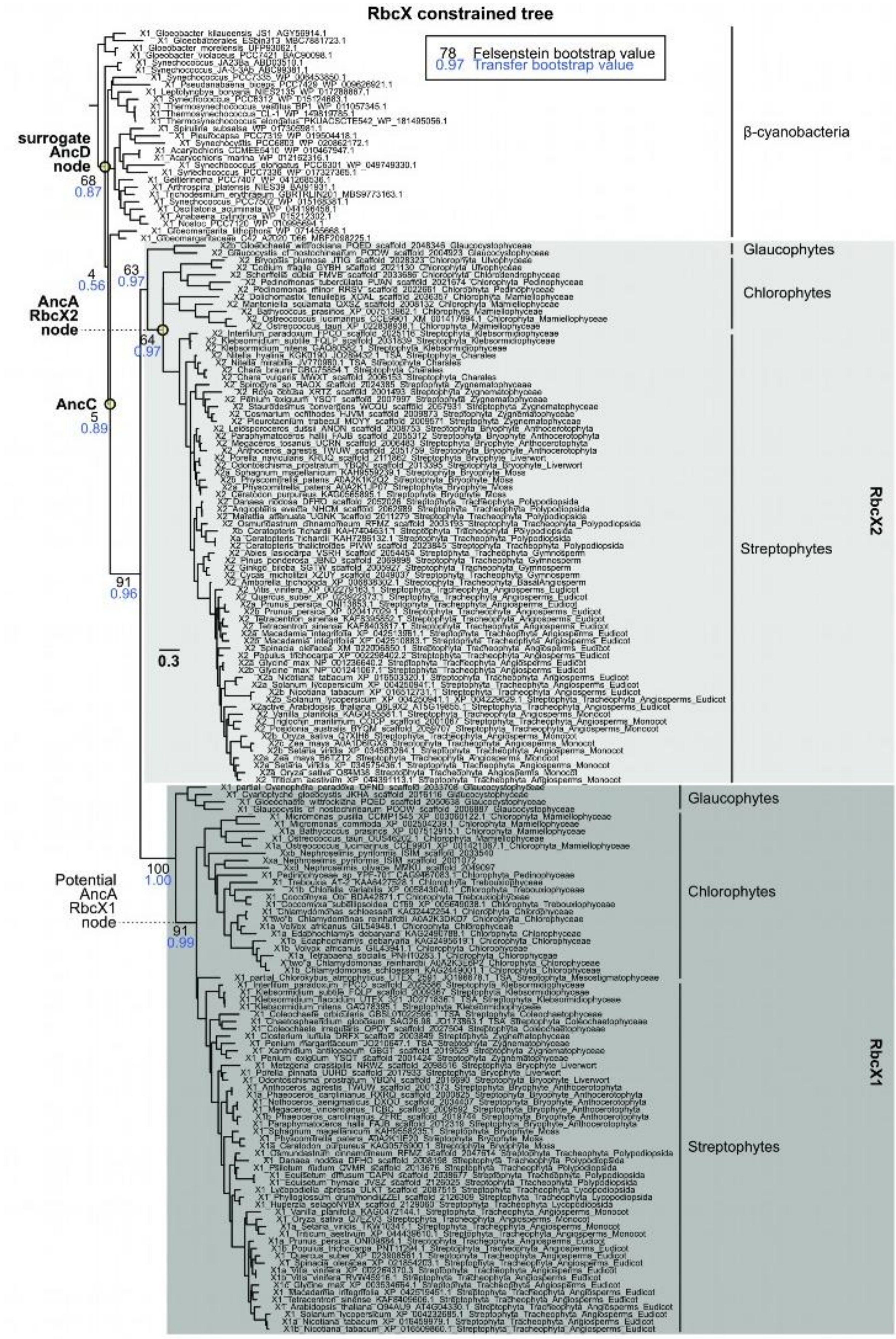
RbcX constrained tree
78 Felsenstein bootstrap value
0.97 Transfer bootstrap value
β-cyanobacteria
surrogate AncD node
68
0.87
Glaucophytes
Chlorophytes
4
0.56
63
0.97
AncA RbcX2 node
64
0.97
AncC
5
0.89
Streptophytes
RbcX2
91
0.96
0.3
Glaucophytes
Chlorophytes
100
1.00
Potential AncA RbcX1 node
91
0.99
Streptophytes
RbcX1

**Fig. S15. Constrained RbcX phylogeny**

Uncollapsed phylogeny of RbcX assembly chaperone, based on the maximum-likelihood tree in fig. S16, but with minor topological constraints within β-cyanobacteria applied to match the constrained topology of Rubisco's large subunit tree shown in fig. S9. The monophyly of green plastid RbcX paralogs remains unchanged and are labelled accordingly. Yellow circles denote ancestral RbcX nodes inferred to have coexisted with reconstructed Rubisco ancestors within the same ancient organism. For AncD's RbcX, which would correspond to the root, the next ancestral node is used as a surrogate. Node support is shown as Felsenstein bootstrap values (black) and transfer bootstrap values (blue). Scale bar: mean substitutions per site. Sequence identifiers from NCBI Reference Sequence, UniProtKB, or IMG are indicated for each tip.

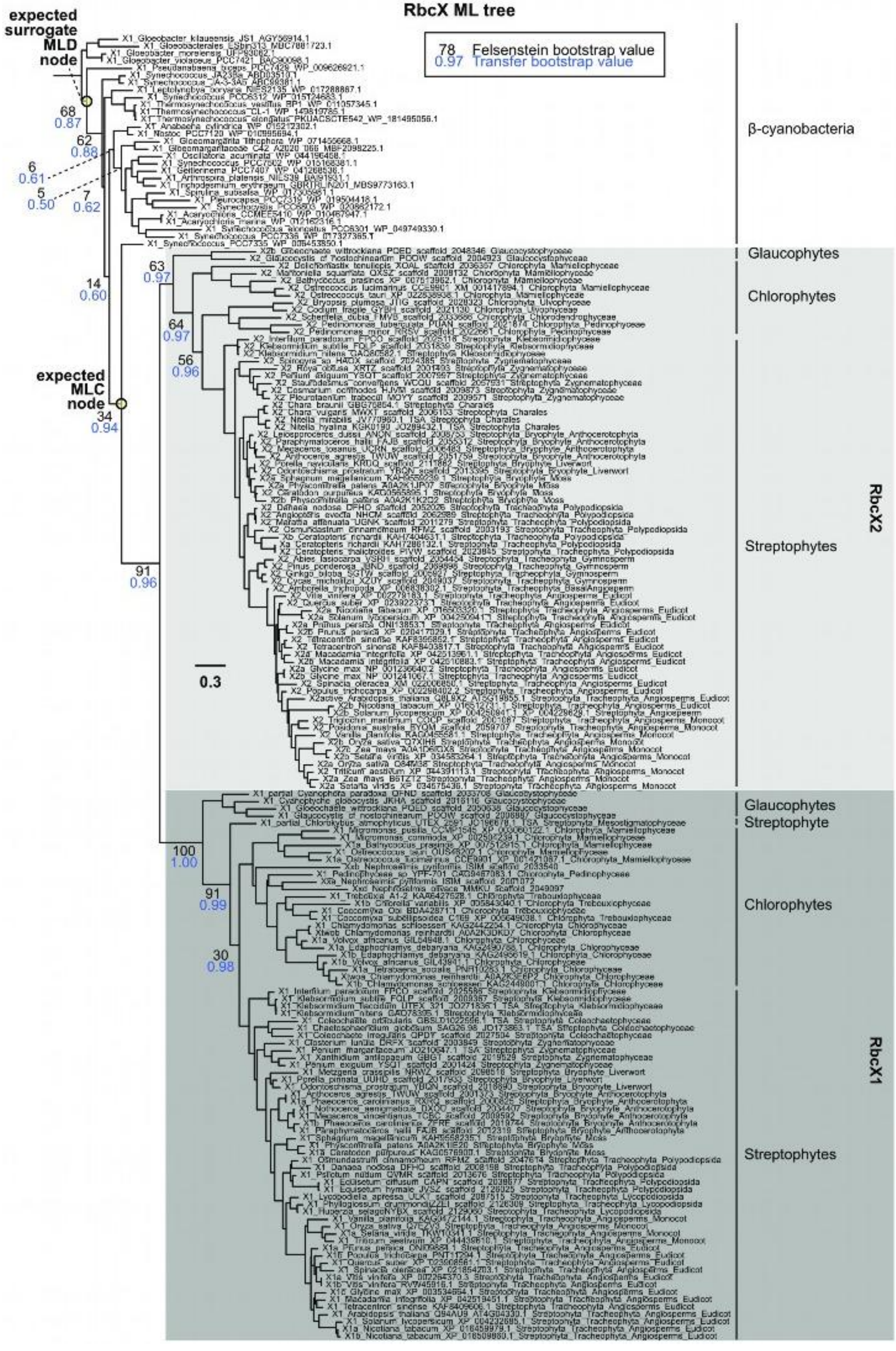
RbcX ML tree
78 Felsenstein bootstrap value
0.97 Transfer bootstrap value
expected surrogate MLD node
expected MLC node
β-cyanobacteria
Glaucophytes
Chlorophytes
Streptophytes
RbcX2
Glaucophytes
Streptophyte
Chlorophytes
Streptophytes
RbcX1
0.3

**Fig. S16. Maximum-likelihood RbcX phylogeny**

Uncollapsed maximum-likelihood phylogeny of RbcX rooted using the basal branching *Gloeobacter* clade as the outgroup. Monophyletic clades of green plastid RbcX paralogs are highlighted and labeled. Expected ML ancestral nodes for RbcX are indicated but not examined, since chaperone dependence is determined by changes in the LSU of Rubisco. Felsenstein bootstrap values (black) and transfer bootstrap values (blue) are indicated next to nodes of interest. Scale bar: average substitution per site. NCBI Reference Sequence, UniprotKB, or IMG gene identifiers are listed. Felsenstein bootstrap values (black) and transfer bootstrap values (blue) are indicated at selected nodes of interest. Scale bar: average substitution per site.

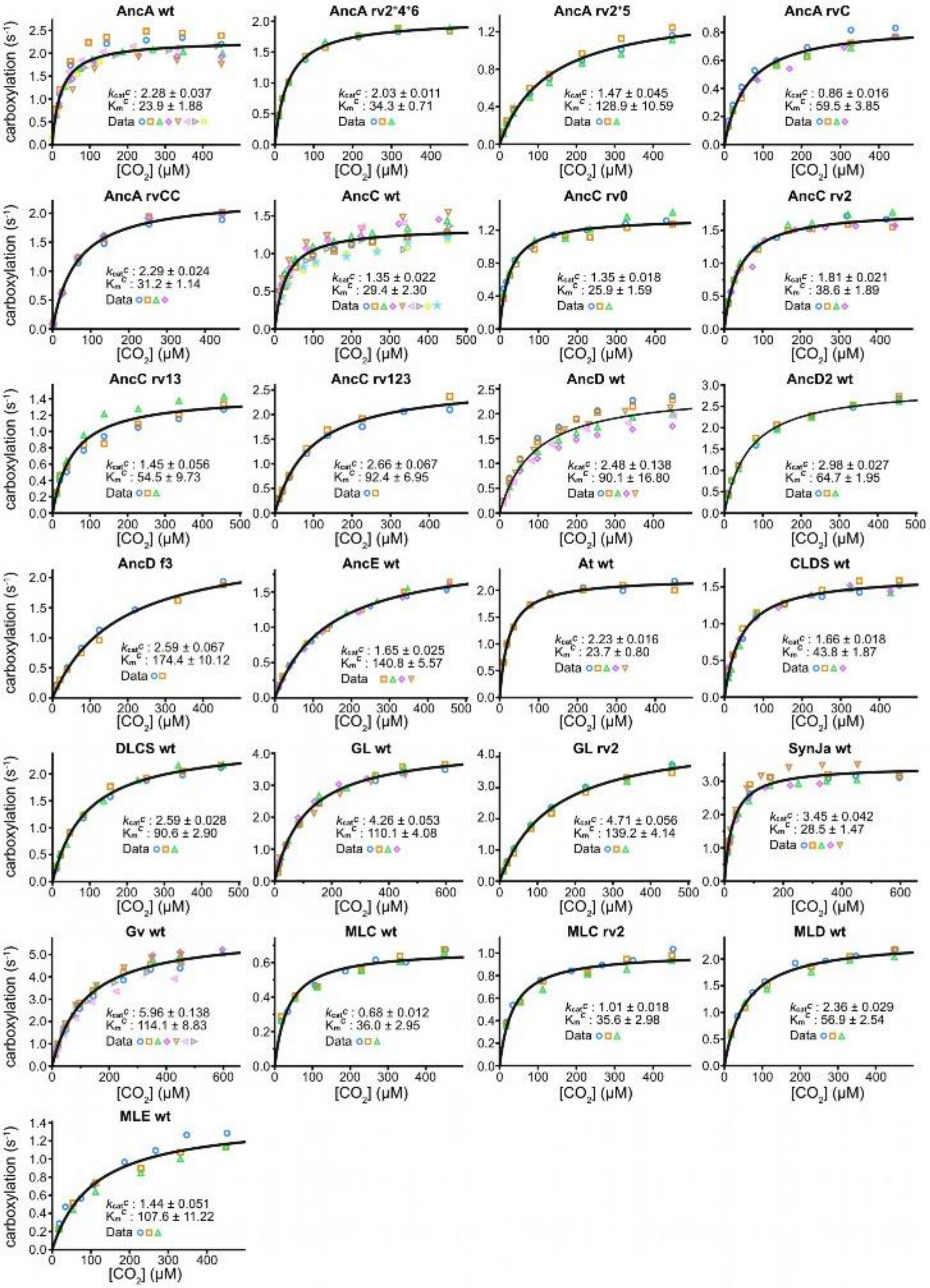
AncA wt
kcatc : 2.28 ± 0.037
Kmc : 23.9 ± 1.88
AncA rv2*4*6
kcatc : 2.03 ± 0.011
Kmc : 34.3 ± 0.71
AncA rv2*5
kcatc : 1.47 ± 0.045
Kmc : 128.9 ± 10.59
AncA rvC
kcatc : 0.86 ± 0.016
Kmc : 59.5 ± 3.85
AncA rvCC
kcatc : 2.29 ± 0.024
Kmc : 31.2 ± 1.14
AncC wt
kcatc : 1.35 ± 0.022
Kmc : 29.4 ± 2.30
AncC rv0
kcatc : 1.35 ± 0.018
Kmc : 25.9 ± 1.59
AncC rv2
kcatc : 1.81 ± 0.021
Kmc : 38.6 ± 1.89
AncC rv13
kcatc : 1.45 ± 0.056
Kmc : 54.5 ± 9.73
AncC rv123
kcatc : 2.66 ± 0.067
Kmc : 92.4 ± 6.95
AncD wt
kcatc : 2.48 ± 0.138
Kmc : 90.1 ± 16.80
AncD2 wt
kcatc : 2.98 ± 0.027
Kmc : 64.7 ± 1.95
AncD f3
kcatc : 2.59 ± 0.067
Kmc : 174.4 ± 10.12
AncE wt
kcatc : 1.65 ± 0.025
Kmc : 140.8 ± 5.57
At wt
kcatc : 2.23 ± 0.016
Kmc : 23.7 ± 0.80
CLDS wt
kcatc : 1.66 ± 0.018
Kmc : 43.8 ± 1.87
DLCS wt
kcatc : 2.59 ± 0.028
Kmc : 90.6 ± 2.90
GL wt
kcatc : 4.26 ± 0.053
Kmc : 110.1 ± 4.08
GL rv2
kcatc : 4.71 ± 0.056
Kmc : 139.2 ± 4.14
SynJa wt
kcatc : 3.45 ± 0.042
Kmc : 28.5 ± 1.47
Gv wt
kcatc : 5.96 ± 0.138
Kmc : 114.1 ± 8.83
MLC wt
kcatc : 0.68 ± 0.012
Kmc : 36.0 ± 2.95
MLC rv2
kcatc : 1.01 ± 0.018
Kmc : 35.6 ± 2.98
MLD wt
kcatc : 2.36 ± 0.029
Kmc : 56.9 ± 2.54
MLE wt
kcatc : 1.44 ± 0.051
Kmc : 107.6 ± 11.22
Data
carboxylation (s-1)
[CO2] (μM)

**Fig. S17. Rubisco kinetic measurements and Michaelis-Menten fits.**

Carboxylation kinetics of Rubiscos shown in fig. S1 and various point plots for comparisons of catalytic activity utilized in this study. Substrate $CO_2$ concentrations are listed in each plot, while a fixed saturating concentration of the co-substrate RuBP at approximately 1.08 mM was utilized for all assays. Global Michaelis-Menten curve fits are depicted in black, with individual replicate sets indicated by distinct colours and symbols as indicated. Determined catalytic parameters are shown in each plot with SEs of the global fit. $k_{cat}^{C}$ = maximal rate of carboxylation under saturating substrate concentrations; $K_m^{CO2}$ = Michaelis constant for $CO_2$. *At*, *Arabidopsis thaliana*; *Gv*, *Gloeobacter violaceus*; *SynJA*, *Synechococcus sp. JA-2-3B'a (2-13)*; *Gl*, *Gloeomargarita lithophora*; DLCS, AncD's RbcL with AncC's RbcS; CLDS, AncC's RbcL with AncD's RbcS; wt, wild type.

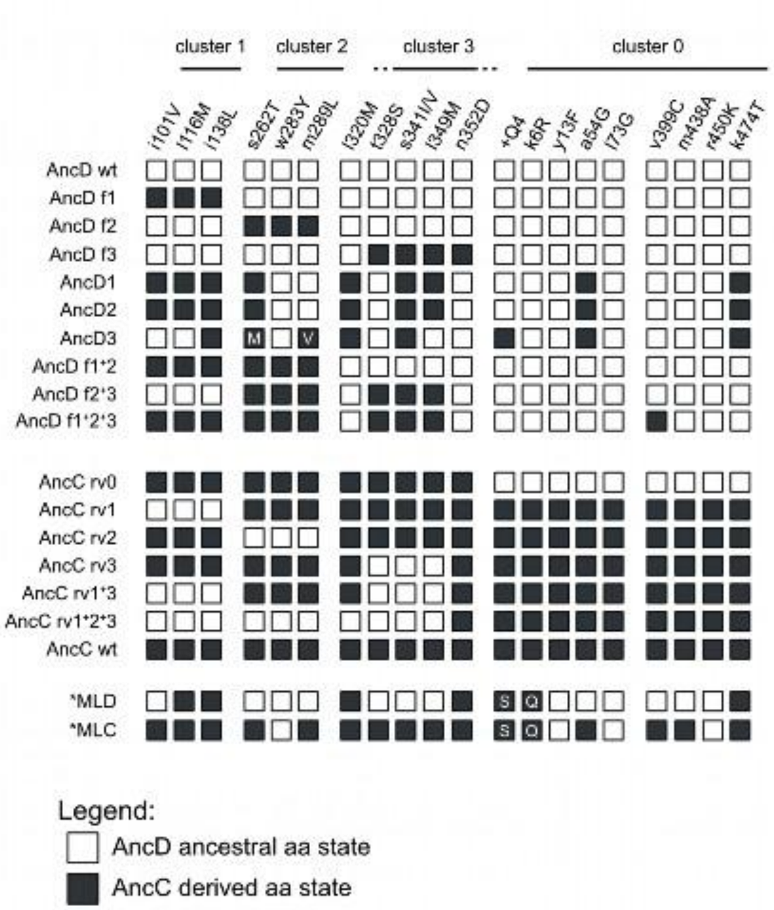

**Fig. S18. Identities of amino-acid states within mutant variants tested between AncD and AncC.**

Simple schematic list of all 20 historical substitutions and the presence of ancestral or derived substitutions in all tested mutant variants shown in Fig. 3B-C and S3B-C. Mutants containing amino-acid states corresponding to AncD are indicated in white, while derived AncC amino acid states are kept black. Substitutions are grouped into clusters as described at the top. For each historical substitution site, small alphabets denote ancestral states that correspond to the color of their respective squares, while capitalized letters denote derived states. Sites containing derived amino-acid states that differ from the derived amino-acid states of AncC or MLC are kept black but modified with their respective amino-acid identities in white capitalized alphabets.

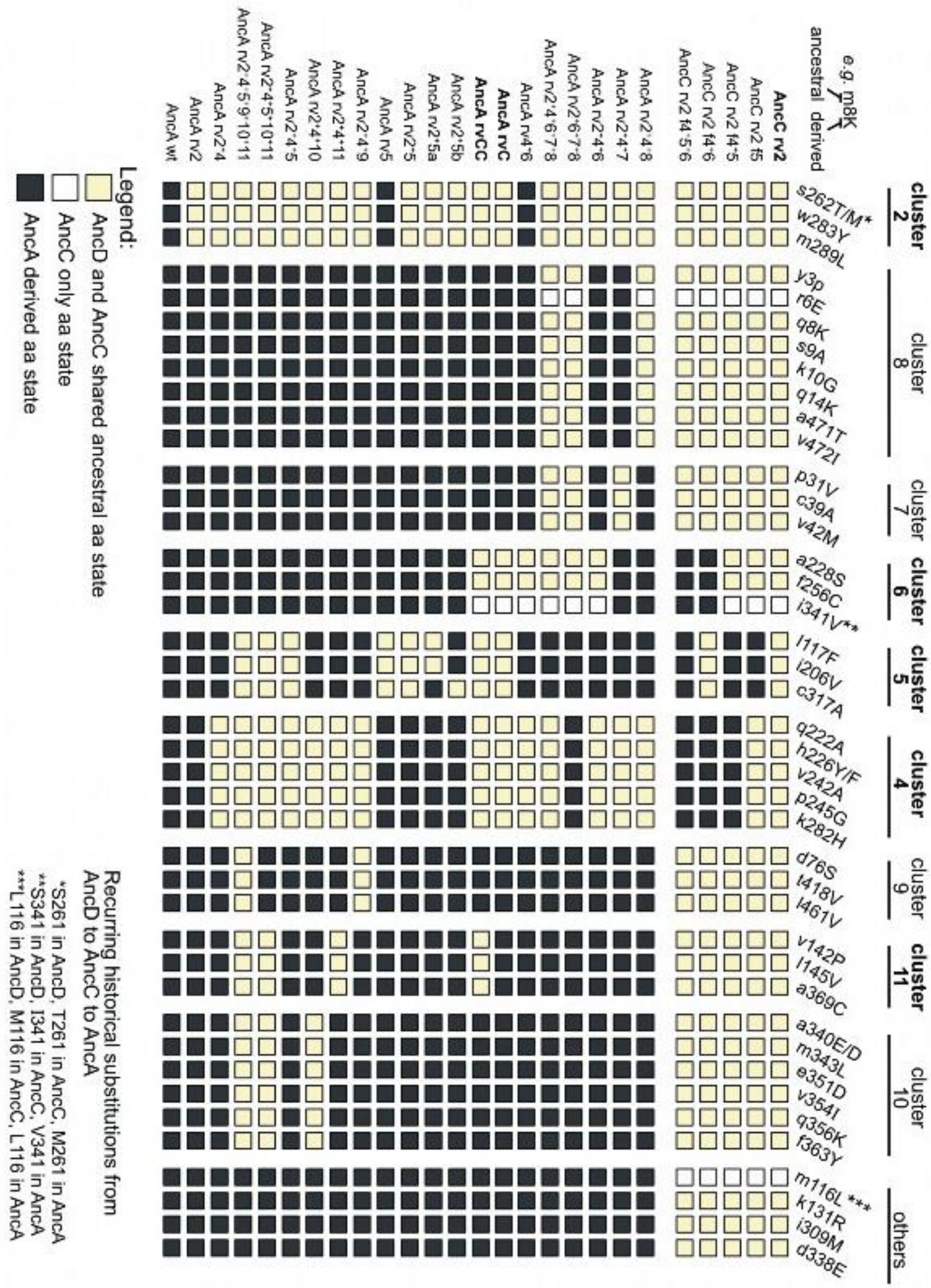

**Fig. S19. Identities of amino-acid states within mutant variants tested between AncC and AncA.**

Schematic list of all 39 historical substitutions and the presence of ancestral or derived states in tested mutant variants shown in Fig. 4B-C, S5E-F, and S5H. Since cluster 2 reversions are also present depending on the variant, amino acid states corresponding to AncD states are coloured yellow. Mutants containing historical substitutions that underwent another change between AncD to AncC are shown as white, while derived AncA amino acid states are kept black. Substitutions are grouped into clusters indicated at the top. For each historical substitution site, small alphabets denote ancestral states that correspond to the color of their respective squares, while capitalized letters denote derived states. Sites experiencing recurring historical substitutions are further denoted by an asterisk (*) and described at the bottom, while sites with variable amino-acid states across different ancestors are modified with a forward slash (/).

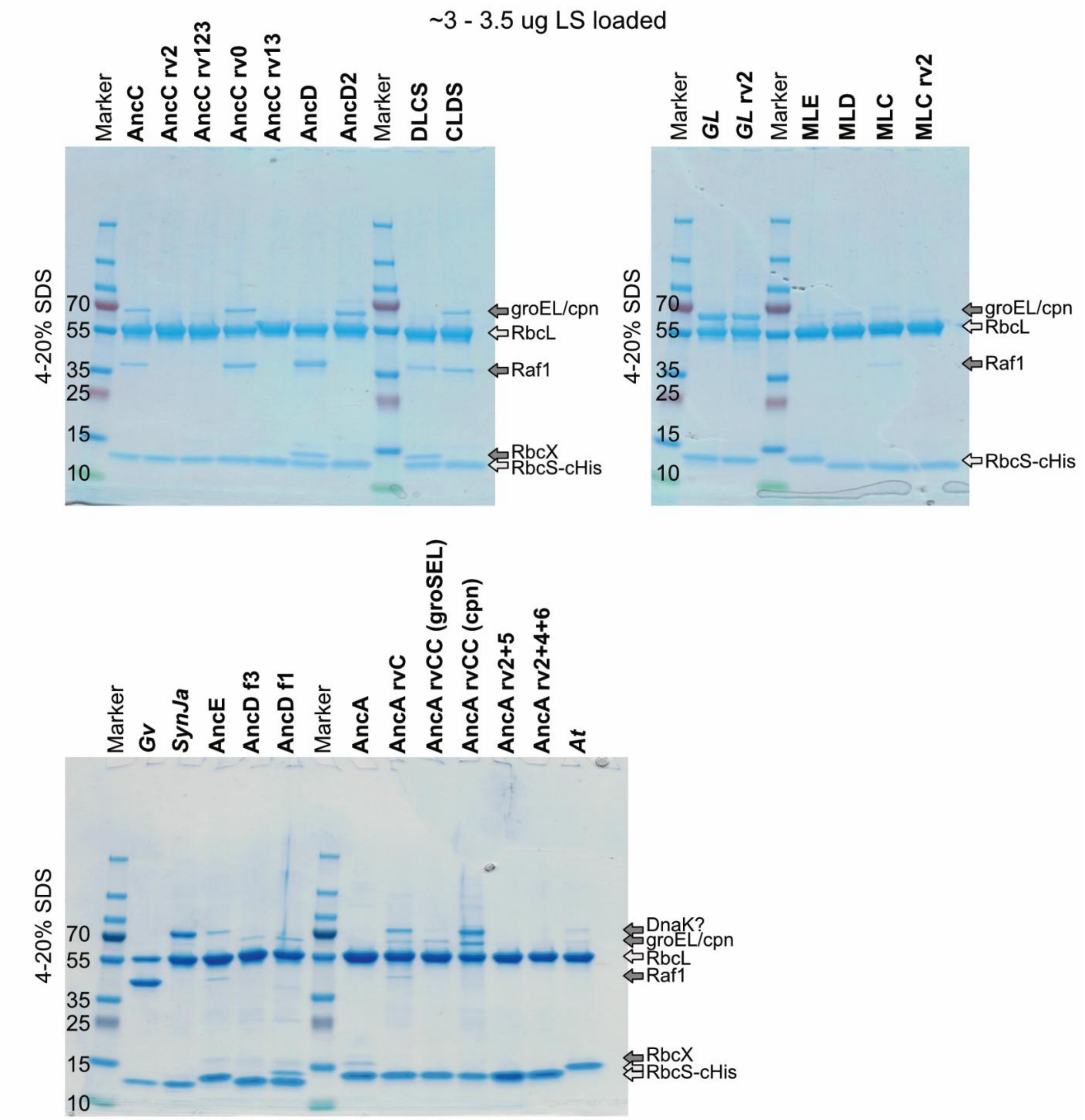

**Fig. S20. Assessment of purified Rubisco enzymes**

SDS-PAGE analysis of purified ancestral and extant wild-type or variant Rubiscos utilized in this study as indicated. Approximately 3.0 – 3.5 µg Rubisco active sites (defined as one LSU and SSU) determined by A280 nm quantification were loaded in each lane. The presence of co-purified assembly chaperones is inferred based on their expected molecular weights. Chaperones corresponding to their cognate Rubiscos or time-matched ancestors were utilized accordingly for purification when required (see Methods). All Rubisco variants were purified via IMAC followed by SEC, with exception of AncD f1 which was directly utilized following IMAC and desalting.

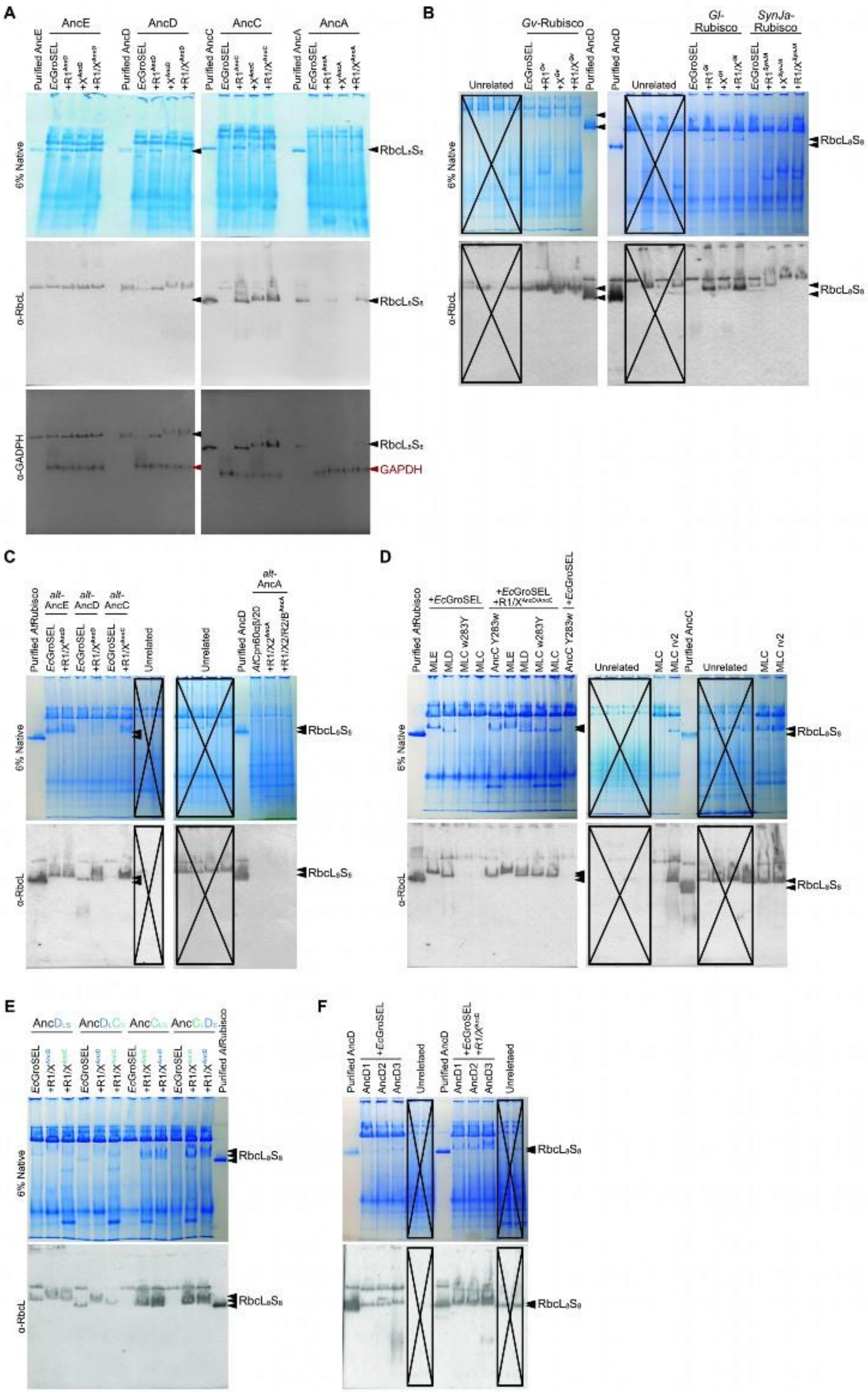
A
B
C
D
E
F
AncE
AncD
AncC
AncA
6% Native
α-RbcL
α-GADPH
RbcL8S8
GAPDH
Gv-Rubisco
Gl-Rubisco
SynJa-Rubisco
Unrelated
Purified AncD
Purified AtRubisco
alt-AncE
alt-AncD
alt-AncC
alt-AncA
+EcGroSEL
MLE
MLD
MLC
MLC rv2
Purified AncC
AncD1
AncD2
AncD3

741

**Fig. S21. Full gels and blots with controls of Fig. 1C, S2B-D, 2E, S3G, and S3I.**

Native-PAGE and western blot analysis of soluble lysate extracts obtained from heterologous expression of indicated chaperonins, Rubisco variants, and assembly chaperones in *E. coli*. Full gels and blots examining the assembly characteristics of **(A)** ancestral Rubisco variants from the constrained topology utilized throughout the study, **(B)** extant Rubisco enzymes from selected key cyanobacteria species based on the known relationships of β-cyanobacteria, **(C)** alternative ancestral Rubiscos from the constrained topology containing a less likely ancestral variant of the large subunit, **(D)** Rubisco ancestors from the unconstrained maximum-likelihood topology that would require many HGT-events to explain the branching order, **(E)** hybrid Rubisco ancestors containing a reciprocal switch of large and small subunits from AncD and AncC, and **(F)** intermediate AncD ancestors that occur on the path towards AncC on the constrained topology. Approximately 40 µg of sample soluble protein was loaded for each lane based on BCA standard curve estimation, or 1-2 µg of purified protein based on A280nm measurements. For (A), a sequential anti-GAPDH immunoblot was performed on the same membrane following anti-RbcL detection, without stripping. GAPDH signals (red) were observed as distinct bands that did not overlap with the previously detected RbcL bands. Expected assembled Rubisco bands are indicated by arrows, with multiple arrows shown when differing migration patterns from distinct assembled Rubiscos of different species are observed.

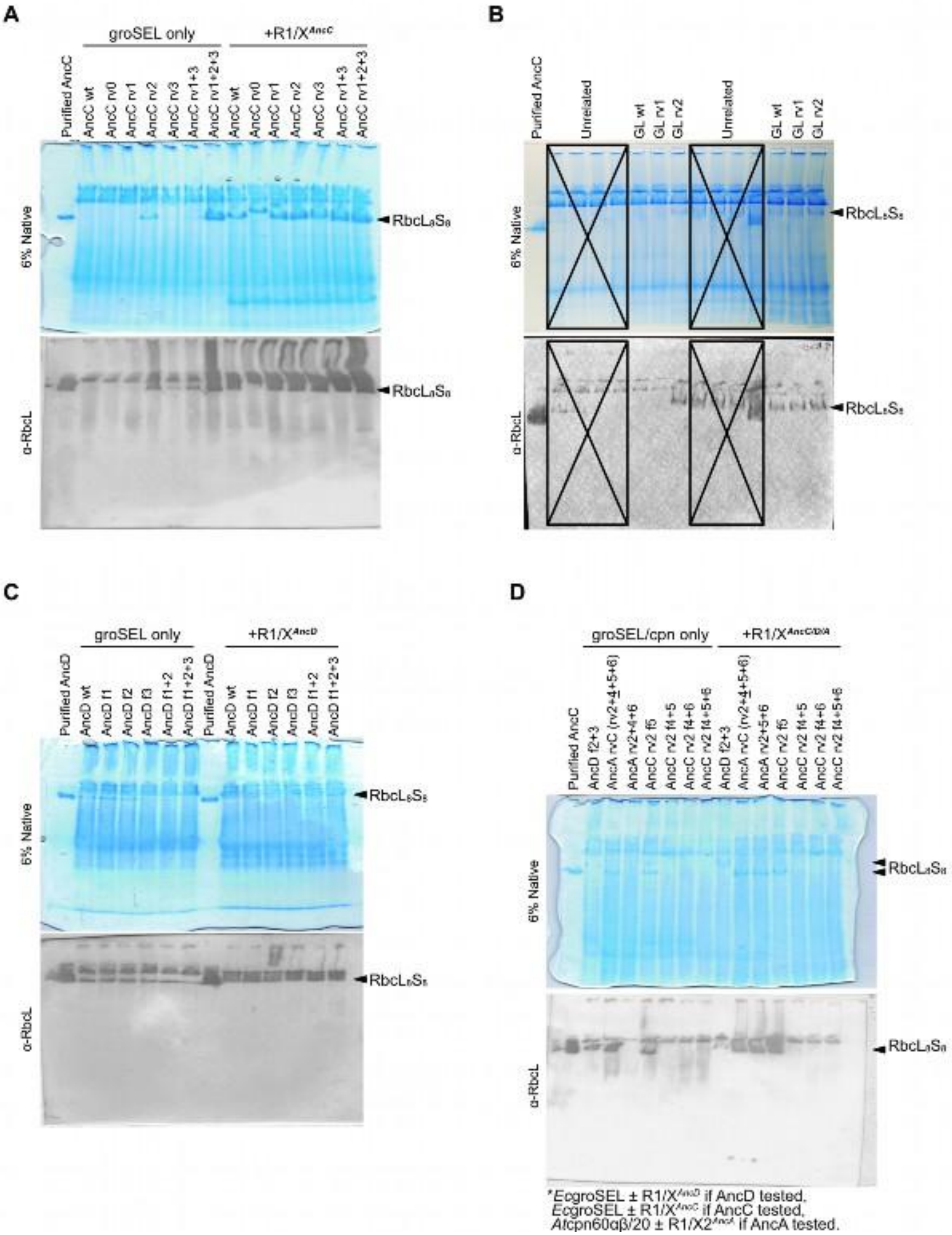

**Fig. S22. Full gels and blots with controls of Fig. 3B, 3E, S4B, and S5H.**

Native-PAGE and western blot analysis of soluble lysate extracts obtained from heterologous expression of indicated chaperonins, Rubisco variants, and assembly chaperones in *E. coli*. Full gels and blots examining the assembly characteristics of **(A)** AncC reversion variants, **(B)** extant wild type and reverted Rubisco variants from the βcyanobacterium *G. lithophora*, **(C)** AncD variants containing derived amino-acid states of indicated clusters, and **(D)** AncC variants containing reverted cluster 2 states that rescues independent assembly, combined with derived amino acid states from indicated clusters that deepens the chaperone entrenching phenotype occurring between AncC and AncA. Expected assembled Rubisco bands are indicated by arrows, with multiple arrows shown when differing migration patterns from distinct assembled Rubiscos of different species are observed. Approximately 40 µg of sample soluble protein was loaded for each lane based on BCA standard curve estimation, or 1-2 µg of purified protein based on A280nm measurements.

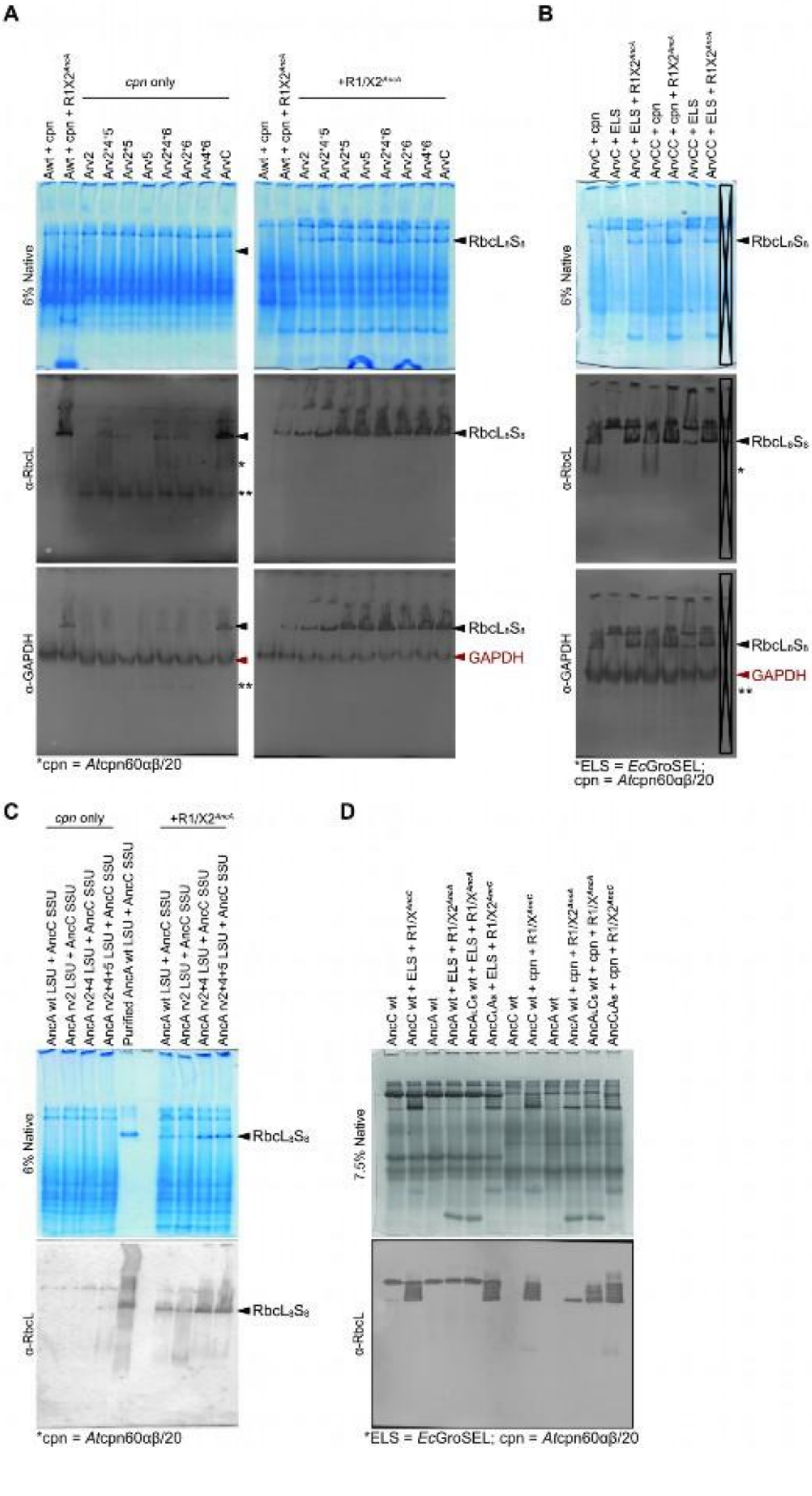
A
B
C
D
cpn only
+R1/X2AncA
6% Native
α-RbcL
α-GAPDH
RbcL8S8
GAPDH
*cpn = Atcpn60αβ/20
*ELS = EcGroSEL;
cpn = Atcpn60αβ/20
7.5% Native
*ELS = EcGroSEL; cpn = Atcpn60αβ/20

777

**Fig. S23. Full gels and blots with controls of Fig. 4B-C, S5F, and S5C-D.**

Native-PAGE and western blot analysis of soluble lysate extracts obtained from heterologous expression of indicated chaperonins, Rubisco variants, and assembly chaperones in *E. coli*. Full gels and blots examining the assembly characteristics of **(A-B)** AncA chaperone or chaperonin reversion variants with growth and co-expression conducted at 28°C as compared to 30°C in fig. S23A, hybrid Rubiscos **(C)** containing wild type or reverted variant AncA RbcL and wild type AncC RbcS, or **(D)** reciprocal switches of AncA RbcL and AncC RbcS subunits with co-expression of their corresponding type chaperonins or assembly chaperones as indicated. For (A-B), a sequential anti-GAPDH immunoblot was performed on the same membrane following anti-RbcL detection, without stripping. GAPDH signals (red) were observed as distinct bands that mostly do not overlap with the previously detected RbcL bands, except for a lower migrating band (*) that mostly occurs when reverting chaperone/ chaperonin entrenching substitutions or their assembly competent subsets in the presence of plastidal chaperonins. Together with another distinct lower migrating band (**), these bands may represent chaperonin-bound RbcL complexes of differing lower stoichiometries. Expected assembled Rubisco bands are indicated by arrows, with multiple arrows shown when differing migration patterns from distinct assembled Rubiscos of different species are observed. Approximately 40 µg of sample soluble protein was loaded for each lane based on BCA standard curve estimation, or 1-2 µg of purified protein based on A280nm measurements.

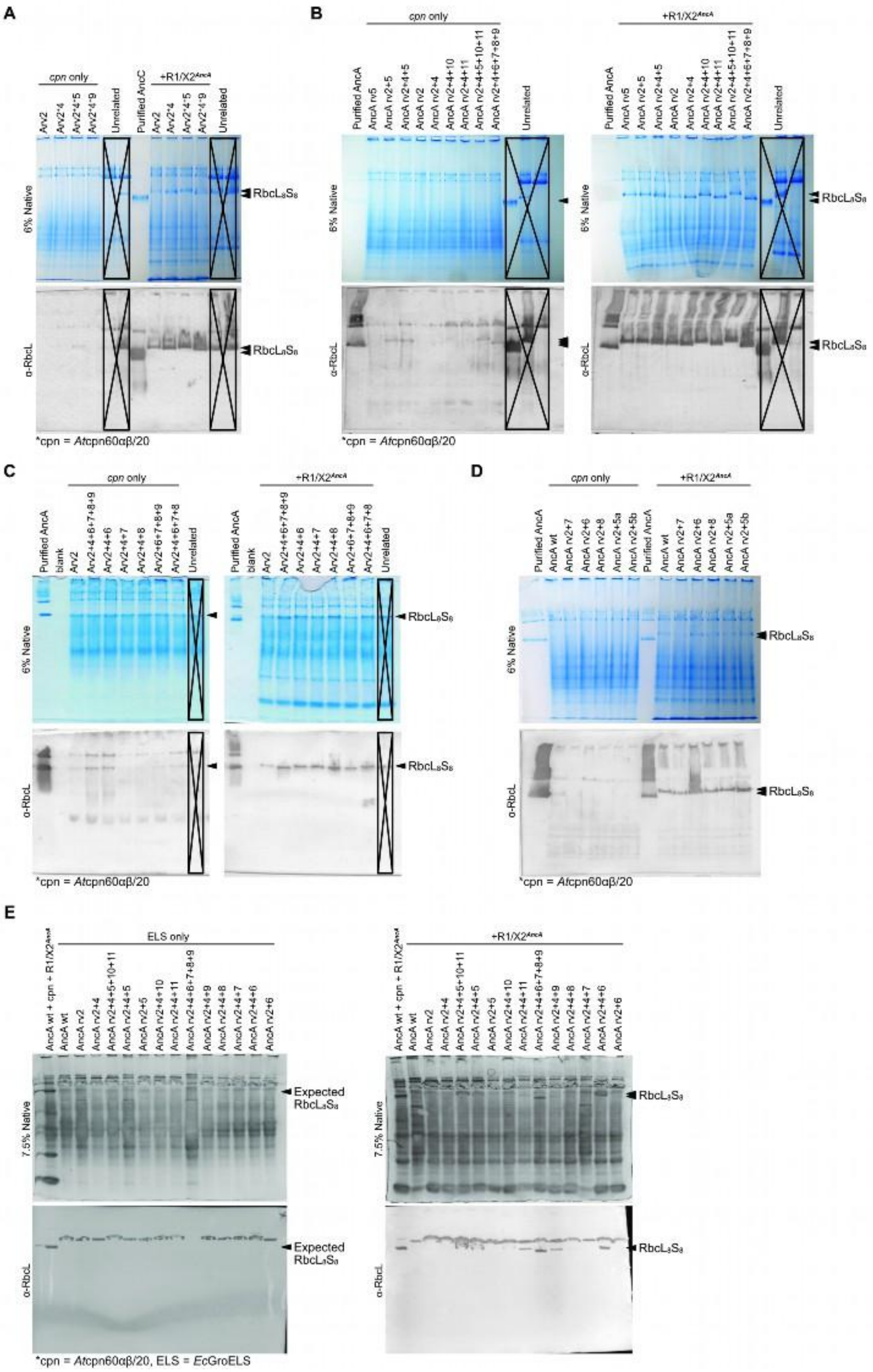
A
B
C
D
E
cpn only
+R1/X2^AncA
6% Native
7.5% Native
α-RbcL
RbcL8S8
Expected RbcL8S8
ELS only
Purified AncA
Purified AncC
Unrelated
blank
*cpn = Atcpn60αβ/20
*cpn = Atcpn60αβ/20, ELS = EcGroELS

799

**Fig. S24. Full gels and blots with controls for Fig. S5E and S5G.**

Native-PAGE and western blot analysis of soluble lysate extracts obtained from heterologous expression of indicated chaperonins, Rubisco variants, and assembly chaperones in *E. coli*. Full gels and blots conducted as screens to identify casual genetic determinants that further entrench green plastid type Rubisco with chaperones, based on the assembly characteristics of **(A-D)** AncA variants containing reversions of various combinations of historical substitution clusters that occur between AncC and AncA, and **(E)** selected AncA assembly competent reversion combinations co-produced with prokaryotic GroELS chaperonins instead of the compositionally more complex plastidal chaperonins, with or without assembly chaperones. Approximately 40 µg of sample soluble protein was loaded for each lane based on BCA standard curve estimation, or 1-2 µg of purified protein based on A280nm measurements. Expected assembled Rubisco bands are indicated by arrows, with multiple arrows shown when differing migration patterns from distinct assembled Rubiscos of different species are observed.

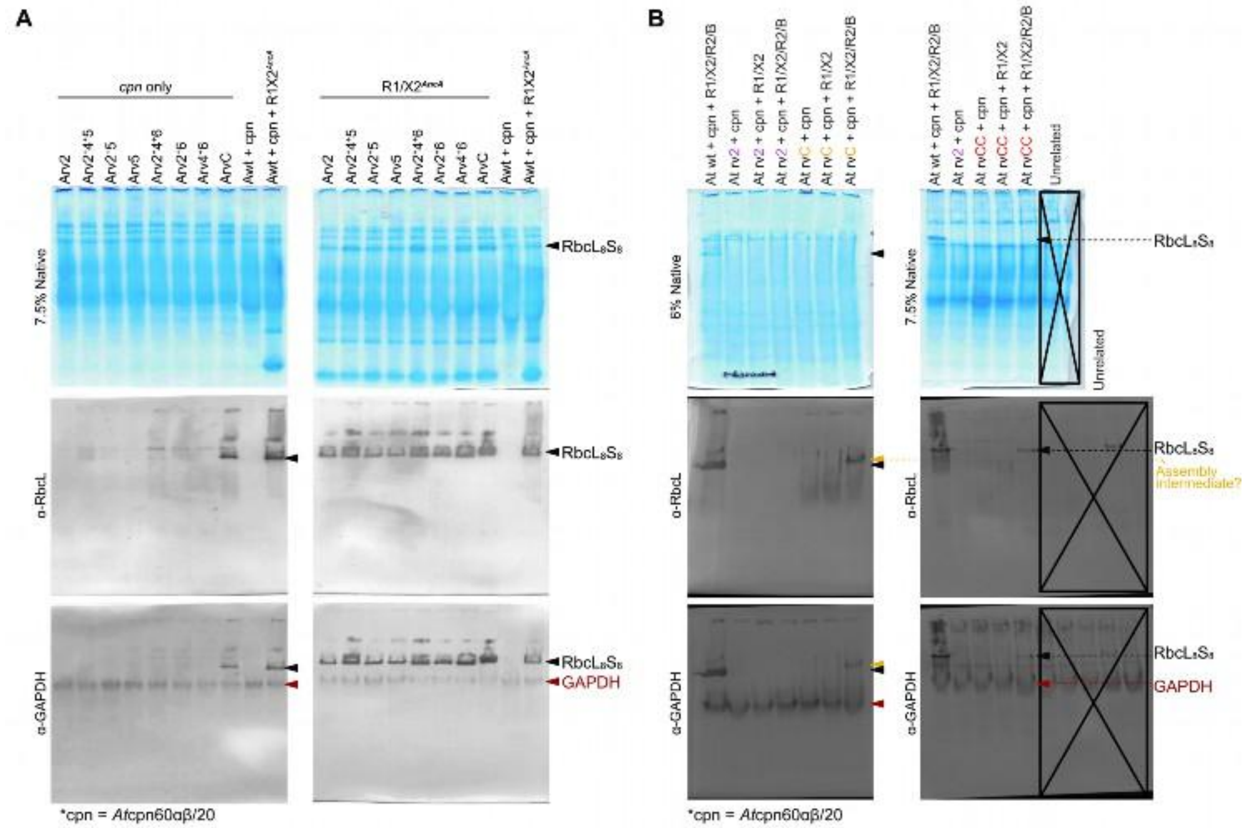

**Fig. S25. Full gels and blots with controls for Fig. S6B and S6D.**

Native-PAGE and western blot analysis of soluble lysate extracts obtained from heterologous expression of indicated chaperonins, Rubisco variants, and assembly chaperones in *E. coli*. Full gels and blots examining the assembly characteristics of **(A)** AncA chaperone reversion variants as tested in fig. S23A but with growth and co-expression carried out at 30°C instead of 28°C, resulting in abolishment of weak rescue from chaperone dependent assembly for subset combinations of AncA rvC. **(B)** A similar Native-PAGE and western blot analysis of soluble lysate fractions extracted from *E.coli* cells co-expressing *A. thaliana* (*At*) Rubisco wild type and reverted variants under indicated combinations of its cognate chaperonin system and assembly chaperones. For both gel and blot sets, a sequential anti-GAPDH immunoblot was performed on the same blotted membrane following anti-RbcL detection, without treatment with stripping agents. GAPDH signals (red) were observed as distinct bands that mostly do not overlap with the detected RbcL bands corresponding to assembled Rubisco complexes.

**Table S1. Kinetic parameters of ancestral and extant Rubisco variants at 25°C.**

All Rubiscos were purified with their respective small subunits and minimal chaperone requirements when possible unless indicated (also see Methods section). AncA rvC contains reversions of clusters 2, 4, 5, and 6 to their ancestral states and rescues chaperone dependent assembly when plastidal chaperonins are utilized. AncA rvCC contains reversion of clusters as per AncA rvC with the addition of cluster 11 reversion, restoring wild type catalytic activities and self-assembly even when only prokaryotic chaperonins are provided. *A. thaliana*, *Arabidopsis thaliana*; *G. violaceus*, *Gloeobacter violaceus*; *SynJA*, *Synechococcus sp. JA-2-3B'a (2-13)*; *G. lithophora*, *Gloeomargarita lithophora*; DLCS, AncD's RbcL with AncC's RbcS; CLDS, AncC's RbcL with AncD's RbcS. $k_{cat}^{C}$ = maximal rate of carboxylation under saturating substrate concentrations; $K_m^{CO2}$ = Michaelis constant for $CO_2$; $k_{cat}^{C}/K_m^{CO2}$ = derived carboxylation efficiencies based on determined $k_{cat}^{C}$ and $K_m^{CO2}$ values, $S_{C/O}$: determined specificity values for carboxylation over oxygenation reactions, where $S_{C/O} = [k_{cat}^{C}/K_m^{CO2}]/[k_{cat}^{O}/K_m^{O2}]$. Full kinetic curves are shown in fig. S15. Values are global fits with SEs of kinetic curves for $k_{cat}^{C}$, $K_m^{CO2}$, and $k_{cat}^{C}/K_m^{CO2}$, or means with SDs of carboxylation to oxygenation ratios for $S_{C/O}$, with the number of replicates (N) indicated in parentheses. A corresponding SDS-PAGE analysis of Rubiscos characterized is shown in fig. S20. n.d., not determined.

| **Rubisco name** | $k_{cat}^{C}$ ($s^{-1}$) | $K_m^{CO2}$ (μM) | $k_{cat}^{C}/K_m^{CO2}$ ($M^{-1}\ s^{-1}$) | **Specificity** ($S_{C/O}$) |
|---|---|---|---|---|
| *A. thaliana* (*At*) | 2.23 ± 0.02 (2) | 23.74 ± 0.80 (2) | 93.86 ± 3.25 (2) | 79.11 ± 1.05 (4) |
| AncE | 1.65 ± 0.03 (4) | 140.78 ± 5.57 (4) | 11.72 ± 0.50 (4) | 42.36 ± 0.97 (4) |
| AncD | 2.48 ± 0.14 (6) | 90.08 ± 16.80 (6) | 27.51 ± 5.35 (6) | 45.28 ± 0.52 (7) |
| AncD2 | 2.98 ± 0.03 (3) | 64.67 ± 1.95 (3) | 46.08 ± 1.45 (3) | 52.91 ± 0.65 (3) |
| AncD f1 | n.d. | n.d. | n.d. | 50.70 ± 1.06 (4) |
| AncD f3 | 2.59 ± 0.07 (2) | 174.45 ± 10.12 (2) | 14.85 ± 0.94 (2) | 52.73 ± 1.11 (4) |
| DLCS | 2.59 ± 0.03 (3) | 90.61 ± 2.90 (3) | 28.58 ± 0.97 (3) | 46.65 ± 0.60 (4) |
| CLDS | 1.66 ± 0.02 (4) | 43.76 ± 1.87 (4) | 37.99 ± 1.68 (4) | 63.09 ± 1.20 (4) |
| *G. violaceus* (*Gv*) | 5.96 ± 0.14 (7) | 114.13 ± 8.83 (7) | 52.22 ± 4.22 (7) | 60.01 ± 1.30 (3) |
| *Syn. JA-2-3B'a* (*SynJA*) | 3.45 ± 0.04 (5) | 28.53 ± 1.47 (5) | 121.02 ± 6.43 (5) | 57.46 ± 1.78 (3) |
| *G. lithophora* (*Gl*) | 4.26 ± 0.05 (5) | 110.13 ± 4.08 (5) | 38.67 ± 1.51 (5) | 66.07 ± 1.22 (4) |
| *Gl* rv2 | 4.71 ± 0.06 (3) | 139.17 ± 4.14 (3) | 33.88 ± 1.08 (3) | 67.45 ± 0.87 (4) |
| AncC | 1.35 ± 0.02 (9) | 29.36 ± 2.30 (9) | 45.98 ± 3.68 (9) | 60.30 ± 1.19 (4) |
| AncC rv2 | 1.81 ± 0.02 (4) | 38.63 ± 1.89 (4) | 46.85 ± 2.36 (4) | 64.46 ± 0.52 (4) |
| AncC rv0 | 1.35 ± 0.02 (3) | 25.91 ± 1.59 (3) | 52.25 ± 3.28 (3) | 60.01 ± 1.93 (2) |
| AncC rv123 | 2.66 ± 0.07 (2) | 92.37 ± 6.95 (2) | 28.78 ± 2.28 (2) | 50.83 ± 0.96 (2) |
| AncC rv13 | 1.45 ± 0.06 (3) | 54.49 ± 9.73 (3) | 26.55 ± 4.85 (3) | n.d. |
| AncA | 2.28 ± 0.04 (8) | 23.95 ± 1.88 (8) | 95.31 ± 7.64 (8) | 71.21 ± 0.77 (10) |
| AncA rvC | 0.86 ± 0.02 (4) | 59.51 ± 3.85 (4) | 14.38 ± 0.97 (4) | 65.05 ± 1.03 (3) |
| AncA rvCC | 2.29 ± 0.02 (4) | 31.22 ± 1.14 (4) | 73.48 ± 2.79 (4) | 70.65 ± 0.66 (10) |
| AncA rv2+5 | 1.47 ± 0.05 (3) | 128.86 ± 10.59 (3) | 11.40 ± 1.00 (3) | 70.40 ± 0.99 (4) |
| AncA rv2+4+6 | 2.03 ± 0.01 (3) | 34.31 ± 0.71 (3) | 59.26 ± 1.27 (3) | 62.07 ± 0.66 (3) |

| | | | | |
|---|---|---|---|---|
| MLE | 1.44 ± 0.05 (3) | 107.64 ± 11.22 (3) | 13.42 ± 1.48 (3) | 50.06 ± 0.80 (4) |
| MLD | 2.36 ± 0.03 (3) | 56.93 ± 2.54 (3) | 41.51 ± 1.92 (3) | 55.05 ± 0.18 (2) |
| MLC | 0.68 ± 0.01 (3) | 35.99 ± 2.95 (3) | 18.80 ± 1.58 (3) | 61.75 ± 1.15 (4) |
| MLC rv2 | 1.01 ± 0.02 (3) | 35.64 ± 2.99 (3) | 28.24 ± 2.42 (3) | 62.03 ± 1.28 (4) |

**Table S2. List of primers used in this study.**

| Primer | Target | Sequence | Purpose |
|---|---|---|---|
| pRSFd BsaI_del for | pRSF-Duet1 | AAGCTGACGACCGAGTCTCCGCAAGTG | Domestication of internal BsaI site via site-directed mutagenesis |
| pRSFd BsaI_del rev | pRSF-Duet1 | CACTTGCGGAGACtCGGTCGTCAGCTT | Domestication of internal BsaI site via site-directed mutagenesis |
| pRSFd BsaIpcr for | pRSF-Duet1 | GGTCTCGCTAAAGCCAGGATCCGAATTCG | Linearization of domesticated pRSFDuet-1 with BsaI flanks |
| pRSFd BsaIpcr rev | pRSF-Duet1 | CTGAGGTCTCTAGAGGGGAATTGGATCC | Linearization of domesticated pRSFDuet-1 with BsaI flanks |
| rsf_A2_x10for | pRSF-Duet1 | TCTGAAGGGTCTCGCTAAAGCCA | Extension of linearized pRSF acceptor containing BsaI flanks |
| rsf_A2_x10rev | pRSF-Duet1 | TCTGCGAGGTCTCTAGAGGGGAATTG | Extension of linearized pRSF acceptor containing BsaI flanks |
| 5' Twist | Twist gene fragments | CAATCCGCCCTCACTACAACCG | Amplification of Twist gene fragments |
| 3' Twist | Twist gene fragments | TCCCTCATCGACGCCAGAGTAG | Amplification of Twist gene fragments |
| AmpRdom_Up | pET-21(+) | TGAGCGTGGGTCACGCGGTATCATT | Domestication of internal BsaI site via site-directed excision and ligation |
| AmpRdom_Down | pET-21(+) | AATGATACCGCGTGACCCACGCTCA | Domestication of internal BsaI site via site-directed excision and ligation |
| 3' 21-vecflanks | pET-21(+) | TACGTGATGGGTCTCGTTGCTGGCGCCTATATC | Linearization of domesticated pET-21(+) with BsaI flanks |
| 5' 21-vecflanks | pET-21(+) | TGACGTGCTGGTCTCGCTAACAAAGCCCGAAAGG | Linearization of domesticated pET-21(+) with BsaI flanks |
| cdf-A6A9_x10-rev | pCDFDuet-1 | TCTGACAGGTCTCTGGCAATTTCCTAATGC | Linearization of pCDFDuet-1 with BsaI flanks for both A6 and A9 configurations |
| cdf_EP_all2 | pCDFDuet-1 | AACGTCAGGTCTCTGGGAGTCTGGTAAAGAAACCGCTG | Linearization of pCDFDuet-1 with BsaI flanks for A6 configuration |
| cdf_EP_TTAC | pCDFDuet-1 | AACGTCAGGTCTCTTTACGTCTGGTAAAGAAACCGCTG | Linearization of pCDFDuet-1 with BsaI flanks for A9 configuration |
| De_L319M_Q5for | pRSFDuet-1 AncD RbcLSchis | TGTCAGGTGGGGACCA | Q5 mutagenesis |

| | | | |
|---|---|---|---|
| De_L319M_Q5Rev | pRSFDuet-1 AncD RbcLSchis | TACGGAGACACTTGGC | Q5 mutagenesis |
| De_M319L_Q5For | pRSFDuet-1 AncD RbcLSchis | TTGTCAGGTGGGGACCA | Q5 mutagenesis |
| De_M319L_Q5Rev | pRSFDuet-1 AncD RbcLSchis | ACGGAGACACTTGGCG | Q5 mutagenesis |
| De_M288L_Q5For | pRSFDuet-1 AncD RbcLSchis | TGCTTTTACACATACACAG | Q5 mutagenesis |
| De_M288L_Q5Rev | pRSFDuet-1 AncD RbcLSchis | ATCCATTGTCTCGACA | Q5 mutagenesis |
| De_S261T_Q5For | pRSFDuet-1 AncD RbcLSchis | CTCCCATCATAATGCACGA | Q5 mutagenesis |
| De_S261T_Q5Rev | pRSFDuet-1 AncD RbcLSchis | AACCGAGCTCCTTTGC | Q5 mutagenesis |
| De_W282Y_q5For | pRSFDuet-1 AncD RbcLSchis | TACTGTCGAGACAATGGA | Q5 mutagenesis |
| De_W282Y_q5Rev | pRSFDuet-1 AncD RbcLSchis | CTTTGCCAAAGAAGTATTTG | Q5 mutagenesis |
| De_C399V_q5For | pRSFDuet-1 AncD RbcLSchis | CGGAGTCGTCTCCGAAG | Q5 mutagenesis |
| De_C399V_q5Rev | pRSFDuet-1 AncD RbcLSchis | TTCTCCAATTCGGTGGTG | Q5 mutagenesis |
| Ce_Y283W_q5For | pRSFDuet-1 AncC RbcLSchis | GTGTCGAGACAATGGA | Q5 mutagenesis |
| Ce_Y283W_q5Rev | pRSFDuet-1 AncC RbcLSchis | CACTTTGCTAATGAAGTATTTG | Q5 mutagenesis |
| De_D351N_q5For | pRSFDuet-1 AncD RbcLSchis | ATACGTTGAGCAAGACCG | Q5 mutagenesis |
| De_D351N_q5Rev | pRSFDuet-1 AncD RbcLSchis | TCTCCCGCATTAAGTCTACG | Q5 mutagenesis |
| 100s_gibins_FOR | pRSFDuet-1 AncD f1 RbcLSchis | TGGTGAGGACAATCAATACATCTGTTACGT | Gibson cloning to transplant derived cluster 1 substitutions |
| 100s_gibins_REV | pRSFDuet-1 AncD f1 RbcLSchis | TTTTCAAGTAAGCAACTGGGATTCGGA | Gibson cloning to transplant derived cluster 1 substitutions |
| 100s_gibvec_FOR | pRSFDuet-1 AncD RbcLSchis variants | CCCAGTTGCTTACTTGAAAACTTTCCAAG | Gibson cloning to transplant derived cluster 1 substitutions |

| | | | |
|---|---|---|---|
| 100s_gibvec_REV | pRSFDuet-1 AncD RbcLSchis variants | TGTATTGATTGTCCTCACCAGGGACA | Gibson cloning to transplant derived cluster 1 substitutions |

**Table S3. List of plasmids used in this study.**

Vectors utilized for the cloning and expression of Rubisco, chaperonin, or assembly chaperone(s) cassettes. Entries marked with an asterisk denote generalized vector classes grouped by their functional module to represent all Golden Gate derived Rubisco, chaperonin, or assembly chaperone(s) combinations without enumerating each individual extant or ancestral variant.

| Plasmid | Notable features | Source |
|---|---|---|
| pRSFDuet-1 | *E. coli* expression vector, T7 promoter, Kan$^{R}$, *lacI*, lac operator, RBS, pRSF1030 replicon | Novagen |
| pET-21(+) | *E. coli* expression vector, T7 promoter, Amp$^{R}$, *lacI*, lac operator, RBS, pMB1 replicon | Twist Bioscience |
| pCDFDuet-1 | *E. coli* expression vector, T7 promoter, Spec/Strep$^{R}$, *lacI*, lac operator, RBS, pCloDF13 replicon | Novagen |
| pTwist Chlor High Copy | *E. coli* cloning vector, Chlor$^{R}$, pUC replicon | Twist Bioscience |
| pRSFDuet-1 acceptor | *E. coli* expression vector, T7 promoter, Kan$^{R}$, *lacI*, lac operator, RBS, pRSF1030 replicon, BsaI cloning flanks | This study |
| pCDFDuet-1 acceptor A9 | *E. coli* expression vector, T7 promoter, Spec/Strep$^{R}$, *lacI*, lac operator, RBS, pCloDF13 replicon, BsaI cloning flanks | This study |
| pCDFDuet-1 acceptor A6 | *E. coli* expression vector, T7 promoter, Spec/Strep$^{R}$, *lacI*, lac operator, RBS, pCloDF13 replicon, BsaI cloning flanks | This study |
| pET-21(+) acceptor A9L | *E. coli* expression vector, T7 promoter, Amp$^{R}$, *lacI*, mScarlet operon negative marker, lac operator, RBS, pMB1 replicon, PaqCI and BsaI cloning flanks | This study |
| pET-21(+) acceptor A6L | *E. coli* expression vector, T7 promoter, Amp$^{R}$, *lacI*, mScarlet operon negative marker, lac operator, RBS, pMB1 replicon, PaqCI and BsaI cloning flanks | This study |
| pTwist acceptor | *E. coli* cloning vector, Chlor$^{R}$, lacZ negative marker, pUC replicon, BbsI cloning flanks | This study |
| *pRSFDuet-1 RbcLS-cHis | *E. coli* expression vector, T7 promoter, Kan$^{R}$, *lacI*, lac operator, RBS, pRSF1030 replicon, containing respective RbcL and RbcS-cHis of interest | This study |
| *pCDFDuet-1 Assembly chaperones | *E. coli* expression vector, T7 promoter, Spec/Strep$^{R}$, *lacI*, lac operator, RBS, pCloDF13 replicon, containing respective assembly chaperone(s) of interest | This study |
| *pET-21(+) Chaperonins | *E. coli* expression vector, T7 promoter, Amp$^{R}$, *lacI*, lac operator, RBS, pMB1 replicon, containing respective GroL and GroS of interest | This study |

**Table S4. List of strains used in this study.**

| Strain | Genotype | Source or reference |
|---|---|---|
| *E. coli* NEB® 5-alpha | *fhuA2Δ(argF-lacZ)U169 phoA glnV44 Φ80Δ(lacZ)M15 gyrA96 recA1 relA1 endA1 thi-1 hsdR17* | New England Biolabs, #C2987 |
| *E. coli* BL21(DE3) | *fhuA2 [lon] ompT gal (λ DE3) [dcm] ∆hsdS; λ DE3 = λsBamHIo ∆EcoRI-B int::(lacI::PlacUV5::T7 gene1) i21 ∆nin5* | New England Biolabs, #C2527 |

### Materials Design Analysis Reporting (MDAR) Checklist for Authors

The MDAR framework establishes a minimum set of requirements in transparent reporting applicable to studies in the life sciences (see Statement of Task: doi:10.31222/osf.io/9sm4x.). The MDAR checklist is a tool for authors, editors, and others seeking to adopt the MDAR framework for transparent reporting in manuscripts and other outputs. Please refer to the MDAR Elaboration Document for additional context for the MDAR framework.

#### For all that apply, please note where in the manuscript the required information is provided.

#### Materials:

| Newly created materials | indicate where provided: page no/section/legend) | n/a |
|---|---|---|
| The manuscript includes a dedicated "materials availability statement" providing transparent disclosure about availability of newly created materials including details on how materials can be accessed and describing any restrictions on access. | In acknowledgement section | |

| Antibodies | indicate where provided: page no/section/legend) | n/a |
|---|---|---|
| For commercial reagents, provide supplier name, catalogue number and RRID, if available. | Described in Methods section under the subheading of Protein Biochemistry | |

| DNA and RNA sequences | indicate where provided: page no/section/legend) | n/a |
|---|---|---|
| **Short novel DNA or RNA including primers, probes:** Sequences should be included or deposited in a public repository. | Table S2 and S3 | |

| Cell materials | indicate where provided: page no/section/legend | n/a |
|---|---|---|
| **Cell lines:** Provide species information, strain. Provide accession number in repository **OR** supplier name, catalog number, clone number, **OR** RRID. | n/a | X |
| **Primary cultures:** Provide species, strain, sex of origin, genetic modification status. | n/a | X |

| Experimental animals | indicate where provided: page no/section/legend) | n/a |
|---|---|---|
| **Laboratory animals or Model organisms:** Provide species, strain, sex, age, genetic modification status. Provide accession number in repository **OR** supplier name, catalog number, clone number, **OR** RRID. | n/a | X |
| **Animal observed in or captured from the field:** Provide species, sex, and age where possible. | n/a | X |

| **Plants and microbes** | **indicate where provided: page no/section/legend)** | **n/a** |
|---|---|---|
| **Plants:** provide species and strain, ecotype and cultivar where relevant, unique accession number if available, and source (including location for collected wild specimens). | n/a | X |
| **Microbes:** provide species and strain, unique accession number if available, and source. | Table S5 | |

| **Human research participants** | **indicate where provided: page no/section/legend) or state if these demographics were not collected** | **n/a** |
|---|---|---|
| If collected and within the bounds of privacy constraints report on age, sex and gender or ethnicity for all study participants. | n/a | X |

#### Design:

| **Study protocol** | **indicate where provided: page no/section/legend)** | **n/a** |
|---|---|---|
| If study protocol has been pre-registered, provide DOI. For clinical trials, provide the trial registration number **OR** cite DOI. | n/a | X |

| **Laboratory protocol** | **indicate where provided: page no/section/legend)** | **n/a** |
|---|---|---|
| Provide DOI **OR** other citation details if detailed step-by-step protocols are available. | n/a | X |

| **Experimental study design (statistics details)** | | |
|---|---|---|
| **For in vivo studies:** State whether and how the following have been done | **indicate where provided: page no/section/legend. If it could have been done, but was not, write not done** | **n/a** |
| Sample size determination | n/a (in-vitro study) | X |
| Randomisation | n/a (in-vitro study) | X |
| Blinding | n/a (in-vitro study) | X |

| | | |
|---|---|---|
| Inclusion/exclusion criteria | n/a (in-vitro study) | X |

| Sample definition and in-laboratory replication | indicate where provided: page no/section/legend | n/a |
|---|---|---|
| State number of times the experiment was replicated in laboratory. | Number of replicates is stated in the associated legend description of figure and table where data is shown | |
| Define whether data describe technical or biological replicates. | Data type is defined in the associated legend description of figure and table where data is shown | |

| Ethics | indicate where provided: page no/section/legend | n/a |
|---|---|---|
| **Studies involving human participants:** State details of authority granting ethics approval (IRB or equivalent committee(s), provide reference number for approval. | Study does not involve human participants. | X |
| **Studies involving experimental animals:** State details of authority granting ethics approval (IRB or equivalent committee(s), provide reference number for approval. | Study does not involve human participants. | X |
| **Studies involving specimen and field samples:** State if relevant permits obtained, provide details of authority approving study; if none were required, explain why. | Study does not involve human participants. | X |

| Dual Use Research of Concern (DURC) | indicate where provided: page no/section/legend | n/a |
|---|---|---|
| If study is subject to dual use research of concern regulations, state the authority granting approval and reference number for the regulatory approval. | Study is not subject to dual use research of concern regulations | X |

#### Analysis:

| Attrition | indicate where provided: page no/section/legend | n/a |
|---|---|---|
| Describe whether exclusion criteria were preestablished. Report if sample or data points were omitted from analysis. If yes report if this was due to attrition or intentional exclusion and provide justification. | No data was excluded from the analysis. | |

| Statistics | indicate where provided: page no/section/legend | n/a |
|---|---|---|
| Describe statistical tests used and justify choice of tests. | Head-to-head comparisons between wild type and mutant conditions were evaluated using one-way ANOVA followed by multiplicity-adjusted post hoc tests to control familywise Type I error when comparing more than two groups, rather than | |

| | |
|---|---|
| | performing multiple uncorrected t-tests. For pairwise localization of differences, Tukey's HSD was used when all pairwise contrasts were of interest. We treated group variances as approximately equal because assembly-yield measurements were generated under standardized assay conditions and exhibited similar dispersion across groups (absolute values and within-group variability generally differed by less than ~3-fold), a regime under which the pooled-error ANOVA has been typically shown to be robust. |

| Data availability | indicate where provided: page no/section/legend | n/a |
|---|---|---|
| For newly created and reused datasets, the manuscript includes a data availability statement that provides details for access or notes restrictions on access. | Rawdata is uploaded to Edmond, the data repository of<br><br>the Max Planck Society and publicly available. | |
| If newly created datasets are publicly available, provide accession number in repository **OR** DOI **OR** URL and licensing details where available. | The DOI is cited in the acknowledgements. | |
| If reused data is publicly available provide accession number in repository **OR** DOI **OR** URL, **OR** citation. | n/a | X |

| Code availability | indicate where provided: page no/section/legend | n/a |
|---|---|---|
| For all newly generated custom computer code/software/mathematical algorithm or re-used code essential for replicating the main findings of the study, the manuscript includes a data availability statement that provides details for access or notes restrictions. | No custom computer code was generated. Scripts and publicaly available programs required to replicate the main findings of the study are cited in the methods section and are available from the listed citations. | X |
| If newly generated code is publicly available, provide accession number in repository, **OR** DOI **OR** URL and licensing details where available. State any restrictions on code availability or accessibility. | No code was newly generated herein | X |
| If reused code is publicly available provide accession number in repository **OR** DOI **OR** URL, **OR** citation. | n/a | X |

#### Reporting

MDAR framework recommends adoption of discipline-specific guidelines, established and endorsed through community initiatives. Journals have their own policy about requiring specific guidelines and recommendations to complement MDAR.

| Adherence to community standards | indicate where provided: page no/section/legend | n/a |
|---|---|---|
| State if relevant guidelines (e.g., ICMJE, MIBBI, ARRIVE) have been followed, and whether a | n/a | X |

| |
|---|
| checklist (e.g., CONSORT, PRISMA, ARRIVE) is provided with the manuscript. |
